# A unifying comparative phylogenetic framework including traits coevolving across interacting lineages

**DOI:** 10.1101/069518

**Authors:** Marc Manceau, Amaury Lambert, Hélène Morlon

## Abstract

Models of phenotypic evolution fit to phylogenetic comparative data are widely used to make inferences regarding the tempo and mode of trait evolution. A wide range of models is already available for this type of analysis, and the field is still under active development. One of the most needed developments concerns models that better account for the effect of within- and between-clade interspecific interactions on trait evolution, that can result from processes as diverse as competition, predation, parasitism, or mutualism. Here, we begin by developing a very general comparative phylogenetic framework for (multi)-trait evolution that can be applied to both ultrametric and non-ultrametric trees. This framework not only encapsulates all previous classical models of univariate and multivariate phenotypic evolution, but also paves the way for the consideration of a much broader series of models in which lineages co-evolve, meaning that trait changes in one lineage are influenced by the value of traits in other, interacting lineages. Next, we provide a standard way for deriving the probabilistic distribution of traits at tip branches under our framework. We show that a multivariate normal distribution remains the expected distribution for a broad class of models accounting for interspecific interactions. Our derivations allow us to fit various models efficiently, and in particular greatly reduce the computation time needed to fit the recently proposed phenotype matching model. Finally, we illustrate the utility of our framework by developing a toy model for mutualistic coevolution. Our framework should foster a new era in the study of coevolution from comparative data.

---

Evolutionary biologists have long been interested in the long-term evolution of phenotypic traits (Simpson 1944). In 1973, Felsenstein introduced one of the first models of phenotypic evolution, with the initial goal to account for shared ancestry when testing for statistical correlation between pairs of traits in extant species. In this founding paper, Felsenstein proposed that a one-dimensional quantitative trait evolving on a tree could be modeled as a Brownian process that splits into two independent Brownian processes at branching times. This model mimics a trait that would evolve as a mere effect of stochastic drift; it is now often used as a null model, but also to estimate the relative lability (or rate of evolution) of various traits in a given group of organisms or of a given trait across different groups of organisms (Thomas et al. 2006; Harmon et al. 2010).

Since these early developments, evolutionary biologists have designed a series of models to better understand the evolutionary processes that shape phenotypic evolution (see Pennell and Harmon 2013 for a review). The Ornstein-Uhlenbeck (OU) process has been proposed to model evolution under stabilizing selection, i.e. with a selective pressure pushing trait values toward a given optimum (Hansen 1997; Hansen and Martins 1996; Butler and King 2004). The ACDC model has been proposed to account for accelerating (AC) or decelerating (DC) rates of phenotypic evolution through time (Blomberg et al. 2003). The latter scenario, where the evolutionary rate is high early in the history of a clade and subsequently declines toward the present, well known as the early burst (EB) model, has often been used to test support for adaptive radiation theory (Harmon et al. 2010; Moen and Morlon 2014). These univariate models representing the evolution of a single trait have been extended to multivariate models representing the simultaneous evolution of multiple traits, which permit investigators to directly test hypotheses about the coevolution between several phenotypic traits (Hansen et al. 2008; Bartoszek et al. 2012; Jhwueng and Maroulas 2014). Other extensions have been developed to account for variations in model parameters across clades (O'Meara et al. 2006; Revell and Collar 2009; Eastman et al. 2011; Butler and King 2004; Beaulieu et al. 2012). Finally, some of these models have been developed in the context of phylogenies including fossil data (i.e. non ultrametric trees, see Ruta et al. 2006; Slater 2015) in addition to phylogenies with only extant taxa (i.e. ultrametric trees). Most of these models have been implemented in open-access packages (Martins 2004; Harmon et al. 2008; Butler and King 2004; Thomas and Freckleton 2012; Clavel et al. 2015; Morlon et al. 2015), allowing their application to a broad variety of questions and datasets (see, e.g. Labra et al. 2009; Mahler et al. 2010; Dale et al. 2015; Quintero et al. 2015; Slater 2015).

Despite these developments, most currently available models ignore the effect of interspecific interactions on trait evolution. Given the importance of species interactions in classical evolutionary theories, such as Simpson's adaptive radiation (Simpson 1944), Ehlrich & Raven's escape and radiate (Ehrlich and Raven 1964) and Van Valen's Red Queen (Van Valen 1973) theories, building models that better account for such interactions is fundamental. In a first attempt to take into account the role of competition for niche space on character evolution, a diversity-dependent (DD) model has been introduced, where the rate of phenotypic evolution declines as the number of lineages in the clade increases (Mahler et al. 2010; Weir and Mursleen 2013). While this model represents an important first step, it still assumes that trait changes in one lineage are independent from the value of traits in other, interacting lineages, therefore ignoring the widespread idea of trait- (or ecologically-) driven interspecific interactions. More recently, the phenotype matching (PM) model relaxed these hypotheses and more explicitly accounted for interspecific interactions by modeling either the attraction or the repulsion of traits from a clade-wise average trait value (Nuismer and Harmon 2014; Drury et al. 2016). In the first case, referred to as matching mutualism, species traits tend to converge to similar values, whereas in the second case, referred to as matching competition, species traits tend to diverge.

The comparative phylogenetic approach developed by Drury et al. (2016) is one of the first that allows fitting a model where the evolution of trait values in one lineage is influenced by the trait values of other lineages. This approach focused on the evolution of traits within one clade. While within-clade interactions can be particularly relevant for some types of interactions (e.g. in the case of competitively driven character displacement, Brown and Wilson 1956), the effect of other types of antagonistic or mutualistic interactions on trait evolution is often most relevant between distantly related species. For example, host-parasite interactions are thought to drive a coevolutionary race between traits involved in host defence and parasite ability to infect. Similarly, prey-predator interactions may lead to the coevolution of prey traits involved in camouflage, repulsion, or escape strategies, together with predator traits involved in the ability to detect and capture its prey (Ehrlich and Raven 1964; Dawkins and Krebs 1979). Mutualistic plant-pollinator interactions also are thought to drive the coevolution between plant traits involved in pollen accessibility or flower attractiveness to their pollinator (secondary metabolites, floral traits), and pollinator traits involved in the ability to detect suitable plants and to exploit plant rewards (Fenster et al. 2004; Weiblen 2004; Sletvold et al. 2016). While these types of biotic interactions likely play a key role in trait evolution and have been crucial in the development of coevolutionary theories (Ehrlich and Raven 1964; Van Valen 1973), there currently exists no framework for fitting models of phenotypic evolution incorporating the effect of clade-clade interactions.

The current paper expands the work of Bartoszek et al. (2012) who presented a unified framework for studying coevolving traits in independently evolving lineages by providing a unified framework for coevolving traits in coevolving lineages. Our framework, based on linear stochastic differential equations (SDE), encompasses all models of continuous (multi-)trait evolution mentioned above, and allows the treatment of a much broader set of models. The goal of the paper is two-fold: first, by providing general solutions to the distribution of traits at tip branches under our unified framework, we hope to help users to find their way in a dense and potentially overwhelming literature; second, by showing how the framework can be used to treat a broad class of within-clade and clade-clade coevolutionary scenarios, we hope to foster the development of models to test long-standing hypotheses on the role of competition, predation, parasitism and mutualism in evolution.

We begin by presenting our framework and showing how previous models as well as novel clade-clade coevolutionary models fit within this framework; next, we provide general solutions for the distribution of tip trait values under this framework; then, we illustrate how the framework can be used to study a toy model of clade-clade coevolution.

## A general framework for phenotypic evolution

### Notation for trees and traits

We introduce a general formalism to study (multi-)trait (co)evolution when the interaction between lineages within a clade or among several clades potentially affects how traits evolve. We consider a single or several clades represented by a single or several fixed, binary, time-calibrated phylogenetic trees (non-necessarily ultrametric, i.e. the trees can include fossils). Time *t* runs from the root of the oldest tree (*t* = *τ*_0_ = 0) to the most recent tip of all trees (*t* = *T* is the present if at least one of the phylogenies includes extant species). The *K* successive branching and extinction times when considering the various trees altogether are denoted by 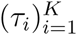 and the time-intervals between two such events are called epochs, following Butler and King (2004). We denote by *n_t_* the total number of lineages that arose before (and at) time *t*.

In the case of trait evolution within a single clade (Fig. 1), we assign numbers (from 1 to *n_t_*) to lineages by order of arrival. At each branching event *τ*, one daughter lineage inherits the number assigned to the ancestral lineage while the other one is assigned *n_τ_*.

**Figure 1:**
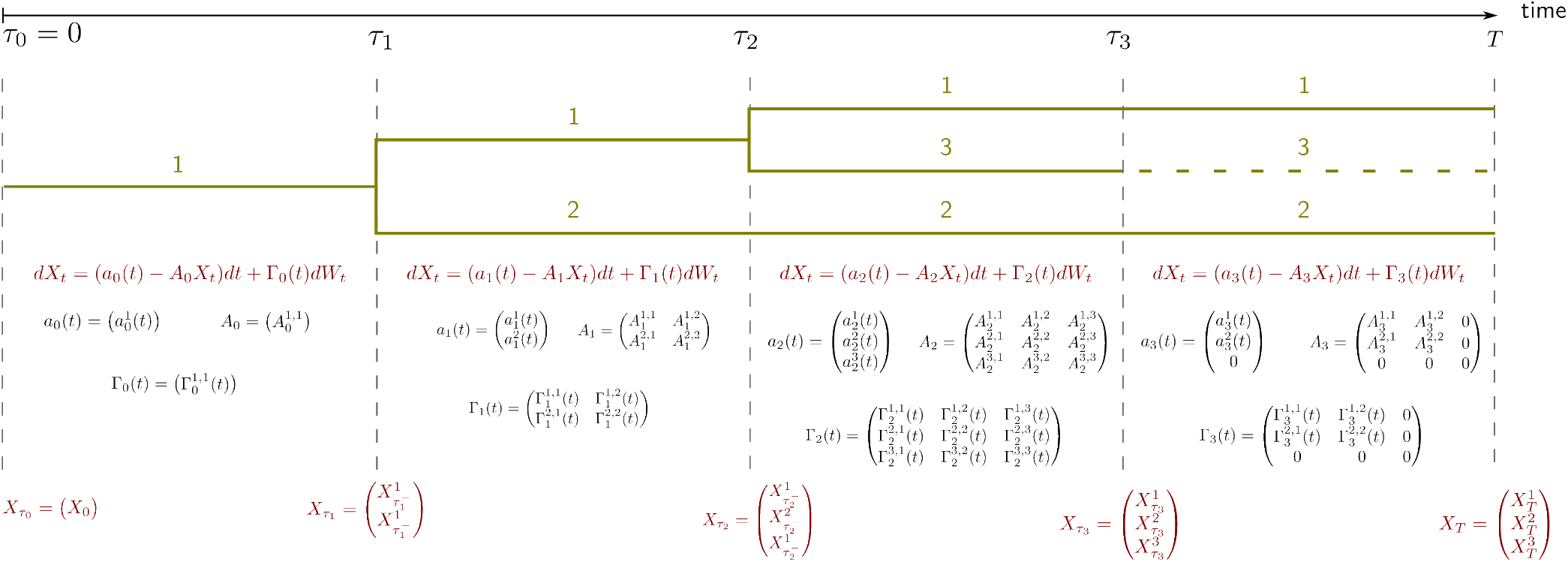
Formalism used throughout the paper, to model the evolution of one trait on a non-ultrametric tree. Epochs are separated with vertical dashed lines.

We model the evolution of *d* one-dimensional quantitative traits. We denote by 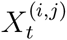 the value of trait *j* (1 ≤ *j* ≤ *d*) on branch *i* at time *t* and *X_t_* the column vector containing the values of all traits on all lineages at time *t*, ordered as follows:
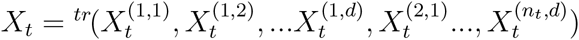, where ^tr^ stands for the transposition.

In the case of trait evolution in *c* distinct (co)evolving clades, we begin by arbitrarily ordering the clades from 1 to *c*; then, we assign numbers to lineages following the formalism introduced above, first numbering lineages from clade 1, then clade 2, and so on. As above, we denote by *X_t_* the column vector containing the values of all traits on all lineages at time *t*, which now is a concatenation of the *c* column vectors corresponding to each clade.

### Trait evolution through time

Given one (or several) phylogenetic tree(s), a model of phenotypic evolution is entirely defined by initial conditions *X*_0_ on the trait values at the root(s) and a set of rules dictating how the vector of traits *X_t_* is updated (i) at branching times, (ii) on each epoch (i.e. between two branching or extinction times), and (iii) after a death time. These rules are illustrated in Figure 1 for a single trait evolving on a single small tree.

In line with most models of phenotypic evolution, we consider anagenetic character evolution, meaning that traits do not change at cladogenesis. Hence, at a given branching time *τ*, each of the daughter lineages inherits the trait value of their mother lineage. Our framework can easily be modified to treat non binary trees including polytomies. In practice, in the case of evolution within a single clade, the new vector *X_τ_* is obtained by concatenating the *d* trait values of the branching lineage at time *τ* at the end of *X*_*τ* −_ (where *τ* − is the time just preceding the branching event). In the case of evolution in several clades, the new vector *X_τ_* is obtained by inserting the *d* trait values of the branching lineage at time *τ* at the appropriate location in *X_τ −_* (i.e., at the end of the part of *X_τ −_* corresponding to the clade in which the branching event is occuring).

On each given epoch (*τ_i_*, *τ*_*i*+1_) (*i* ∈ {0, 1,…, *K* − 1}), we assume that the evolution of the *d* traits on the *n* lineages is driven by a linear stochastic differential equation of the form:

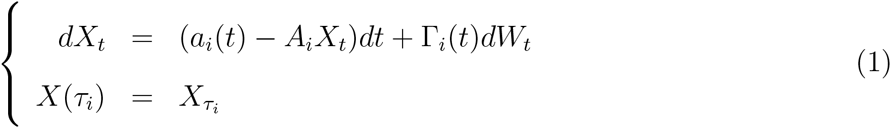

where *a_i_* is a vector of ℝ*^nd^* whose coefficients can vary with time, *A_i_* is a constant square matrix of size ℝ*^nd^* × ℝ*^nd^*, Γ*_i_* is a square matrix of the same size whose coefficients can vary with time, and *W_t_* is a *nd*-Brownian motion (i.e. a vector composed of *nd* independent standard Brownian motions). Intuitively, the deterministic part (*a_i_*(*t*) − *A_i_X_t_*)*dt* reflects the direct effects of trait values for species in the clade(s) at time *t* on the evolution of these traits, including the effect of a trait value in one lineage on both its own evolution (as in the OU process) and the evolution of traits in the other lineages (as in the PM process); the stochastic part Γ*_i_*(*t*)*dW_t_* reflects drift and the environmental noise influencing trait evolution. It has been proposed that correlations within the covariance matrix Γ*_i_* represent non-causal correlations, for example linked to joint evolutionary responses to shared environmental conditions, while correlations within the interaction matrix *A_i_* represent causal effects (Reitan et al. 2012; Liow et al. 2015). More work is needed to assess the relevance of the ‘causal/non-causal’ dichotomy, and the difference of patterns it can yield in the context of trait evolution on phylogenies. For simplicity, in the present paper, we will stick to the term ‘correlations’, and consider only models making the simplifying assumption that Γ*_i_* is diagonal, but the framework could be equally adapted to incorporate correlations through these Γ*_i_* matrices.

Finally, when a lineage goes extinct at a given time *τ*, its *d* trait values no longer evolve (i.e. they are frozen at the extinction time), and they no longer have any influence on the evolution of the traits of other lineages until reaching the end of the process at time *t* = *T*. In practice, this means that the vector *X_τ_* is simply equal to *X_τ−_*, and that the *d* lines and columns in *a_i_*, *A_i_* and Γ*_i_* corresponding to the now extinct lineage are all set to zero.

We will show later that this general formulation encapsulates all classical models of phenotypic evolution, ensures analytical tractability, and further allows the incorporation of a broad set of interspecific coevolutionary scenarios.

Given the above, initial conditions on *X*_0_, and the collection of (*a_i_*), (*A_i_*) and (Γ*_i_*) on each epoch fully define a process of trait evolution on one or several trees.

All models written under the formalism that we propose can easily be simulated numerically. First, the whole trajectory of the process can be simulated using a numerical scheme for SDE such as the Euler-Maruyama scheme (Gardiner et al. 1985) on each epoch, and augmenting the vector of traits at branching times with traits corresponding to the branching lineage (see Appendix D.1 and Fig. 5). Second, we show in the next section how to compute numerically the tip distribution. Tip values can then directly be drawn in a fast way from the tip distribution.

### Application: existing and novel models of trait evolution

We first show that the general formulation above encapsulates all classical models of phenotypic evolution, before showing how it further allows considering a much broader set of models, including models of within and between clades coevolution.

Models of phenotypic evolution have traditionally been characterized by a stochastic differential equation specifying how a given trait evolves along a single lineage. Applying Equation (1) to trait *k*, on epoch *i* yields:

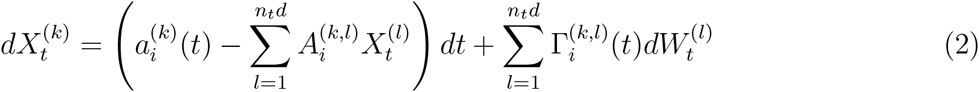

where the two sums are taken over all traits and all lineages. The term 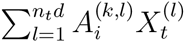 is the term that specifies how the value of trait *k* and all other traits in all other lineages influence the evolution of trait *k*. Given a well-known differential equation specifying how a given trait evolves along a single lineage for a previously proposed model of phenotypic evolution (second column in Table 1), deriving the corresponding expressions for *a*, *A* and Γ using Equation (2) is straightforward. Table 1 summarizes these expressions for existing univariate models running on ultrametric trees.

**Table 1:**
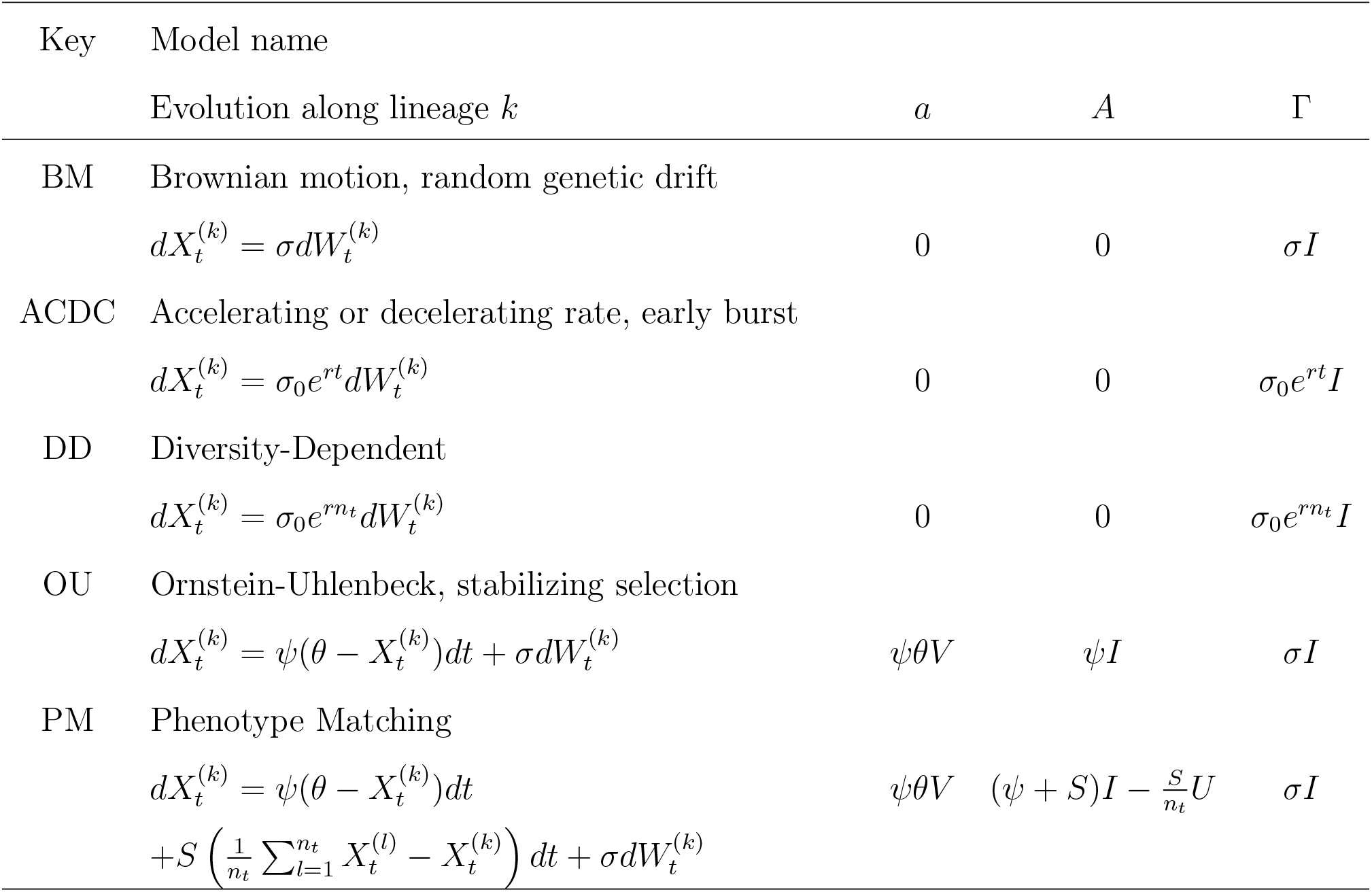
Expression of *a*, *A* and Γ for models of trait evolution that have been proposed in the literature. The unity vector (vector full of 1) is denoted by *V*, *I* refers to the identity matrix (diagonal matrix with diagonal values equal to 1), and *U* refers to the unity matrix (matrix full of 1). Their size is the same as the size of the vector of traits *X_t_* considered. Parameters are *σ*: rate of neutral phenotypic evolution; *ψ*: strength of stabilizing selection; *θ*: optimal phenotype; *S*: strength of between-lineage competition driving individual phenotypes away from clade-wise average phenotype; *σ*_0_: rate of phenotypic evolution at the root of the tree; *r*: parameter controling the exponential rise or decay of the rate of phenotypic evolution with time (ACDC) or with the number of lineages (DD).

| Key | Model name |  |  |  |
| --- | --- | --- | --- | --- |
| | Evolution along lineage $k$ | $a$ | $A$ | $\Gamma$ |
| BM | Brownian motion, random genetic drift<br>$dX_t^{(k)} = \sigma dW_t^{(k)}$ | 0 | 0 | $\sigma I$ |
| ACDC | Accelerating or decelerating rate, early burst<br>$dX_t^{(k)} = \sigma_0 e^{rt} dW_t^{(k)}$ | 0 | 0 | $\sigma_0 e^{rt} I$ |
| DD | Diversity-Dependent<br>$dX_t^{(k)} = \sigma_0 e^{rn_t} dW_t^{(k)}$ | 0 | 0 | $\sigma_0 e^{rn_t} I$ |
| OU | Ornstein-Uhlenbeck, stabilizing selection<br>$dX_t^{(k)} = \psi(\theta - X_t^{(k)})dt + \sigma dW_t^{(k)}$ | $\psi\theta V$ | $\psi I$ | $\sigma I$ |
| PM | Phenotype Matching<br>$dX_t^{(k)} = \psi(\theta - X_t^{(k)})dt$<br>$+ S \left( \frac{1}{n_t} \sum_{l=1}^{n_t} X_t^{(l)} - X_t^{(k)} \right) dt + \sigma dW_t^{(k)}$ | $\psi\theta V$ | $(\psi + S)I - \frac{S}{n_t}U$ | $\sigma I$ |

The first three models (BM, ACDC and DD) are models in which trait evolution along a lineage is influenced neither by the trait value of this lineage nor the trait value of any other lineage. The corresponding *A* matrices are null matrices, as would be the case for any model with the latter property. The fourth model (OU) is a model in which trait evolution along a lineage is influenced by its own trait value, but not the trait values of other lineages. The corresponding *A* matrix is diagonal, as would be the case for any model with this property. Finally, the last model (PM) is a model in which trait evolution along a lineage is influenced by its own trait value and the trait values of other lineages, such that *A* has non-negative off-diagonal values. A remarkable property of *A* under this model is that all its off-diagonal values are identical. This is explained by the fact that the PM model is a neutral model, in the sense that the effect *A*^(*k,l*)^ of lineage *l* on lineage *k* is the same for all lineages *k* ≠ *l*. All other models in which the off-diagonal elements of *A* are identical would have this same property, known in probability theory as exchangeability.

Several variations around these models can still be embedded in our general framework: i) Models in which the rate of phenotypic evolution depends on a variable *Y* (*t*) that itself varies through time (see, e.g. global temperature *T* (*t*) in Clavel and Morlon 2016) can be formalised similarly to ACDC, with time *t* replaced by *Y* (*t*). ii) Models accounting for the biogeographic background in which species coevolved (e.g. all the “+GEO” models in Drury et al. 2016) can be incorporated in our framework through the design of the *A* matrix (see details in Appendix C.3). iii) Considering non-ultrametric trees including fossils amounts to replacing vector *V* and matrices *I* and *U* by their homologs *V*_alive_, *I*_alive_ and *U*_alive_, where the subscript specifies that the vector and matrices have 0 on lines and columns corresponding to lineages that are extinct in the given epoch. iv) Considering subclades in which trait evolution follows distinct modes or similar modes with distinct parameter values (as in Butler and King 2004) is also straightforward. One just needs to specify distinct parameters in *a*, *A* and Γ on the lines and columns corresponding to lineages in the distinctive subclade. v) Multivariate trait evolution models, in which several distinct traits evolve in a correlated manner (Hansen et al. 2008; Bartoszek et al. 2012) are easily written in our framework, as shown with some examples in Appendix B.2. In multivariate models with lineages evolving independently from one another (e.g. multivariate combinations of BM, ACDC, DD and OU models), *A* and Γ are block diagonal matrices, with blocks of size the number of traits, each of them describing correlated multivariate evolution along a particular lineage. In this case, trait-trait correlations introduced through the *A* matrix correspond, as in Bartoszek et al. (2012), to the case when a given trait on a lineage is attracted to (or repulsed from) a linear combination of other traits in this lineage.

By considering previous models under this light, it becomes very clear that the set of models that have been considered so far represents a very small fraction of all the models that could potentially be considered. In particular the *A* matrix, which dictates how the value of a given trait influences the evolution of other traits – either different traits in the same lineage, or the same trait in other lineages, or yet different traits in other lineages – has so far been very constrained. It has been considered to be zero (BM, ACDC, DD), diagonal (OU), block diagonal (multivariate), and only recently with non-zero off-diagonal values (PM). Relaxing these constraints means that a much broader array of models incorporating the effect of interspecific interactions on phenotypic evolution can be considered. In particular, lineages do not need to be interchangeable. Evolution in complex networks of interactions can be considered by designing a priori the *A* matrix according to the known network. The effect of clade-clade interactions can be modeled by filling the *A* matrix with non-zero entries *A*^(*k,l*)^ with *k* and *l* corresponding to lineages from different clades. For example, under a scenario of two clades coevolving with no effect of within-clade interactions, this leads to a *A* matrix with two off-diagonal blocks.

We can thus imagine a variety of coevolutionary scenarios, the only major constraint being that the effect of a trait value on the evolution of other traits is assumed to be linear (Equations (1) & (2)). Given a scenario, we can write the corresponding evolution of each trait on a given lineage on each epoch (Equation (2)), and deduce the collection of (*a_i_*), (*A_i_*) and (Γ*_i_*) defining the evolutionary process (Equation (1)). Below, we first show how to derive the probabilistic distribution of traits at tip branches for any model that can be written under this framework before illustrating the approach with a particular model of clade-clade interaction.

## Distribution of tip trait values

### The distribution of traits is Gaussian

Deriving the probabilistic distribution of traits at tip branches is key to our ability to fit phenotypic models to comparative data using maximum likelihood or Bayesian approaches. It also provides a very efficient way to simulate tip values for specific models, by drawing from the expected tip distribution.

We show (Appendix A.1) that if *X*_0_ has a Gaussian distribution (including the particular case when *X*_0_ is constant) and *X_t_* evolves according to our general framework, then *X_t_* remains a Gaussian vector at each time *t*. The trait vector *X_t_*, of size *n_t_d*, is thus uniquely defined by its expectation vector *m_t_* and covariance matrix Σ*_t_*, and has the following density:

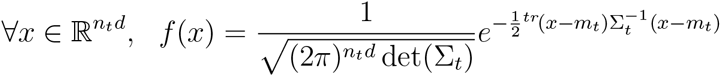

In particular, the distribution of tip trait values at present time *T* is Gaussian with expectation vector *m_T_* and covariance matrix Σ*_T_*. We can compute *m_T_* and Σ*_T_* iteratively: starting with initial conditions *m*_0_ and Σ_0_ for *X*_*τ*0_ = *X*_0_, we compute, until reaching the present:

1. 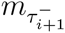 and 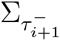 at the end of each epoch *i*
2. 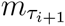 and 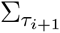 at the branching time *τ*_*i*+1_

### Evolution of the distribution on each epoch

Knowing the expectation vector and covariance matrix 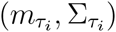 at the beginning of epoch *i*, we show (Appendix A.2) that 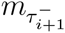 and 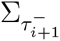 at the end of epoch *i* are given by the following analytical expressions:

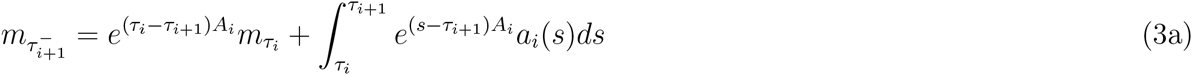

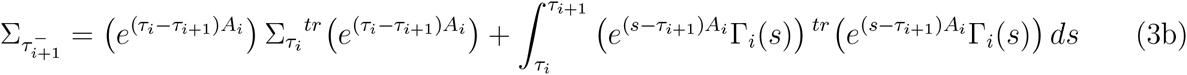

Alternatively, we can write the evolution of *m* and Σ on epoch *i* as a set of ordinary differential equations (ODE), and integrate these ODEs numerically, with initial conditions given by 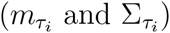. Each component *k* (resp. (*k, l*)) of the expectation vector (resp. Covariance matrix) evolves on epoch *i* according to (Appendix A.3):

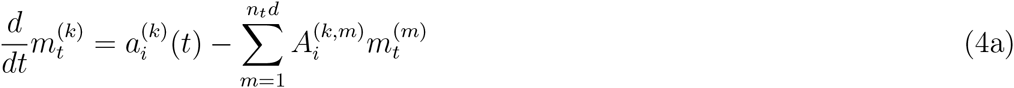

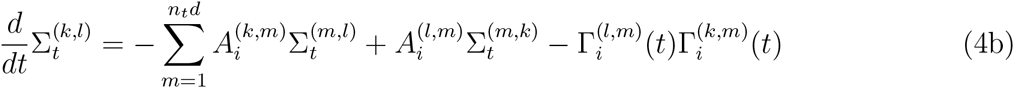

Equations (3a, 3b) and the ODE system described by Equations (4a, 4b) are mathematically equivalent. The first formulation is more computationally efficient when the integrals can be simplified analytically. For example when *A* is symmetric Equations (3a, 3b) can be simplified (Appendix C.1) and computed very efficiently. The second one provides a more intuitive interpretation of the components that influence the evolution of trait distribution, and is easily implementable for any model.

### Evolution of the distribution at branching times

Knowing the expectation vector and covariance matrix 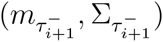 at the end of epoch *i*, which precedes the branching of a given lineage *j*, we build 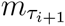 and 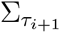 at the branching event, as illustrated in Figure 2.

**Figure 2:**
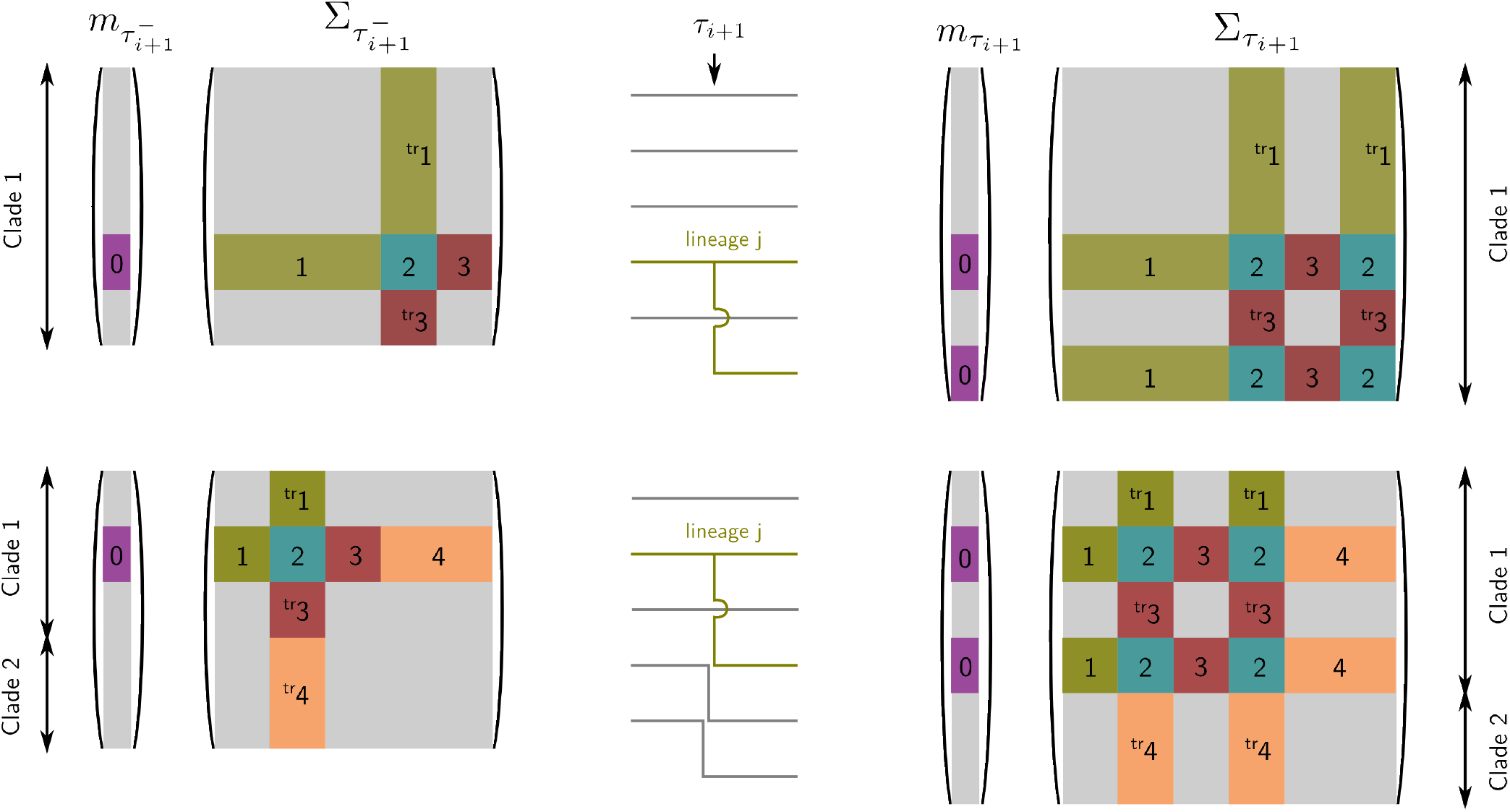
Toy example illustrating how to build the expectation vector and covariance matrix at branching times when there is one (top row) or two (bottom row) clades. Lineage *j* branches at time *τ_i_*_+1_ (middle). The vector 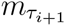 and matrix 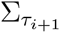 (on the right) are constructed by augmenting 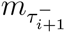 and 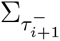 (on the left) with copies of blocks corresponding to lineage *j* (materialized by colors and numbers).

Recall that in the case of evolution within a single clade, 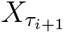 is obtained by concatenating the *d* trait values of lineage *j* at time 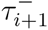 at the end of 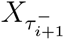. The *d* new components in 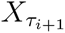 are thus the exact copies of the trait values of lineage *j*, and have the same expectation and covariance matrix. Hence, the expectation vector 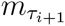 is simply obtained by concatenating 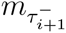 with the *d* components of 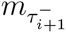 corresponding to lineage *j*. The covariance matrix 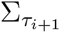 is obtained as follows: the covariance matrix corresponding to the previously existing lineages is unchanged, given by 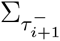; to this main block, we add below (and to the right) a copy of the *d* lines (respectively the *d* columns) corresponding to the covariances between the *d* traits in lineage *j* and all the other traits; finally, we fill the last missing block in the bottom right corner of 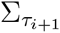 with the block corresponding to the covariance matrix among the *d* traits in lineage *j* (i.e. the *d* × *d* diagonal block of 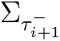 starting from line (*j* − 1)*d* + 1).

In the case of evolution in multiple clades, 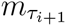 and 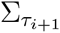 are constructed following a similar procedure, by augmenting 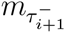 and 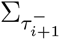 with copies of blocks corresponding to lineage *j*, inserted at the appropriate location. We illustrate this update step in Figure 2.

### Tip trait distribution for particular models

Applying this general iterative procedure along a phylogenetic tree provides closed analytic tip distribution formulae for a wide variety of models. In Appendix B, we re-derive known tip distributions for models without lineage-lineage interaction, thus providing a unified review of mathematical results associated to these models. Tip distributions for classical univariate models (BM, ACDC, DD, OU) on ultrametric and non-ultrametric trees are summarized in Table S1. We confirm, as has been shown before (Uyeda et al. 2015), that the OU and AC models have identical tip distributions on ultrametric trees. We also re-derive results that can be found in Bartoszek et al. (2012) providing tip distributions for multivariate models.

Analytical formulae of tip distributions for models with lineage-lineage interactions have not yet been proposed. Drury et al. (2016) developed the inference tools that allow fitting the PM model, using the ODE system given in Equations (4a, 4b) (thereafter referred to as ‘ode’ method). Here, we develop the inference tools based on analytical reduction of Equations (3a, 3b), (thereafter referred to as ‘analytical’ method, see Appendix C.2), and compare the efficiencies of the two methods. Specifically, we simulated 10 pure-birth Yule trees with a per-lineage speciation rate of 1 per time unit, conditioned to having a given number of tips at present, using the ‘phytools’ R package (Revell 2012). We then computed the tip distribution corresponding to the PM model with parameters fixed at (*m*_0_, *v*_0_, *θ, ψ, S, σ*) = (0, 0, 1, 0.1, 1, 2) using both the analytical and the ode methods. The new analytical method is much more efficient than the previous ode method (Fig. 3). While we were previously limited to fitting the PM model to trees of less than 150 tips due to memory issues, the analytical methods allows fitting trees with up to 600 tips on a desktop computer.

**Figure 3:**
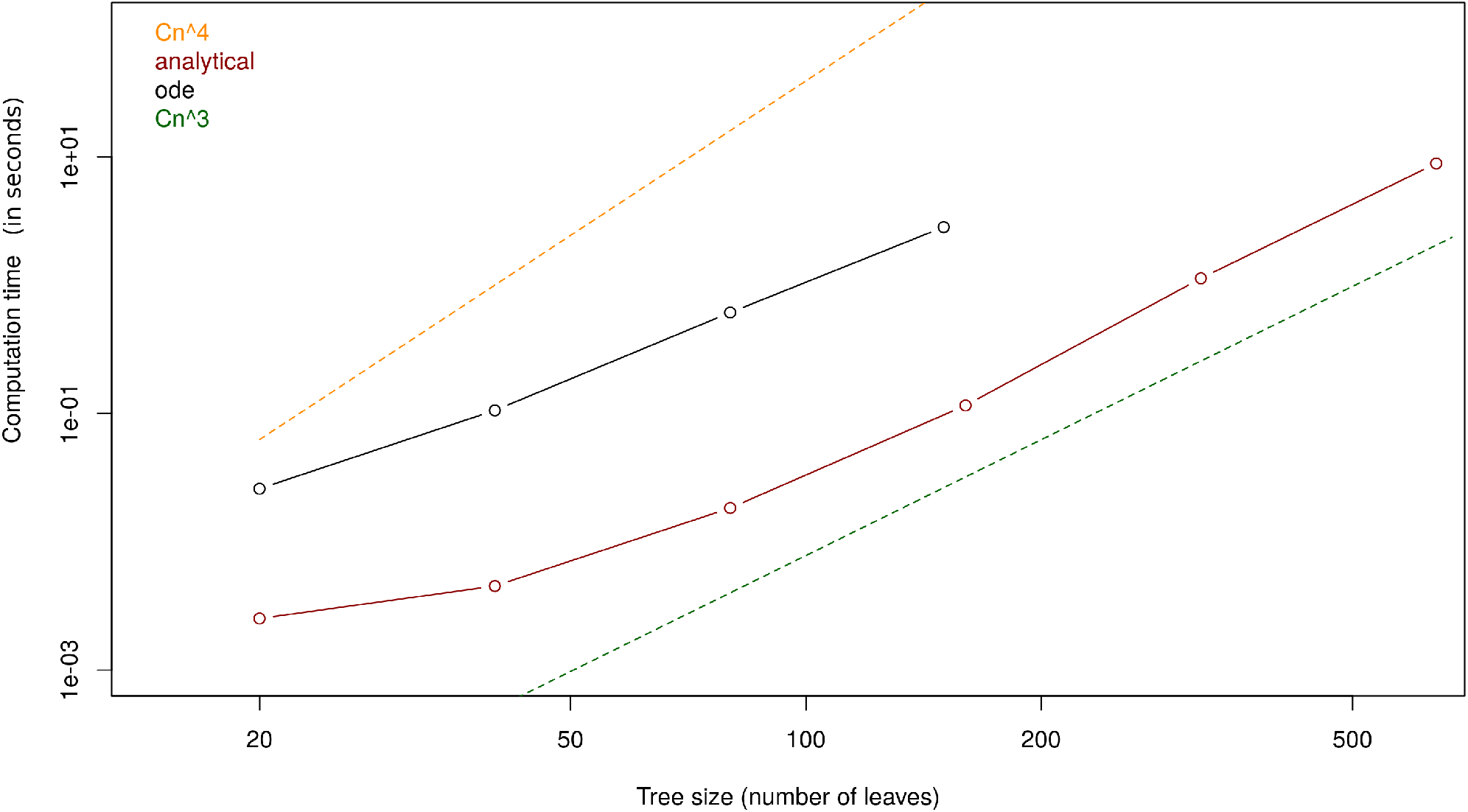
Time needed to compute the distribution of tip data following the PM model with parameters (*m*_0_, *v*_0_, *θ, ψ, S, σ*) = (0, 0, 1, 0.1, 1, 2). Trees are simulated under a pure-birth model conditioned on having a given number of leaves. Red curve: ‘analytical’ implementation; black curve: ‘ode’ implementation. Dashed yellow (resp. green) curve represent the slope of time increase as a power 4 (resp. 3) of the number of leaves.

Drury et al. (2016) also proposed an extension of the PM model accounting for the biogeographic history of lineages. In the case when each lineage is present in at most one location, the ‘analytical’ method can be extended, providing fast likelihood computation (see Appendix C.3). When there are lineages occurring in more than one location at the same time, we need to resolve numerically the ODE system in order to compute the likelihood of tip traits. While this is more time-consuming than finding a good ‘analytical’ reduction, the new implementation is more efficient than the one we previously proposed (Drury et al. 2016).

## Modeling trait evolution on coevolving clades

We illustrate how our framework can be used to study trait coevolution in scenarios of clade-clade interactions. We consider a simple model with two interacting clades (numbered 1 and 2), in which a given trait in clade 1 coevolves with another given trait in clade 2. Following the approach introduced above, we define *X_t_* the vector of trait values containing first the trait values for clade 1, and then the trait values for clade 2, and we write a stochastic differential equation specifying how trait value evolves along a single lineage *k*. In the spirit of the phenotype matching model, we propose here a formulation in which the trait of lineage *k* is attracted to (or repelled from, depending on the sign of *S*) the average trait value of the lineages it interacts with, plus (or minus) a shift:

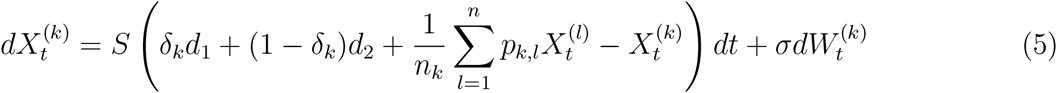

where *S* represents the attracting or repelling strength of species interactions on trait evolution, *d*_1_ (resp. *d*_2_) represents the shift for lineages from clade 1 (resp. clade 2), *σ* is the drift parameter, *δk* equals one if lineage *k* belongs to clade 1 and zero if it belongs to clade 2, *p^k,l^* equals one if lineages *k* and *l* interact and zero otherwise, 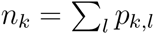 is the number of lineages interacting with lineage *k*, and *n* is the total number of lineages.

When *S* is positive, the trait value of lineage *k* is attracted to an optimal trait value given by the average trait value of the interacting species (plus a shift *d*_1_ or *d*_2_). An example of such a scenario of clade-clade matching mutualism is the coevolution between the length of floral tubes and the length of butterfly proboscis in a plant-pollinator mutualistic network (illustrated in Fig. 4). In this example, we assume that the optimal length of a butterfly proboscis is the average length of the plant floral tubes it pollinates plus a shift *d*_2_, while the optimal length of a plant floral tube is the average proboscis length of its butterfly pollinators plus a shift *d*_1_. With *d*_1_ + *d*_2_ = 0, both traits can reach their optimal state, leading to a stable situation with butterfly proboscis a bit longer (if *d*_1_ > 0) or shorter (if *d*_1_ < 0) than plant floral tubes. With *d*_1_ + *d*_2_ ≠ 0, traits cannot reach their optimal state, resulting in a runaway process where both traits tend to evolve toward an ever-moving optimum. For example, with positive *d*_1_ and *d*_2_, the butterflies proboscis tends to get longer to better access the nectar, while the floral tube also tends to get longer to force the butterfly's body to touch the stamen. The parameters *S* and *σ* control respectively the strength of the interaction effect and the rate of stochastic phenotypic change. The bigger *S*, the closer the traits will track the optimum; the bigger *σ*, the bigger the fluctuations around this optimum.

**Figure 4:**
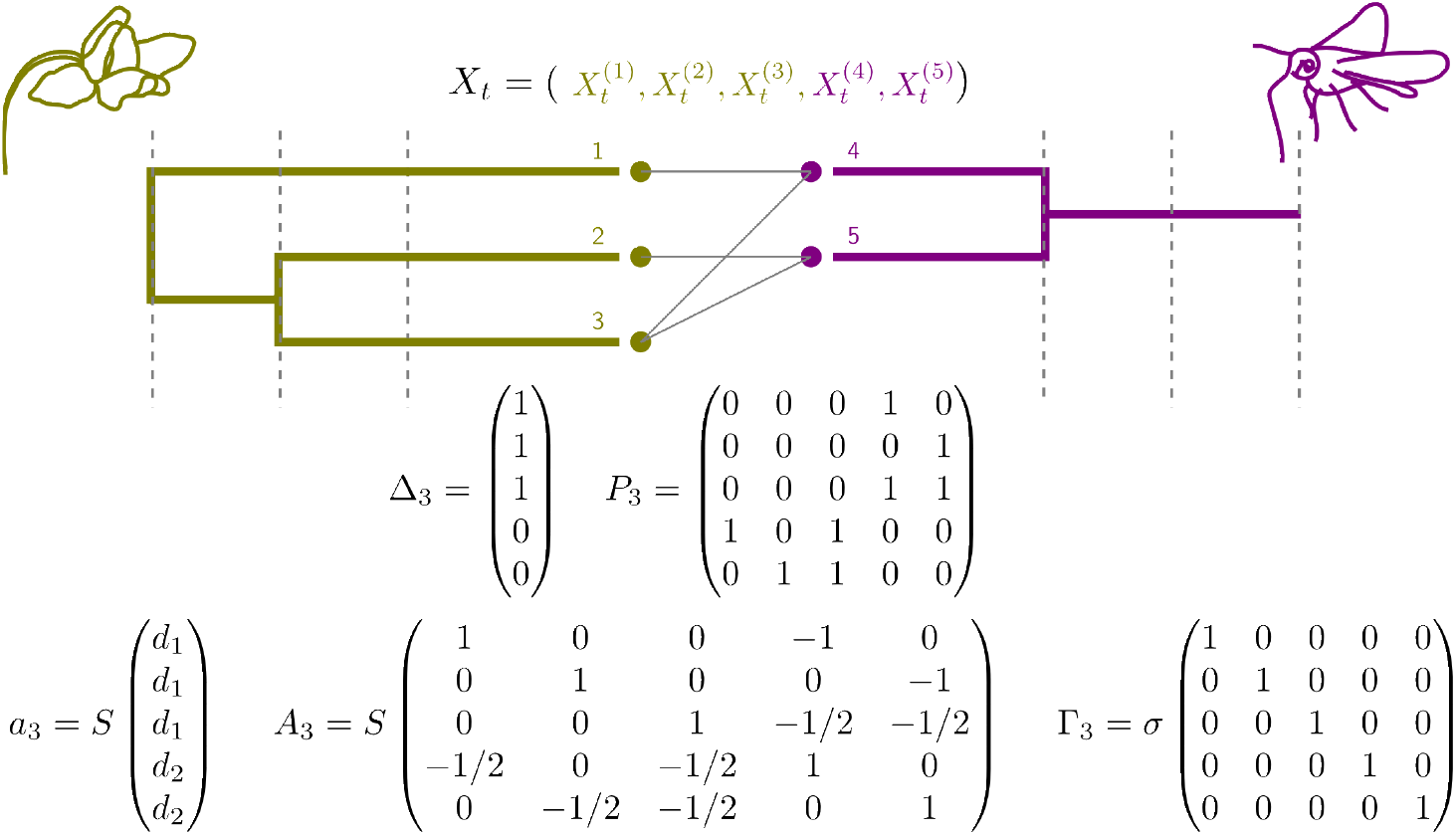
Hypothetical clade-clade coevolutionary scenario. Vertical dashed lines delimitate the successive epochs. The vector *X_t_* contains the trait values on the third (last) epoch, *P*_3_ is the matrix of network interactions, and *a*_3_, *A*_3_ and Γ_3_ together define trait evolution according to the clade-clade matching model defined in Equations (5) and (6).

When *S* is negative, the traits are repelled from the average trait value of the interacting species (plus a shift *d*_1_ or *d*_2_). This may capture natural situations of clade-clade competition driving trait displacement. Finally, some antagonistic interactions between traits could require to introduce two parameters *S*_1_ > 0 and *S*_2_ < 0 to capture match-vs-escape scenarios. For example, parasites might tend to develop cues matching those of their hosts while hosts develop cues to escape their parasites in a co-evolutionary arms race.

From Equation (5) we deduce the corresponding *a*, *A* and Γ on each epoch:

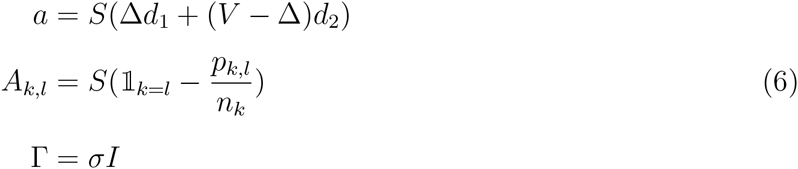

where Δ is the vector of elements *δ_k_* (see Fig. 4 for an illustration). Matrix *A* is in general not symmetric anymore, as all species *k* do not have the same number *n_k_* of species that they interact with.

As shown by Equation (6), entirely defining a model of clade-clade coevolution requires introducing a constant network of interaction on each epoch (the *P* matrix with elements *p_k,l_*). We can potentially re-define epochs to account for events of change in the interaction network in addition to speciation and extinction events, thus allowing interaction networks to evolve along branches. In practice, we typically (at best) have access to the current interaction network (Fig. 4), but not the ancestral networks. A solution to this would consist in treating the ancestral P matrices as parameters of the model, and searching the ancestral network(s) that maximize the fit to the data. Another approach would consist in reconstructing ancestral networks according to rules regarding the inheritance of interactions at speciation times. Developing these approaches is outside the scope of the current study, but we have shown how to compute tip trait distributions once they are developed.

We illustrate the computation of tip trait distributions for a model in which the ancestral networks are known: a generalist model where all species from clade 1 interact with all species from clade 2. We consider a ‘Generalist Matching Mutualism’ model of trait evolution (thereafter referred to as GMM, and illustrated in Fig. 5a), which is captured by Equation (5) with *S* positive and *p_k,l_* = 1 for any two lineages *k* and *l* from different clades and *p_k,l_* = 0 for any two lineages *k* and *l* from the same clade. Given that the model fits within our framework, we know that the trait distribution at the tips is Gaussian, and we can compute the expectation vector and covariance matrix corresponding to the model using Equations (3a, 3b), which we can reduce for this specific model in order to speed up the computation (Appendix C.4).

**Figure 5:**
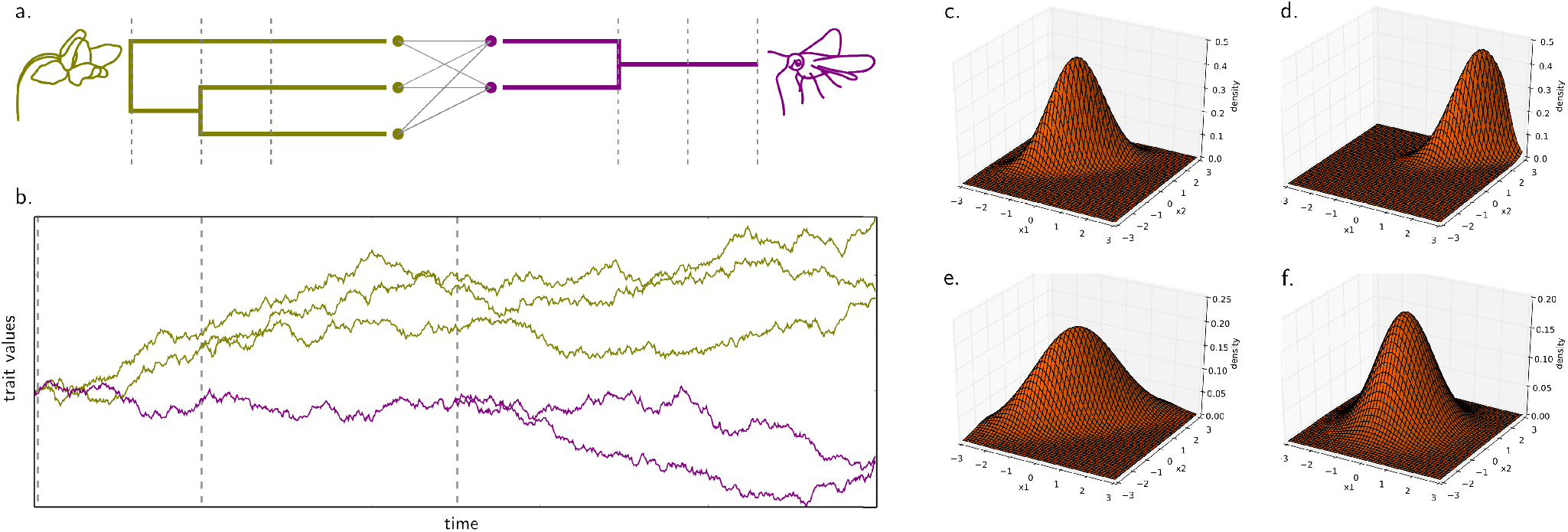
Trait evolution under the Generalist Matching Mutualism (GMM) model a) an illustrative generalist network of interactions between two clades. Vertical dashed lines delimitate the successive epochs. b) One realisation of trait evolution through time on the two phylogenies, with (*m*_0_, *υ*_0_) = (0, 0) and (*S, d*_1_, *d*_2_, *σ*) = (1, *−*1, 1, 1), simulated using the Euler-Maruyama scheme (see Appendix D.1). cdef) Expected tip distribution for the average trait value in each clade, with parameter values (*S, d*_1_, *d*_2_, *σ*) = c) (2, −1, 1, 1), d) (2, 0, 2, 1). e) (2, −1, 1, 1.5), f) (0.2, −1, 1, 1).

The tip distribution is relatively fast to compute (e.g. in the order of 0.8 seconds with two 100-tip trees on a desktop computer), such that fitting the model by maximum likelihood or in a Bayesian framework should not be problematic for trees with a few hundred tips. However, we do not aim here to carry an in-depth study of this particular model, nor to fit it to empirical data. Rather, we use our ability to rapidly compute tip trait distribution to get a first glimpse of the model behaviour under distinct sets of parameter values.

In Figure 5 (c,d,e,f), we plotted the distribution of the average 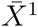 of trait values in clade 1 and the average 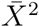 of trait values in clade 2 for traits evolving under the GMM model with four parameter sets chosen to lead to four distinct qualitative behaviours. From Equation (5), we can easily show that under GMM 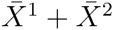 is a drifted Brownian motion with drift term *S*(*d*_1_ + *d*_2_) and 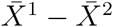 is an OU process with optimum (*d*_1_ − *d*_2_)/2 and selection strength *S*. The shift parameters *d*_1_ and *d*_2_ thus directly determine the position of the optimum of the distribution. When *d*_1_ = −*d*_2_, the more likely values for the two average traits 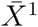 and 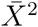 are such that 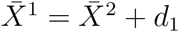 (see Fig. 5c). In contrast, when *d*_1_ ≠ *d*_2_, the two communities have optimal trait values that are non-compatible, and the traits will tend to increase (resp. descrease) if *d*_1_ + *d*_2_ > 0 (resp. *d*_1_ + *d*_2_ < 0) (see Fig. 5d). The position of the peak in the tip distribution will thus depend also on the depth of the root and on the value of the parameter *S*. Moreover, the parameter *S* plays an important role in the hump thickness: the bigger *S*, the more constrained 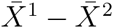 around (*d*_1_ − *d*_2_)/2 (see Fig. 5f). The parameter *σ* also plays a role in the thickness of the hump, but in the orthogonal direction: increasing *σ* flattens the distribution by allowing different 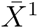 and 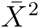 values while retaining the constraint on 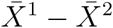 (see Fig. 5e).

### Implementation

Our framework is implemented in the R package ‘RPANDA’ (Morlon et al. 2015), including functions: 1) to compute tip distributions, 2) to simulate trait evolutionary trajectories using the Euler-Maruyama scheme (see Appendix D.1) and tip data by drawing from the expected tip distribution, and 3) to infer model parameters by maximum likelihood. In the most general user-defined use of our framework, the input is one or several phylogenetic tree(s) and the collection of (*a_i_, A_i_*, Γ*_i_*) matrices on each epoch that define a specific model. In this case, the tip distribution is computed using the most general ‘ode’ method that solves numerically the ODEs. In addition, we implemented all models mentioned in Table 1 as well as GMM with the fastest described algorithm to compute their tip distribution. In Appendix E, we provide a tutorial explaining the structure of our code and illustrating how to use it.

## Discussion

We developed a modeling framework for traits coevolving in coevolving lineages and clades. We showed that under a wide variety of models where the evolution of a given trait on a given lineage is linearly related to its own value and the value of other traits on the same lineage, of the same trait on other lineages, and/or of other traits on other lineages, the expected tip trait distribution is Gaussian. We showed how to compute this tip distribution in general, as well as for specific models, including classical models of phenotypic evolution and new models of clade-clade coevolution.

All classical models of phenotypic evolution, such as univariate and multivariate BM, OU, ACDC and DD fit within our framework. They correspond to the situation where the evolution of traits on a given lineage is independent of trait values on other lineages. For these models, we already know that the tip trait distribution is Gaussian. However, finding the relevant computation of the expectation vectors and covariance matrices associated with each model in the dense literature of comparative phylogenetics can be overwhelming for neophytes. Our Appendix B unifies these computations under a common formalism, providing both the expressions for the various existing models and their mathematical underpinning. This is done in the context of trees that are not necessarily ultrametric, meaning that all models can be applied to phylogenies including fossils. We hope that this Appendix can serve as a useful review for navigating phylogenetic approaches for understanding trait evolution.

The fact that the distribution of traits remains Gaussian when traits from different lineages coevolve is a new result. It is also a convenient result, because it means that computing the tip distribution only requires computing the expectation vector and covariance matrix associated with the different models. For example, we used this result in Drury et al. (2016) to compute tip trait distributions for the phenotype matching model (Nuismer and Harmon 2014) and fit it to comparative data by maximum likelihood. Here we vastly extend the set of potential coevolutionary models for which tip trait distributions can be computed and provide two general approaches for computing the expectation vector and covariance matrix. One of these two approaches (the ‘ode’ approach) consists in numerically integrating a set of ODEs. This is the approach that was used in Drury et al. (2016). The other approach (the ‘analytical’ approach) involves computing integrals and is more efficient when these integrals can be analytically reduced, which depends on the form of the model. Applying the ‘analytical’ approach to the PM model, we greatly improved its computational efficiency.

We provide a framework for computing tip trait distributions for a wide class of models accounting for within-clade and clade-clade interactions. We hope that this flexibility will foster the development and study of various models adapted to the specificities of particular scientific questions and biological systems. We did not study at length a particular coevolutionary model in this paper, but the PM model was thoroughly studied elsewhere (Drury et al. 2016). The Generalist Matching Mutualism model that we introduce here can be seen as a clade-clade analogue to the PM model. Both models are ‘generalist’ in the sense that all lineages are assumed to interact (within-clade in the case of PM and between clades in the case of GMM). This assumption can be relaxed by incorporating additional information. In our biogeographic models for example, lineages can only interact if they are sympatric (Drury et al. 2016). More generally, any information or hypothesis concerning the network of interactions between lineages can be accounted for into the *A* matrices.

There are two main limitations to the modeling framework presented here. The first one is that trait evolution is always assumed to respond linearly to trait values in other lineages. Thus, non-linear effects such as a stronger selection for divergence when phenotypes are similar cannot be accounted for. The second one is the issue of model and parameter identifiability, in particular in the absence of fossils. A Gaussian distribution in ℝ*^nd^* can potentially allow identifying several models and parameters, but there are distinct combinations for which a similar (or even identical) distribution is expected. For example, we already know that parameters of the OU model are non identifiable on phylogenies with only extant species (Ho and Ané 2014) and that OU and AC have identical tip distributions on ultrametric trees (Uyeda et al. 2015). Thus, while we wrote our framework in all generality, with *a*, *A* and Γ encompassing as many parameters as desired, and parameters that potentially vary between epochs, it is clear that simplifying assumptions need to be made in order to reduce this parameter space. Identifiability cannot always be checked analytically, as in the case of the OU and AC models. In addition, there can be differences between theoretical and *de facto* identifiability, with models that are identifiable in theory but are difficult to identify in practice. For example, we can show analytically that *ψ* and *S* from the PM model are theoretically identifiable, but in practice in most cases only *ψ* + *S* can be estimated with precision. Also, *de facto* identifiability depends on the data available, such as the size and shape of a particular phylogeny, and whether it includes fossils or not (Slater et al. 2012). Furthermore, models taking into account interactions among lineages will have to assess the influence of extinct lineages in the past. This has been studied in Drury et al. (2016) for the PM model, by simulating trait evolution on trees including dead branches, before fitting the model on the reconstructed tree only. Our recommendation is to check identifiability on a case-by-case basis, by fitting the set of models under consideration to trait datasets simulated directly on the specific empirical phylogenies in hands. We provide the tools for rapidly simulating tip values under various models by sampling expected distributions.

One of the most challenging and exciting developments that we see ahead is to move from generalist models to models that account for specific interaction networks. We show in this paper how to compute tip trait distributions for such models, assuming that the ancestral networks are known. While some fossil species interaction networks have been compiled (Dunne et al. 2008), such data is typically not available. Thus, if we are to really understand if and how species interactions affected long-term phenotypic evolution, we need to start developing models for reconstructing ancestral networks, analogous to the use of ancestral biogeographic models (see Ronquist and Sanmartín 2011 for a review) to incorporate biogeography into models of phenotypic evolution (see e.g. DD + GEO or MC + GEO, in Drury et al. 2016). Interestingly, our modeling framework could provide an approach to do so, informed by species phylogenies, the interaction network of present day species, and current species phenotypes. Indeed, rather than assuming that the ancestral networks are known, we could treat them as additional parameters to optimize upon, and find the ancestral networks that maximize the likelihood of the current data. Whether there will be enough information in the data to distinguish the probability of alternative ancestral networks remains to be tested, but the observed phylogenetic signal in empirical networks of interactions is encouraging (Ives and Godfray 2006; Rafferty and Ives 2013; Hadfield et al. 2014; Hayward and Horton 2014; Martín González et al. 2015). Our ability to distinguish the probability of alternative ancestral networks will be increased by proposing various scenarios regarding the inheritance of interactions at speciation times, such as scenarios in which daughter species interact with many or few of the species that interacted with their mother lineage. These upcoming developments can draw upon the existing literature on the cophylogeny problem (Conow et al. 2010), and will certainly have an important role to play in the on-going effort of understanding the evolution of species interaction networks (Loeuille and Loreau 2005; Martinez 2006; Nuismer et al. 2013).

Our framework for modeling trait evolution on phylogenetic trees includes most previously proposed models and can be used to develop a series of new models of within-clade and clade-clade coevolution. We hope that this will motivate new theoretical and empirical applications aimed at unravelling how species interactions evolve and influence phenotypic evolution over macro-evolutionary time-scales.

## Acknowledgements

We are very grateful to Jonathan Drury, Florence Débarre and Luke Harmon, as well as members of our research teams for helpful discussions and comments on previous versions of the manuscript. MM acknowledges his PhD funding from the École Normale Supérieure; AL acknowledges funding from the Center for Interdisciplinary Research in Biology (Collège de France); HM acknowledges funding from ERC, grant ERC-CoG PANDA.

## Appendix

### A Derivation of the distribution in a general setting

#### A.1 The distribution of trait values is Gaussian

Recall that a vector is Gaussian if all linear combination of its components follows a normal distribution. We will thus show by induction that all linear combinations of the traits follow a normal distribution.

The process of trait evolution starts either at the stem root with a vector of size *d* defined by the initial conditions 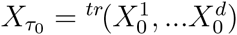, or at the crown root with a vector of size 2*d* defined by the initial conditions: 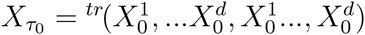, or at any other step, provided the initial conditions are Gaussian by assumption.

Now, assume that 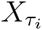 is a Gaussian vector.

Then, 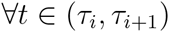, after integration we have the following closed expression for the value of the process *X_t_*.

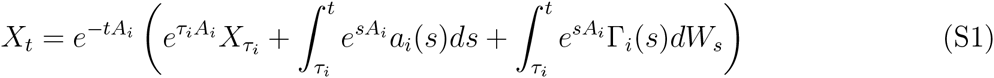

Moreover, we have, for any deterministic function Φ (Gardiner et al. 1985),

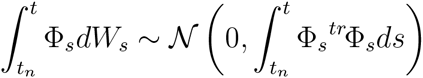

Hence, *X_t_* is a linear combination of Gaussian vectors, which makes it a Gaussian vector.

Last, suppose that at time 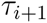, the *j*th branch splits, in which case the vector grows. All linear combinations of the components of *X_t_* at time 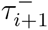 have a normal distribution. And the *d* additional components added at time 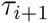 belong to the components at time 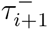. It follows that all linear combinations of the new vector still have a normal distribution.

#### A.2 Integrating the evolution of the distribution on each epoch

Still assuming that we know the (Gaussian) distribution of 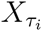 at the beginning of an epoch (*τ_i_, τ_i_*_+1_), a few more lines allow us to provide a closed formula for the distribution of *X_t_* at all time *t* ∈ (*τ_i_*, *τ*_*i*+1_). Indeed, using Equation (S1), and the fact that, if *X* and *Y* are two independent Gaussian vectors with expectation vectors respectively *m_X_* and *m_Y_* and covariance matrices respectively Σ*_X_* and Σ*_Y_*, then:

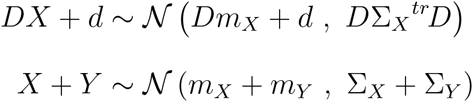

It thus follows that, ∀*t* ∈ [*τ_i_*, *τ*_*i*+1_],

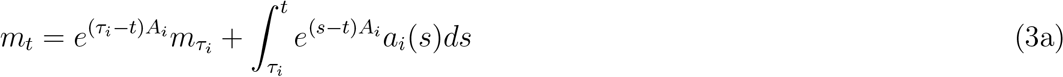

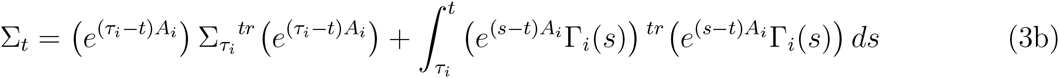

Applying these equations for *t* = *τ*_*i*+1_ thus gives the distribution of the trait vector at time *τ*_*i*+1_, which is the result stated in Equations (3a, 3b) in the main text.

Remark that, unless one of the very first branches immediately dies at the beginning of the process at a fixed initial condition, the density of the tip distribution has support in ℝ*^nd^*. One can check that Σ*_t_* stays positive definite (implying that det Σ*_t_* ≠ 0), even when some Γ*_i_* are not positive definite (except the first one).

#### A.3 Evolution of the distribution through ODE resolution

The expectation and covariance formulae provided in Equations (3a, 3b) require to deal with an integral which is not always straightforward to compute. Alternatively, one can prefer to take the derivative of this expression, get a set of ODEs verified by the expectation and covariance elements through each epoch, and subsequently integrate the ODE system. We show now another way to derive this set of ODEs.

First, we write the stochastic differential equation on any epoch (*τ_i_*, *τ*_*i*+1_) and for each trait *k*, which is given in the most general setting by:

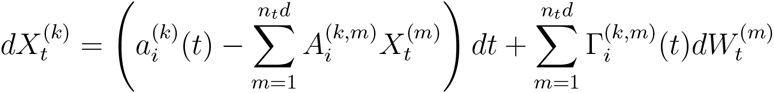

Itô’s formula (Gardiner et al. 1985) then gives us:

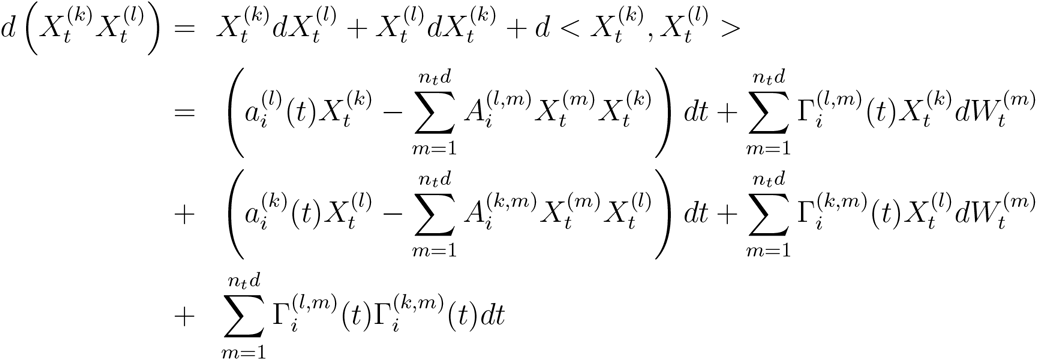

Taking the expectation, it follows that

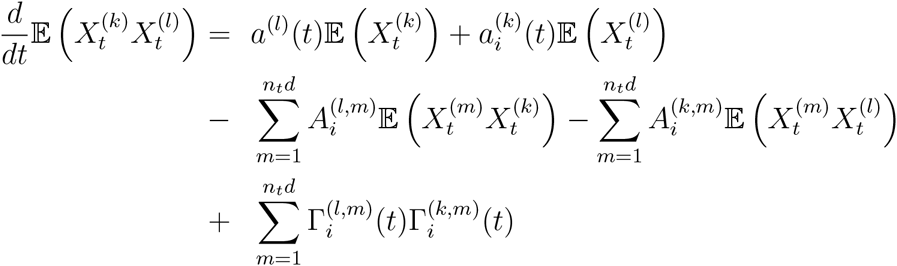

In the same fashion, we get

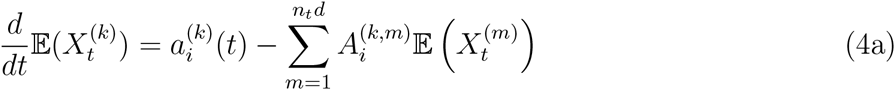

This leads to

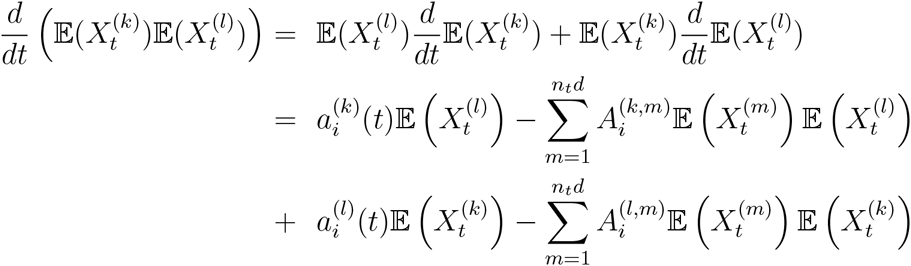

Putting together these different parts gives us the ODE satisfied by all covariances:

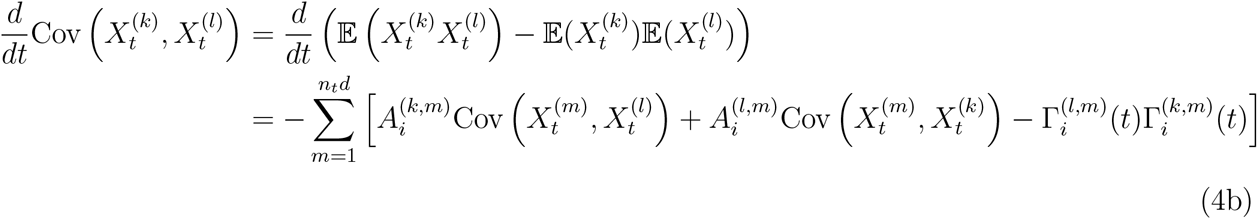

Note that in a vectorial formalism with the expectation vector *m* and covariance matrix Σ, these sets of ODEs can be written equivalently as follows

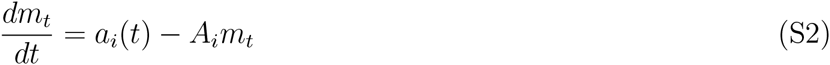

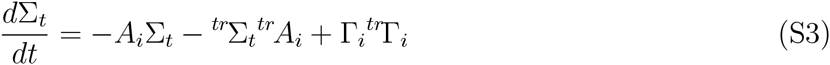

### B Distribution for some models without interactions between lineages

#### B.1 Distribution of classic univariate models

We present in this section how previously known results of analytic tip distribution of univariate models fit in, and can be rediscovered with, our framework. Results are summarized in Table S1.

The scheme is identical for each model:

1. Reduce Equations (3a, 3b) or (4a, 4b) according to the model.
2. Look for an analytical solution at any time *τ_i_*, by calculating manually the expectations and covariances at *τ*_1_, *τ*_2_, *τ*_3_, …
3. Prove by induction that the analytical solution holds at any time *τ_i_*.

We call *t_k,l_* the time of the most recent common ancestor to lineages *k* and *l*, and *t_k,k_* the death time of lineage *k*, equal to *T* if it survives until present (see Fig. S1). We further note 1*k* _alive_(*t*) the quantity that equals one if lineage *k* is alive at time *t* and zero otherwise, and 1_*k*=*l*_ that equals one if *k* = *l* and zero otherwise. Last, *t*_1_ ∧ *t*_2_ stands for the minimum of the two values *t*_1_ and *t*_2_.

The unity vector (vector full of 1) is denoted by *V*, *I* refers to the identity matrix (diagonal matrix with diagonal values equal to 1), and *U* refers to the unity matrix (matrix full of 1). Their size is the same as the size of the vector of traits *X_t_* considered. Considering non-ultrametric trees including fossils amounts to replacing vector *V* and matrices *I* and *U* by their homologs *V*_alive_, *I*_alive_ and *U*_alive_, where the subscript specifies that the vector and matrices have 0 on lines and columns corresponding to lineages that are extinct in the given epoch.

##### Brownian Motion (BM).—

We show how to get the well-known expression of the distribution of a trait evolving under BM, on non-necessarily ultrametric trees. We take *a* = *bV*_alive_, *A* = 0 and Γ = *σI*_alive_, i.e. the process follows the equation:

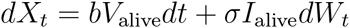

**Figure S1:**
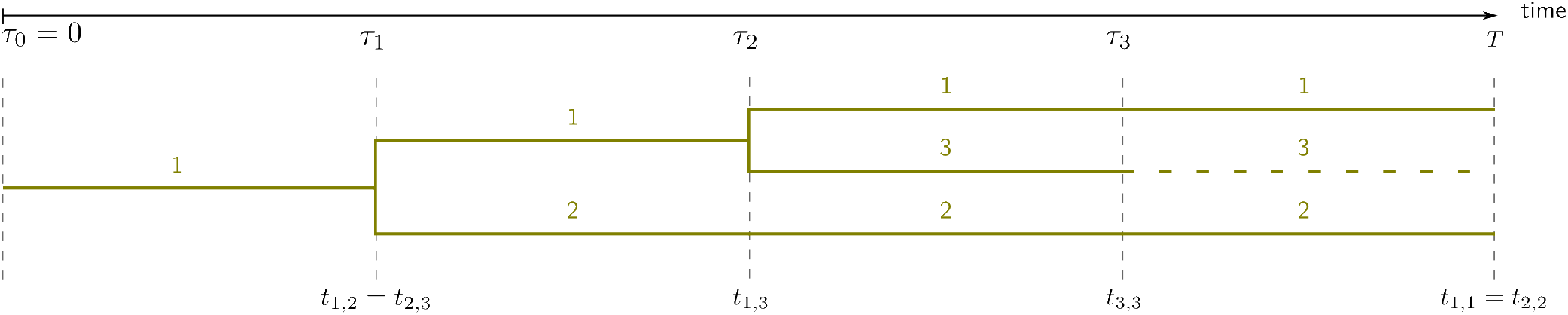
Formalism used in analytic formulae presented in Table S1.

**Table S1:**
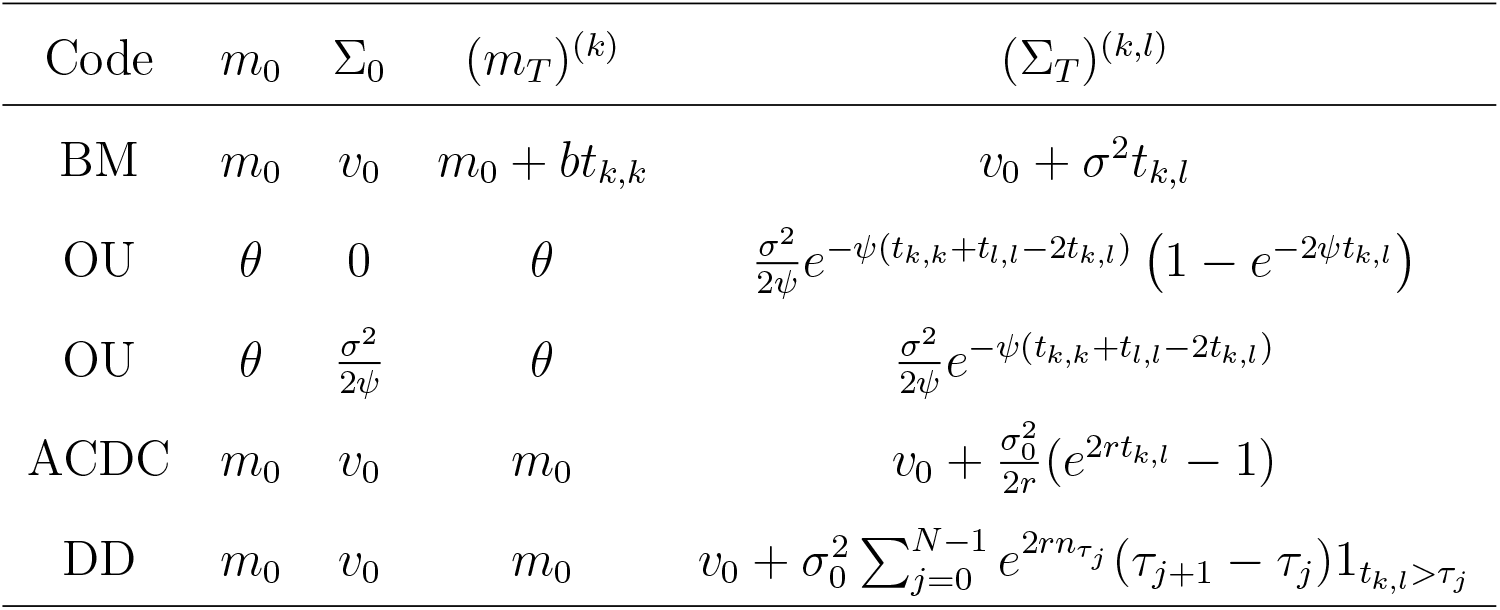
Analytic tip distribution for models without interactions between traits or lineages. We recall that *t_k,l_* is the absolute time of the most recent common ancestor to lineages *k* and *l*, and *t_k,k_* is the death time of lineage *k*, equal to *T* if it survives until present.

Equations (3a) and (3b) lead to the following recurrence formulae driving the law of *X_t_* on each epoch [*τ_i_*, *τ*_*i*+1_):

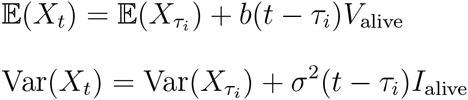

Alternatively, Equations (4a) and (4b) lead to the following recurrence formulae driving the law of *X_t_* on each epoch [*τ_i_, τ_i_*_+1_):

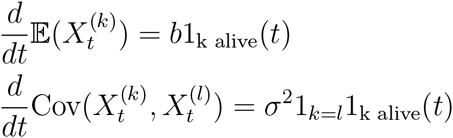

We can show by induction on *i* that for any *i* the expectation and covariance matrix at time *τ_i_* are such that, for any (*k, l*):

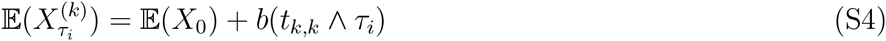

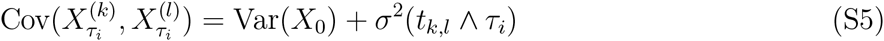

Indeed, we verify Equations (S4, S5) at step *i* = 1.

Now, suppose Equations (S4, S5) hold at step *n*. Using either Equations (3a, 3b) or (4a, 4b), we get:

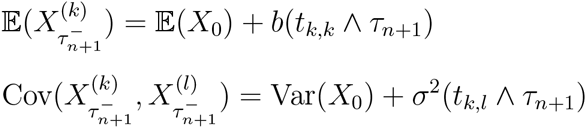

If *τ_n_*_+1_ is a death time of a lineage, Equations (S4, S5) are verified at step *n* + 1.

If *τ*_*n*+1_ is a branching time, we verify that the new lineage inherits the expectation and covariances of its mother, as well as the same coalescence times with other lineages. It also follows that Equations (S4, S5) are verified at step *n* + 1.

Finally, by induction, we get the tip distribution:

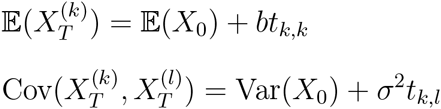

##### Ornstein-Uhlenbeck (OU).—

We can get another well-known distribution for a trait evolving under an Ornstein-Uhlenbeck process on a tree. We take *a* = *ψθV*_alive_, *A* = *ψI*_alive_ and Γ = *σI*_alive_, i.e. the process follows the equation:

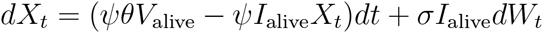

Expressions (3a) and (3b) simplify into the following recurrence formulae:

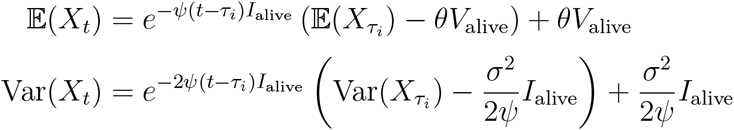

Alternatively, here again, one can prefer to apply Equations (4a) and (4b):

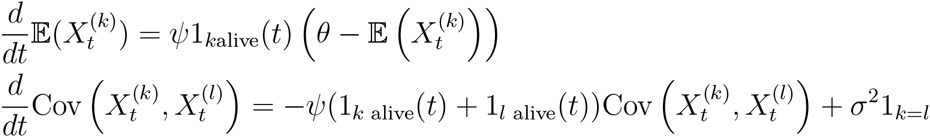

We can show by induction that for any epoch *i*, the expectation and covariance matrix at time *τ_i_* are such that, for all (*k, l*):

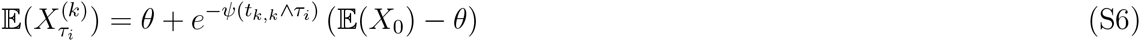

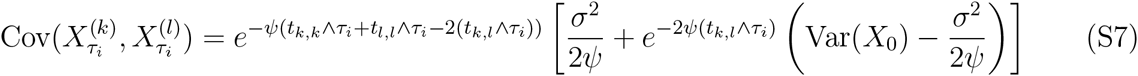

Indeed, we verify Equations (S6, S7) at step *i* = 0.

Now, suppose Equations (S6, S7) hold at step *n*. Using either Equations (3a, 3b) or (4a, 4b), we get:

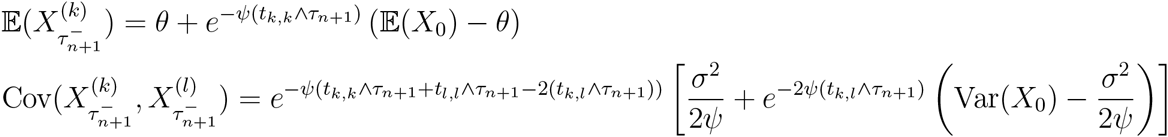

If *τ*_*n*+1_ is a death time of a lineage, Equations (S6, S7) are verified at step *n* + 1.

If *τ*_*n*+1_ is a branching time, we verify that the new lineage inherits the expectation and covariances of its mother, as well as the same coalescence times with other lineages. It also follows that Equations (S6, S7) are verified at step *n* + 1.

Finally, by induction, we get the tip distribution:

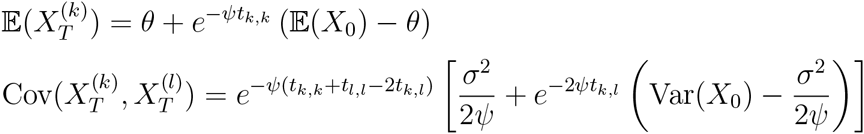

Two classes of initial distributions are typically considered in the literature:

1. If we consider a process starting at *X*_0_ = *θ* (i.e. with 𝔼(*X*_0_) = *θ* and Var(*X*_0_) = 0), we get the following expectation vector *m_T_* and covariance matrix Σ*_T_* at the tips:

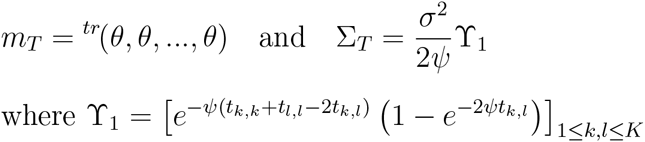
2. When *ψ* > 0, if we consider a process starting under its stationary distribution (i.e. 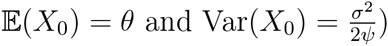, it simplifies into the following expectation vector and covariance matrix:

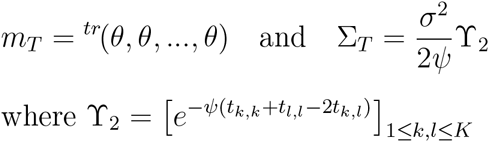

##### ACDC (accelerating or decelerating rate).—

In the ACDC process, the rate of phenotypic evolution varies exponentially through time, with *a* = 0, *A* = 0 and Γ = *σ*_0_*e^rt^I*_alive_ (here, *r >* 0). The process follows the equation:

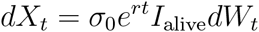

Here again, we can simplify Equations (3a, 3b) or (4a, 4b). With Equations (3a, 3b), we get the following recurrence formulae driving the law of *X_t_* on each epoch (*τ_i_*, *τ*_*i*+1_):

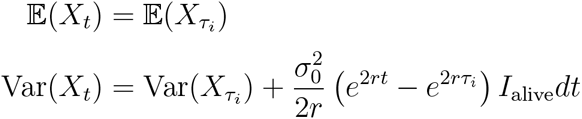

We can show by induction that for any *i*, the expectation and covariance matrix at time *τ_i_* are such that, for any (*k, l*):

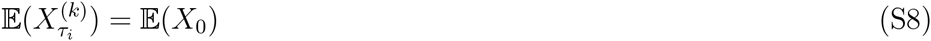

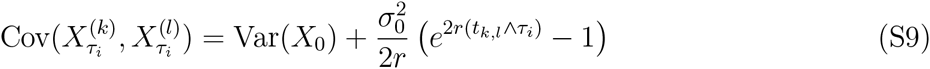

Indeed, we verify Equations (S8, S9) at step *i* = 0.

Now, suppose Equations (S8, S9) hold at step *n*. Using either Equations (3a, 3b) or (4a, 4b), we get:

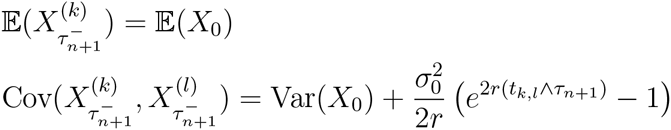

If *τ*_*n*+1_ is a death time of a lineage, Equations (S8, S9) are verified at step *n* + 1.

If *τ*_*n*+1_ is a branching time, we verify that the new lineage inherits the expectation and covariances of its mother, as well as the same coalescence times with other lineages. It also follows that Equations (S8, S9) are verified at step *n* + 1.

Finally, by induction, we get the tip distribution:

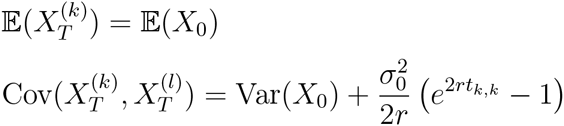

##### ACDC and OU processes lead to the same present-time distributions on ultrametric trees.—

This has been shown previously in Uyeda et al. 2015. More precisely, OU is equivalent to a model with accelerating rates at present, and only on ultrametric phylogenies.

Looking at expressions of expectations and covariance matrices under ACDC and OU with initial conditions *X*_0_ = *θ*, we see that we can choose parameters such that we get the exact same distribution. First take 𝔼(*X*_0_) = *θ*: the two expectation vectors are identical. Moreover, we can choose parameters such that the covariance matrices are equal:

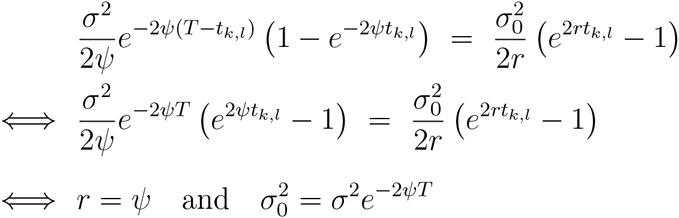

Note that this no longer holds on non-ultrametric trees, neither with different initial conditions on the OU.

##### Diversity-Dependent (DD).—

In the DD process, the rate of phenotypic evolution is fixed at the base of the tree and varies exponentially with the number of lineages in the reconstructed phylogeny, with *a* = 0, *A* = 0 and *B*(*t*) = *σ*_0_*e^rnt^ I*_alive_. The process follows the equation:

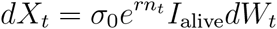

Equations (3a, 3b) lead to the following recurrence formulae driving the law of *X_t_* on each epoch (*τ_i_*, *τ*_*i*+1_):

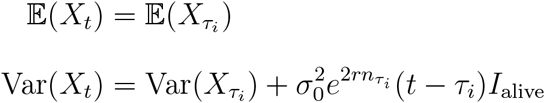

Note that, alternatively, one can again prefer to apply Equations (4a, 4b). We can then show by induction that for any *i*, the expectation and covariance matrix at time *τ_i_* are such that, for any (*k, l*):

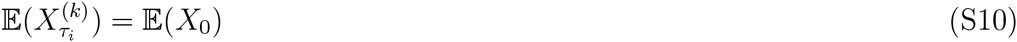

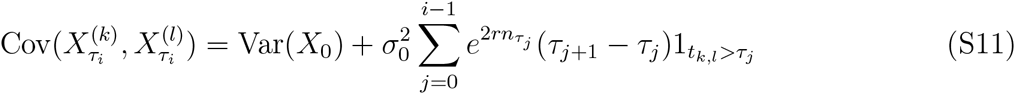

Indeed, we verify Equations (S10, S11) at step *i* = 0.

Now, suppose Equations (S10, S11) hold at step *n*. Using either Equations (3a, 3b) or (4a, 4b), we get:

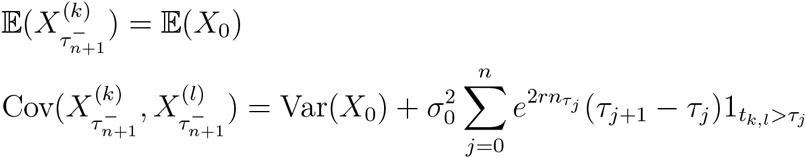

If *τ*_*n*+1_ is a death time of a lineage, Equations (S10, S11) are verified at step *n* + 1.

If *τ*_*n*+1_ is a branching time, we verify that the new lineage inherits the expectation and covariances of its mother, as well as the same coalescence times with other lineages. It also follows that Equations (S10, S11) are verified at step *n* + 1.

Finally, by induction, we get the tip distribution at present time *τ_N_* = *T*:

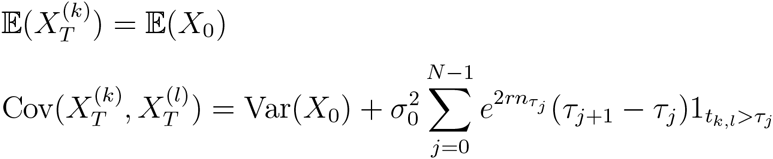

#### B.2 Distribution of classic multivariate models

The same methodology applies to classic multivariate models that incorporate interactions between traits within lineages but not between lineages. In our formalism, for all *i*, *A_i_* and Γ*_i_* are block diagonal, with *d* × *d* blocks on the diagonal corresponding to the traits within each lineage. We call these blocks respectively *A^∗^* and Γ*^∗^*. Moreover, the vector *a_i_* is the repetition of identical sequences *a^∗^* of *d* elements.

Writing the matrix products in Equations (3a, 3b) provides us with *d × d* blocks that behave identically on each epoch. Indeed, we can use:

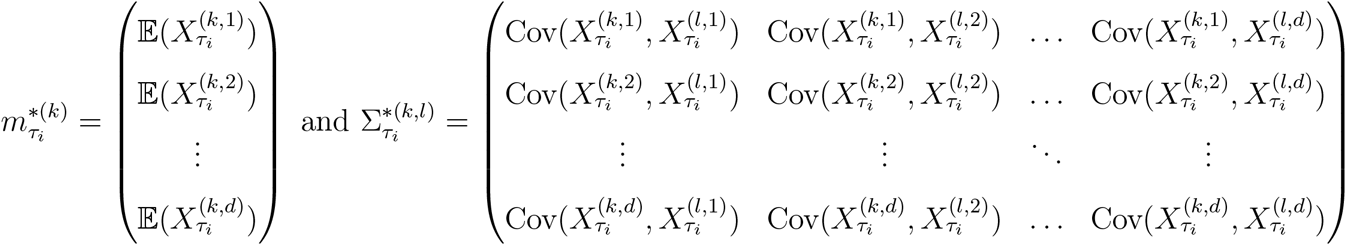

In which case Equations (3a, 3b) lead to the recurrence formulae:

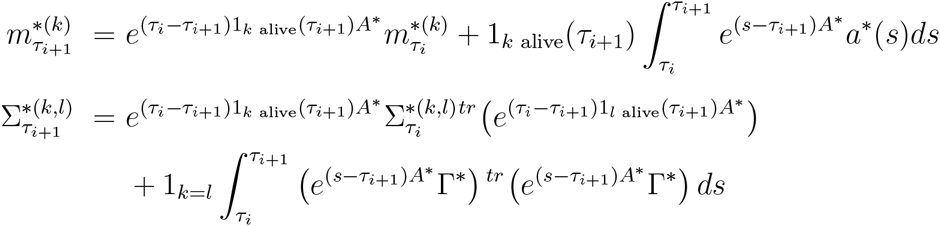

We can then prove by induction that for any epoch *i* and any pair of lineages (*k, l*)

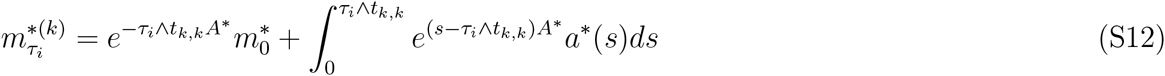

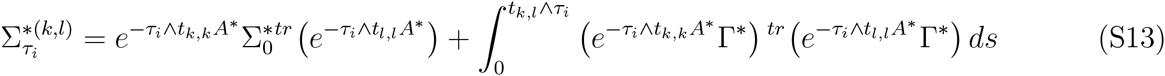

Indeed, we verify Equations (S12, S13) at step *i* = 0.

Now, suppose Equations (S12, S13) hold at step *i*. Using Equations (3a, 3b), we get:

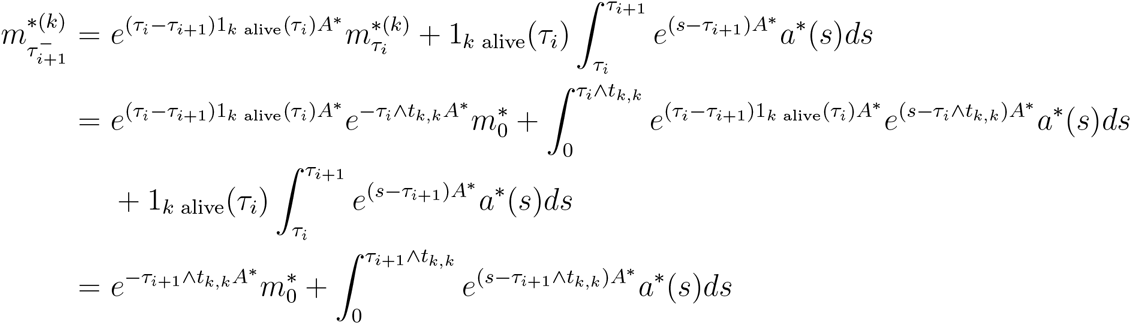

as well as:

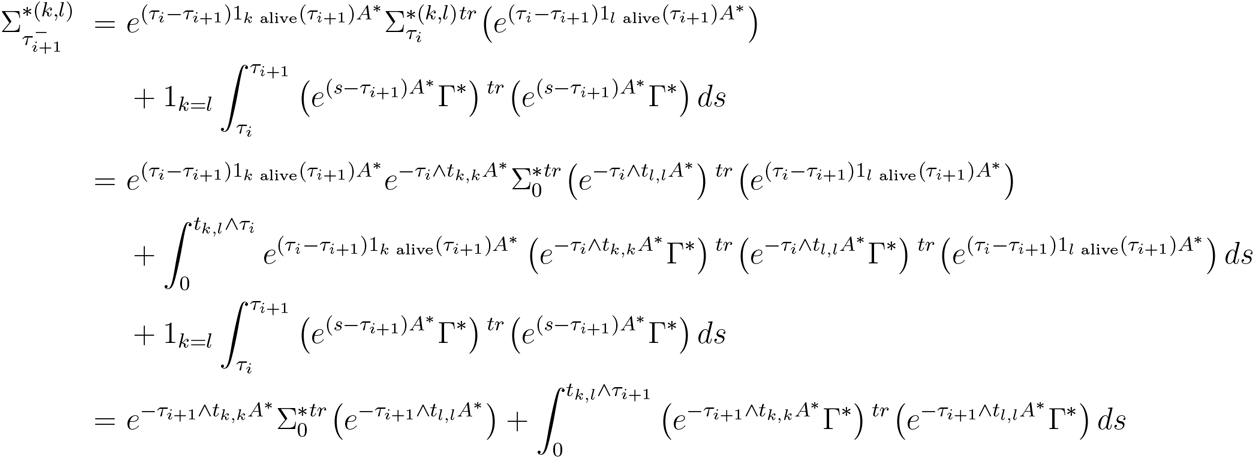

If *τ*_*i*+1_ is a death time of a lineage, Equations (S12, S13) are verified at step *i* + 1.

If *τ*_*i*+1_ is a branching time, we verify that the new lineage inherits the expectation and covariances of its mother, as well as the same coalescence times with other lineages. It also follows that Equations (S12, S13) are verified at step *i* + 1.

Finally, by induction, we get the tip distribution:

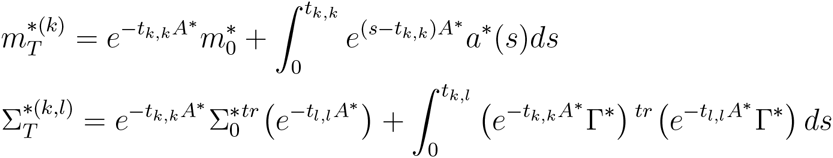

##### OU-BM model.—

As a first illustration, consider a model with *d* = 3 traits with equation on each epoch and on each lineage *k* as follows:

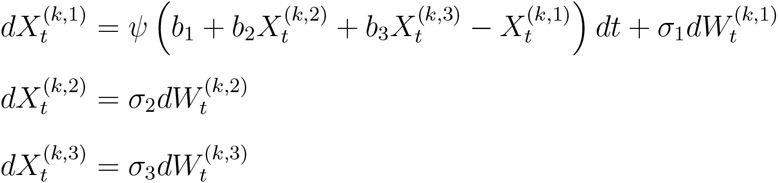

These equations describe the evolution of two independent traits evolving following a BM (traits 2 and 3), and one trait following an OU with optimal trait value given by a linear combination of traits 2 and 3. Its main interest is to infer the dependence of one trait to two other independent traits on a phylogeny. Knowing the distribution at the beginning of a given epoch, we use Equations (3a, 3b) to compute the distribution at the end of the epoch.

*A* is block-diagonal with the following blocks *A^∗^*:

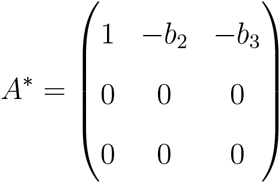

Writing Δ = *s* − *τ*_*i*+1_, it follows that *e*^Δ*A_i_*^ is block diagonal with 3 × 3 elements given by:

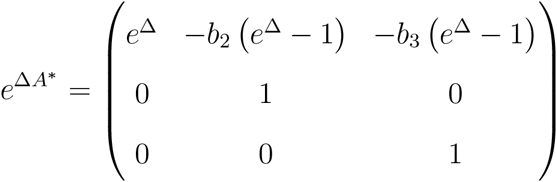

Moreover, Γ*_i_* is block-diagonal with diagonal blocks:

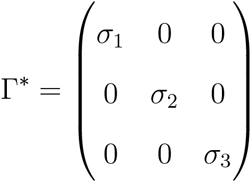

The matrix product (*e*^Δ*A_i_*^ Γ*_i_*)*^tr^*(*e*^Δ*A_i_*^ Γ*_i_*) is thus block-diagonal with 3 × 3 blocks:

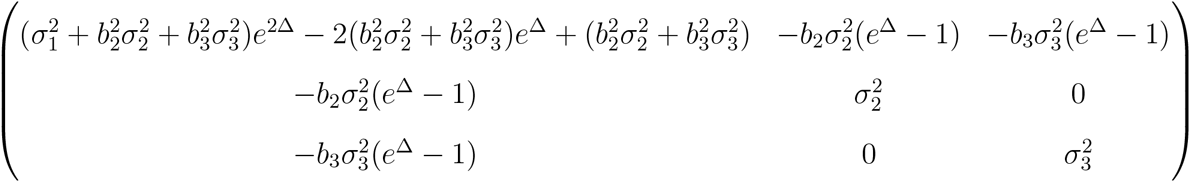

These matrices can be used to compute 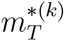 and 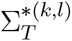, with the help of Equations (S12, S13).

##### OU-OU model.—

Consider now a model with *d* = 2 traits with equation on each epoch and on each lineage *k* given by:

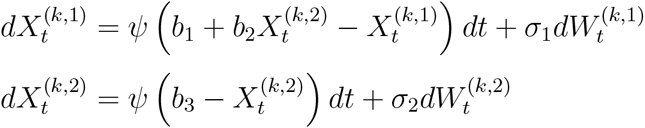

These equations describe the evolution of one trait evolving following an OU (trait 2), and one trait following an OU with optimal trait value given by an affine transformation of trait 2. Its main interest is to infer the dependence of one trait to another trait on a phylogeny. Knowing the distribution at the beginning of a given epoch, we use Equations (3a, 3b) to compute the distribution at the end of the epoch.

*A_i_* is block diagonal, with the following 2 × 2 blocks *A^∗^*:

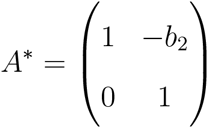

Again, writing Δ = *s* − *τ*_*i*+1_, it follows that *e*^Δ*A_i_*^ is block diagonal with 2 × 2 elements given by:

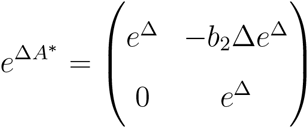

Moreover, Γ*_i_* is diagonal with repeated values:

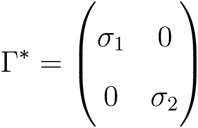

The matrix product (*e*^Δ*A_i_*^ Γ*_i_*)*^tr^*(*e*^Δ*A_i_*^ Γ*_i_*) is thus block-diagonal with 2*2 blocks:

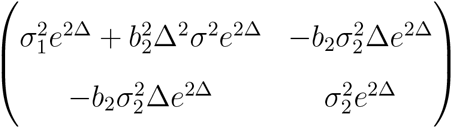

These matrices can be used to compute 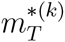 and 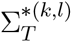, with the help of Equations (S12, S13).

### C Distribution for some models with interactions between lineages

#### C.1 Distribution with a constant, A symmetric, and *Γ* = *σI*

When Γ = *σI* and *A* is symmetric, Equations (3a, 3b) become:

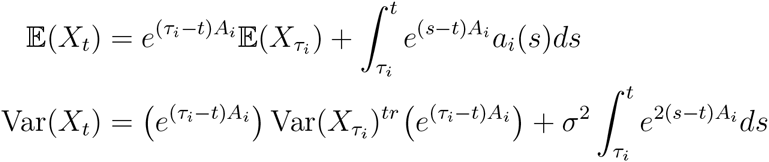

If *A_i_* is symmetric with coefficients in ℝ, it can be diagonalized by orthogonal passage matrices: we can exhibit a matrix *Q* verifying *^tr^QA_i_Q* = Λ*_i_* is diagonal and *Q^−^*^1^ = *^tr^Q*.

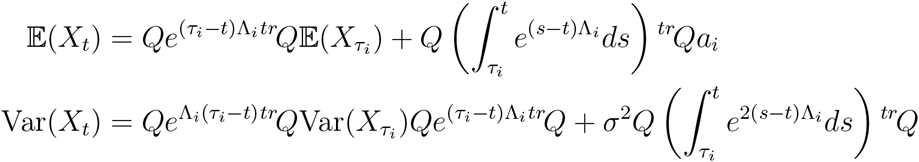

This is the expression that we need for the numerical integration, in particular, of the phenotype matching model.

Note that with *A* diagonalizable but not symmetric, Equations (3a, 3b) can also be reduced, but the transposition of *A* is no longer *A*, and it does not lead exactly to the same expression.

#### C.2 The phenotype matching (PM) model

We consider here the phenotype matching model introduced in Nuismer and Harmon (2014), with the following equation describing the evolution of any trait *k* on each epoch:

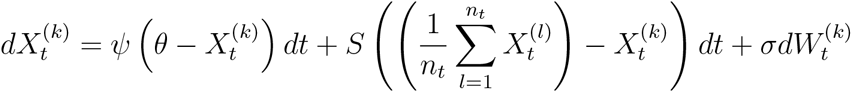

We introduce the line vector *u*, with value *u_j_* that equals 1 if lineage *j* is alive, and 0 otherwise. In order to use our framework, we further want to express the model in the form given by Equation (1). This is achieved by taking:

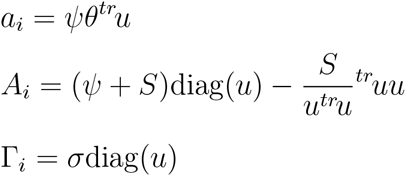

where diag(*u*) is the diagonal matrix with diagonal elements the elements of the vector *u*.

First, the tip distribution can be computed using the general algorithm that numerically resolves the set of ODEs given in Equations (4a, 4b). Second, the PM model falls within the class of models studied in the previous section, that is, with a symmetric *A* matrix. The tip distribution can thus be numerically computed faster using this reduction.

We describe here a third (and faster) way to derive the tip distribution. It is based on an analytical reduction of Equations (3a, 3b) that is specific to the PM model.

Remark that diag(*u*) and *^tr^uu* commute, leading to the following calculus,

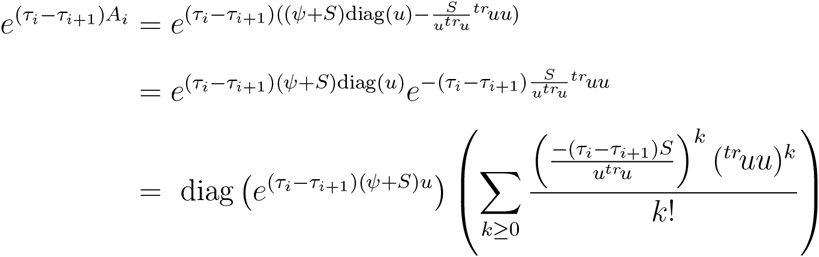

Where *e^w^* is the line vector with elements *e^wj^*. Further, remark that for any *k ≥* 1,

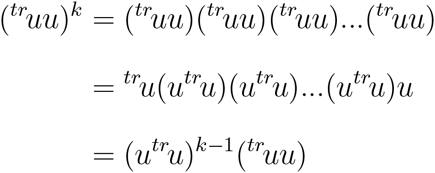

For simplicity, we will write in the following Δ = *τ_i_* − *τ*_*i*+1_, leading us to

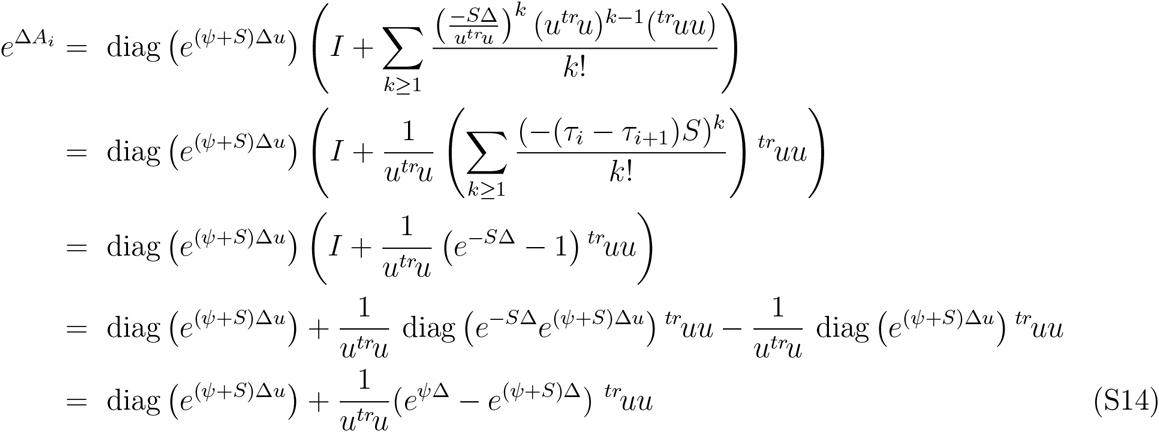

Where the last equality is due to the product by *^tr^u*, allowing to forget the cases where *u_j_* = 0 in the exponential.

We further need to compute

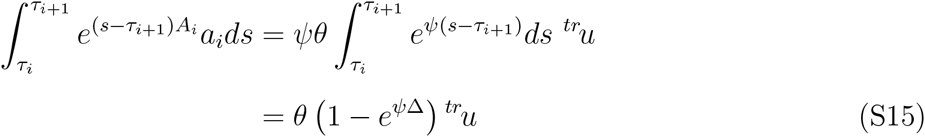

We thus get 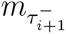 with the help of Equations (S14) and (S15).

Now, in order to simplify Equation (3b), remark that *A_i_* and Γ*_i_* are symmetric, and so are *e*^Δ*A_i_*^ and *e*^Δ*A_i_*^ Γ*_i_*. Moreover, Γ*_i_* is diagonal, and commutes with any other matrix, leading to,

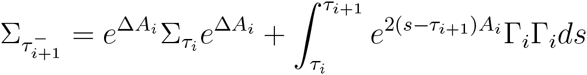

The first term can be computed thanks to Equation (S14). For the second one, remark that *^tr^uu* diag(*u*) = *^tr^uu*, thus leading to

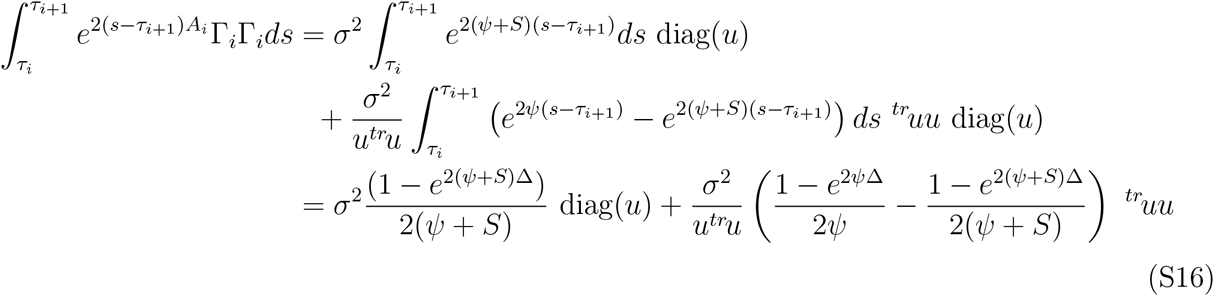

We thus get 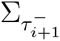 with the help of Equations (S14) and (S16).

#### C.3 The phenotype matching (PM) model with biogeography

In this section we describe ways to compute the tip distribution under the PM model, taking into account the biogeography (that is, species interact only when they co-occur in the same localities). We consider a fixed number of islands *N_I_*. Matrix *U* gives us the presence/absence of lineages in the distinct islands, with element *u_ij_* that equals 1 if lineage *j* is present on island *i* and zero otherwise. Vector *S* gives the strength of interaction on each island. The model states that the trait of lineage *j* evolves through phenotype matching with all species that are sympatric:

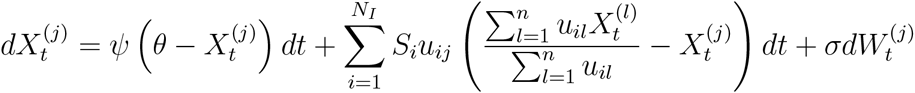

Take for example 5 lineages evolving on 3 distinct islands with the following *U* matrix on a given epoch:

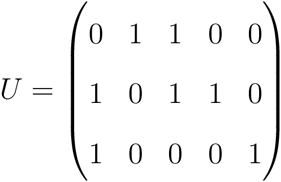

This means that species number 1 is present on island 2 and 3, species number 2 is only present on island 1, and so on… Said differently, we see that species number 3 interacts on island 1 with species 2, and on island 2 with species 1 and 4. Our species traits are driven by the following equations:

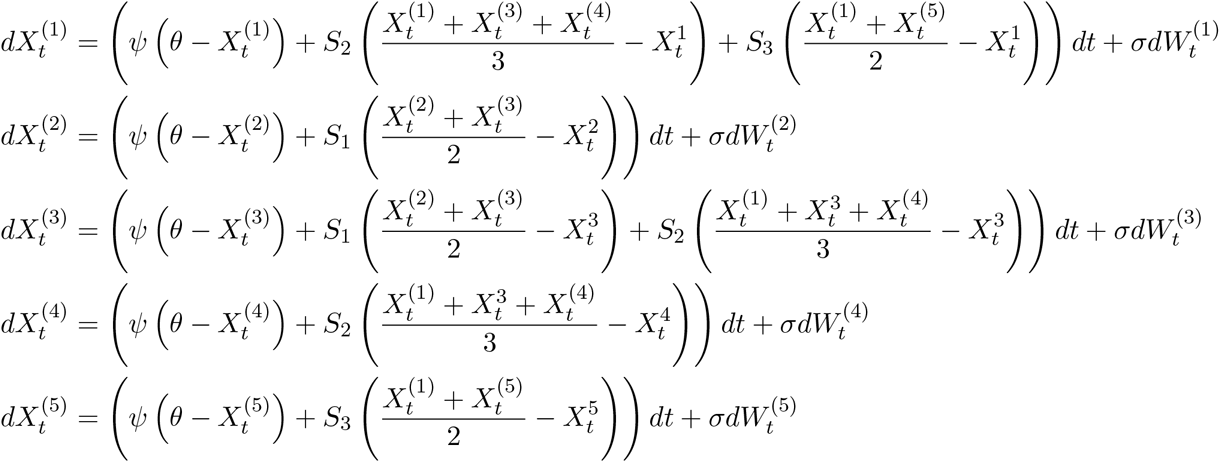

It thus follows that the vectorial equation can be written:

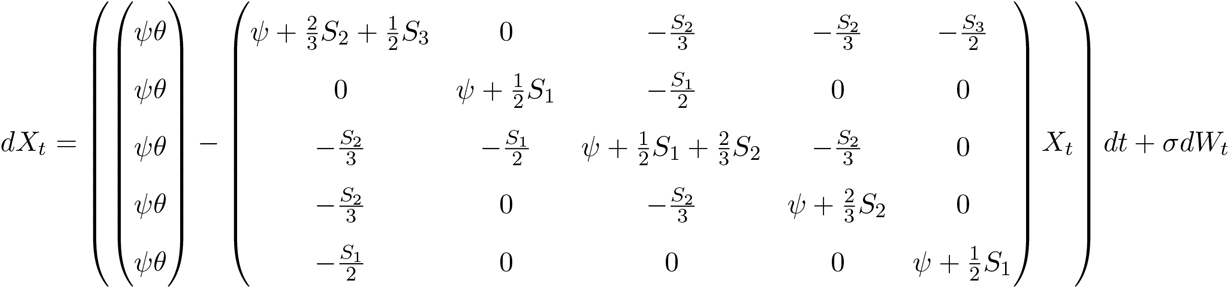

Provided no island is empty, the model can be written in our framework with *a* = *ψθV*, Γ = *σI*, and, finally, *A* which is the matrix with elements:

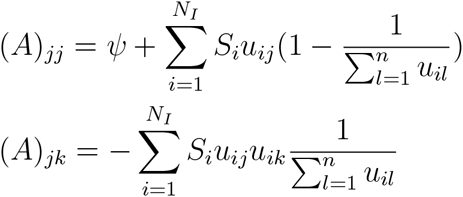

Matrix *A* is symmetric, and we can thus use the developments presented in Appendix C.1 to speed up the computation time.

Nonetheless, a better analytical reduction can be derived when islands are exclusive, meaning that species are allowed to occur on one island only. Under this assumption, matrix *U^T^U* is diagonal with element (*U^T^U)_ii_* being the number of lineages belonging to island *i*. We now introduce the line vector *r*, of size *N_I_*, full of ones. For simplicity, we also write in the following Δ = *τ_i_* − *τ*_*i*+1_. With these notations, and provided no island is empty, the model can be written under our framework with:

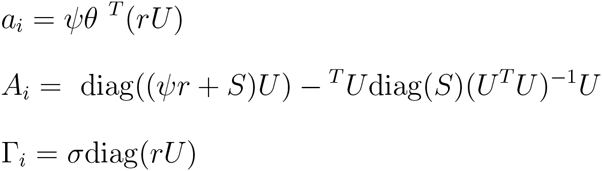

As for the one island case, we can speed up the computation of the exponential by remarking that:

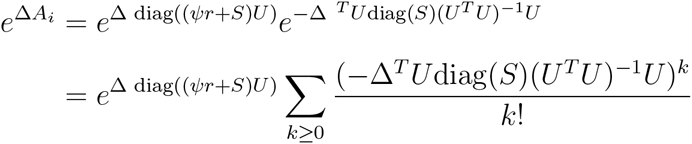

We then observe that:

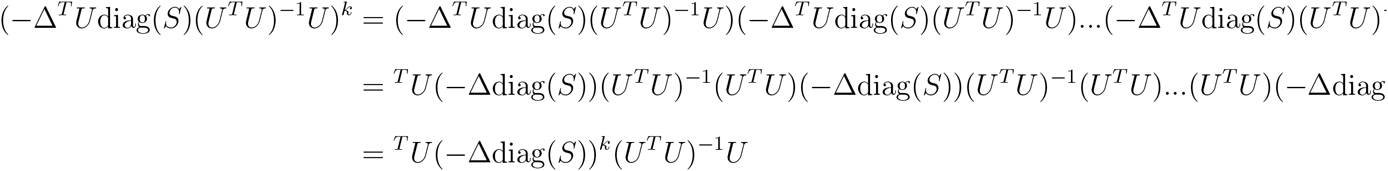

Thus leading to the following expression:

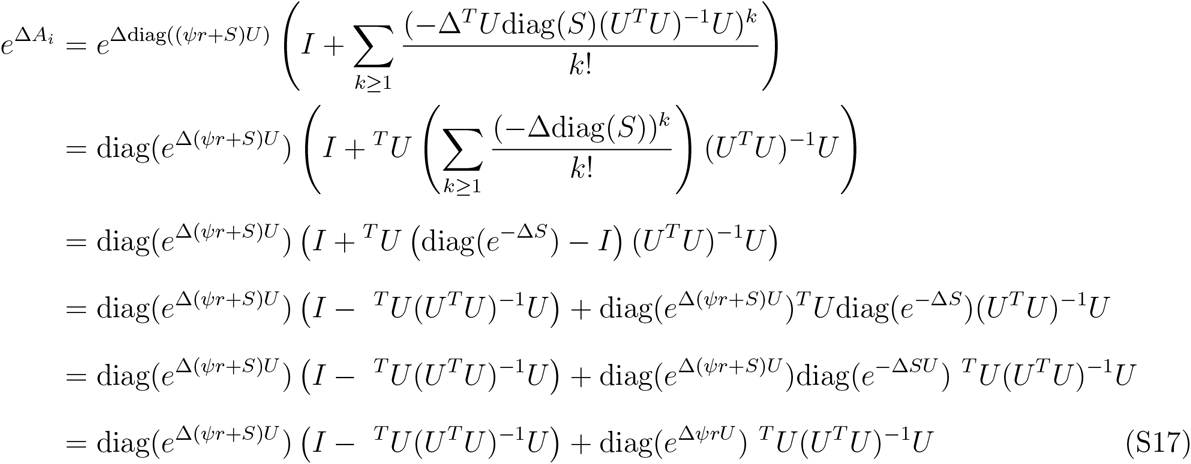

Where the second to last line holds under the assumption that each species belong to at most one island.

We further need to compute

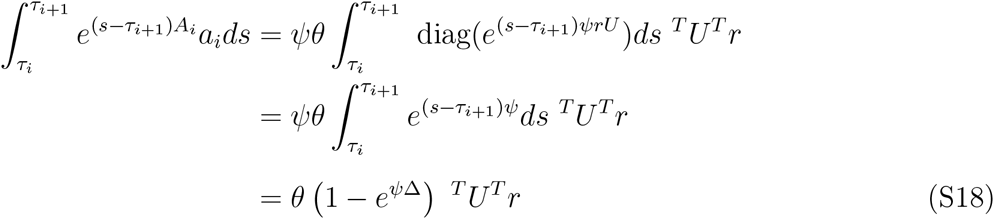

We thus get 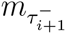 with the help of Equations (S17) and (S18).

We now turn to the reduction of the variance expression. Remark first that *A_i_* and Γ*_i_* are symmetric, and so are *e*^Δ*A_i_*^ and *e*^Δ*A_i_*^ Γ*_i_*. Moreover, Γ*_i_* is diagonal, and commutes with *e*^Δ*A_i_*^, leading to:

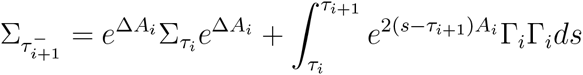

The first term can be computed thanks to equation (S17). For the second one we get

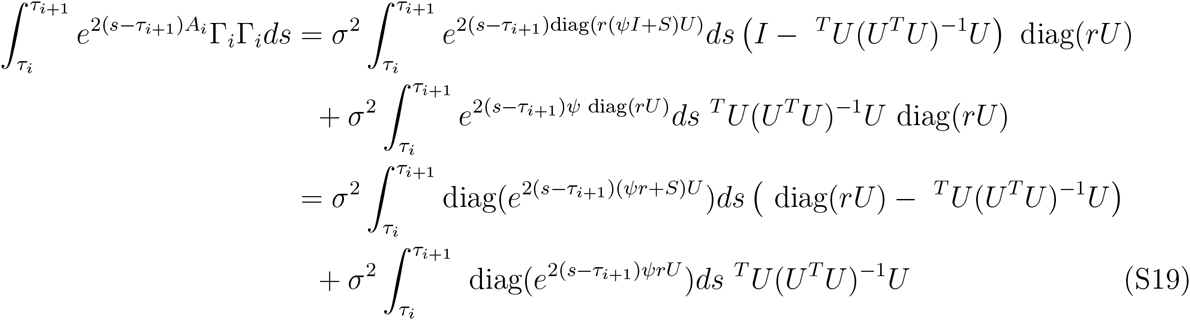

At the end, we get 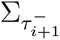 with the help of Equations (S17) and (S19).

#### C.4 The generalist matching mutualism (GMM) model

We recall the model formulation here. Assume that we rank first the *n*_1_ plant traits, before the *n*_2_ butterfly traits in the *X* vector. Traits evolve following the equation:

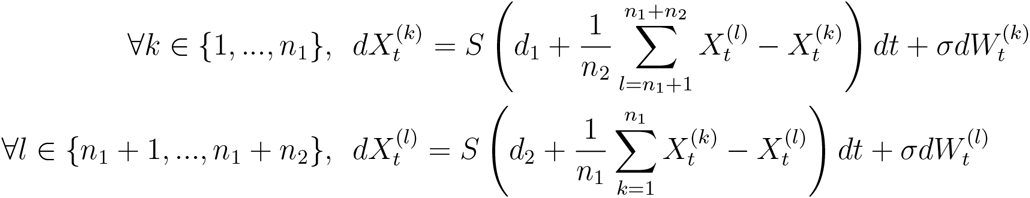

In the general framework formulation, this leads to:

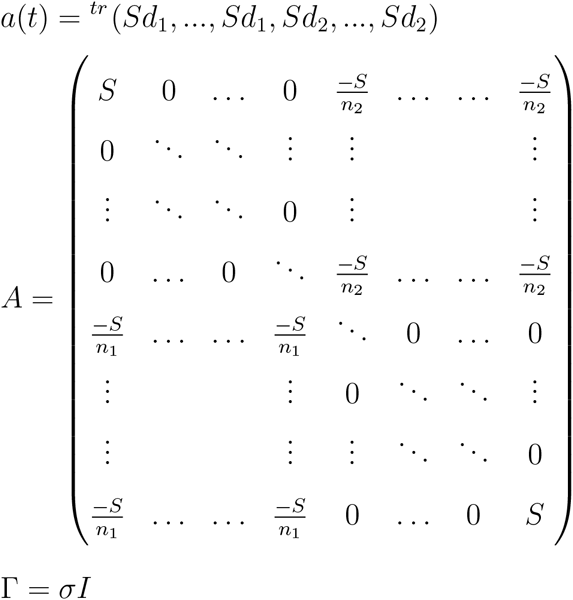

We would like to be able to compute the expectation and variance easily on each epoch. We thus want to reduce Equations (3a, 3b). For simplicity, we will write in the following Δ = *τ_i_ − τ_i_*_+1_. With some work, we can find the generic element of the matrix *e*^Δ*A*^.

First, we decompose *A* = *S*(*I* + *Z*), where *I* is the identity matrix, and *Z* is made of two blocks with elements 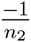 and 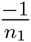. *I* and *Z* commute, meaning that:

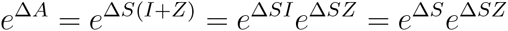

Moreover, we can find by induction the generic element of the matrix *Z^k^*, as presented in Figure (S2).

We then use this to find the generic element of the matrix 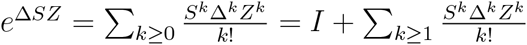. We recall that the odd and even parts of the exponential are:

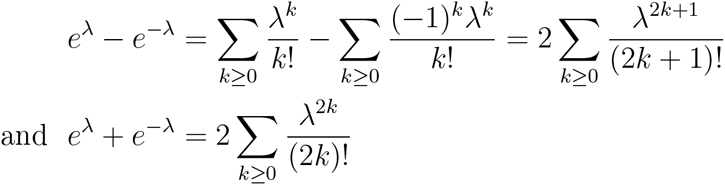

**Figure S2:**
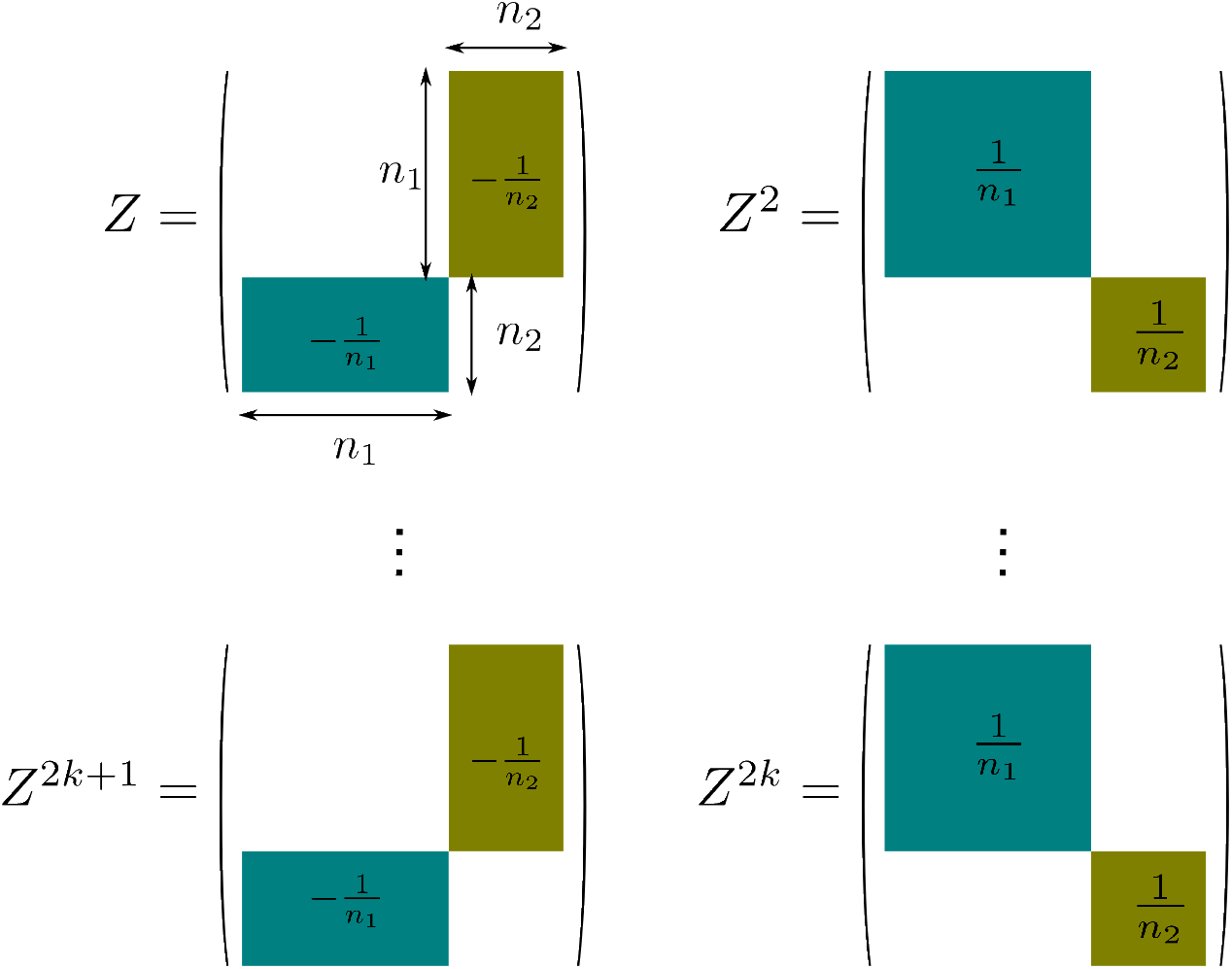
Generic element of the matrix 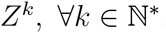.

Then, matrices *e*^Δ*SZ*^ and *e*^Δ*A*^ are composed of four distinct blocks, which expressions are shown in Figure S3.

**Figure S3:**
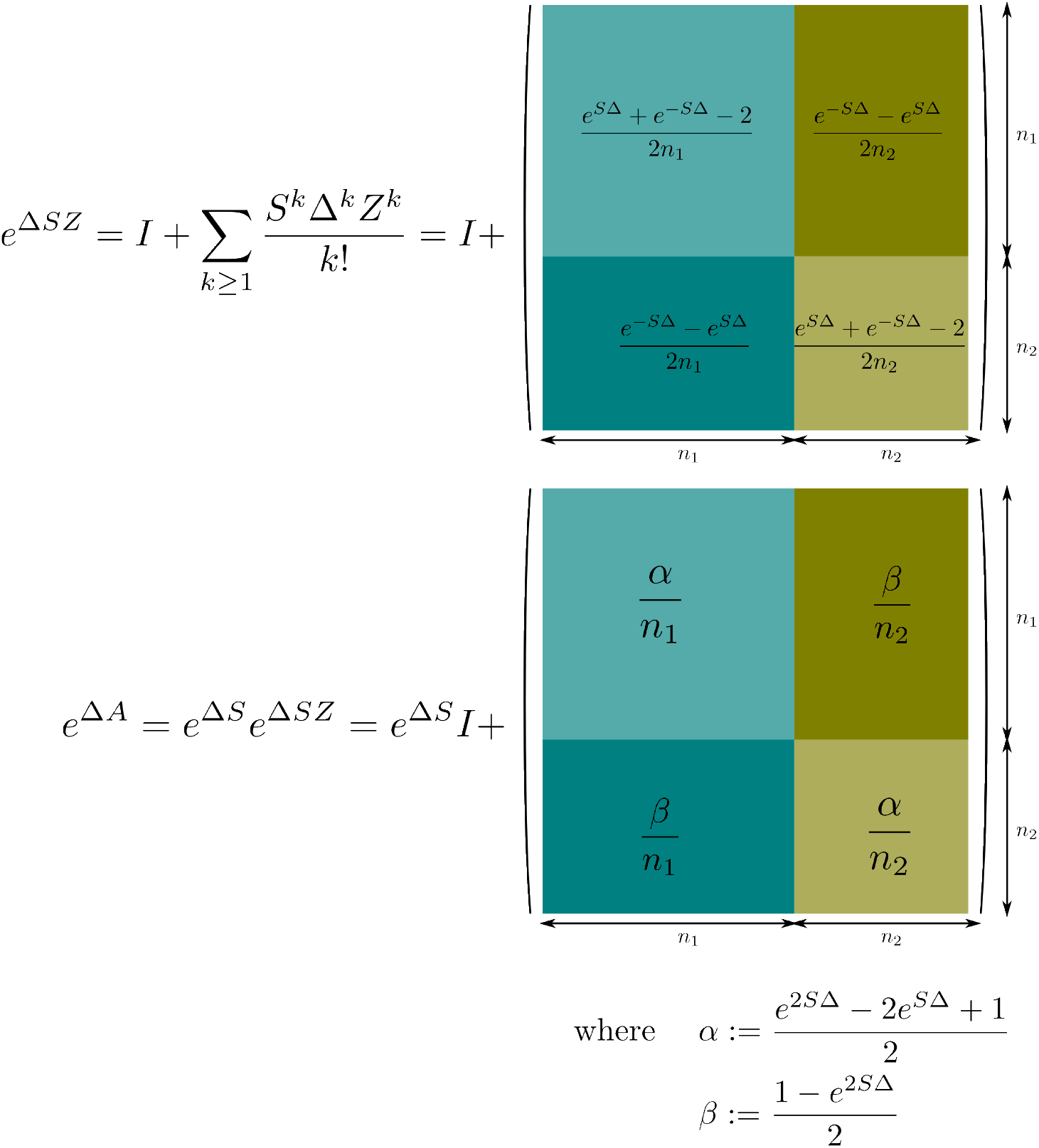
Generic elements of matrices *e*^Δ*SZ*^ and *e*^Δ*A*^.

We thus got the main element from which we can derive the expectation vector 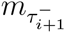:

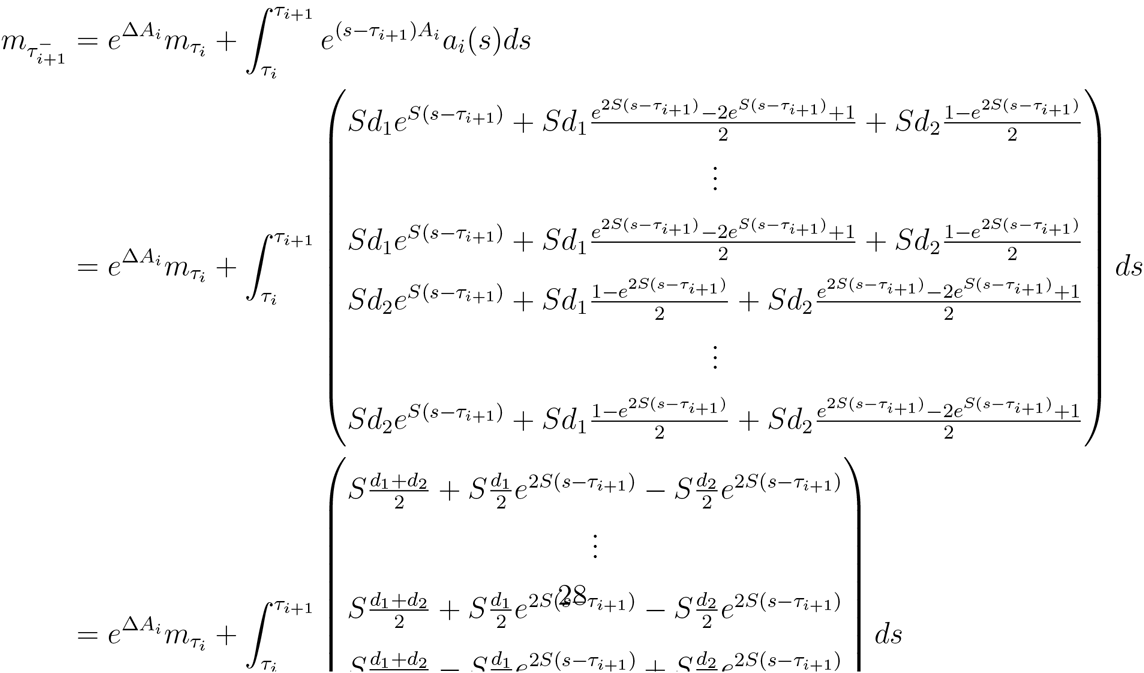

We now turn to the derivation of the covariance matrix, which requires simplifying:

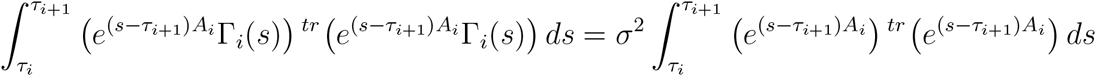

The expression of this last matrix is given in Figure S4.

**Figure S4:**
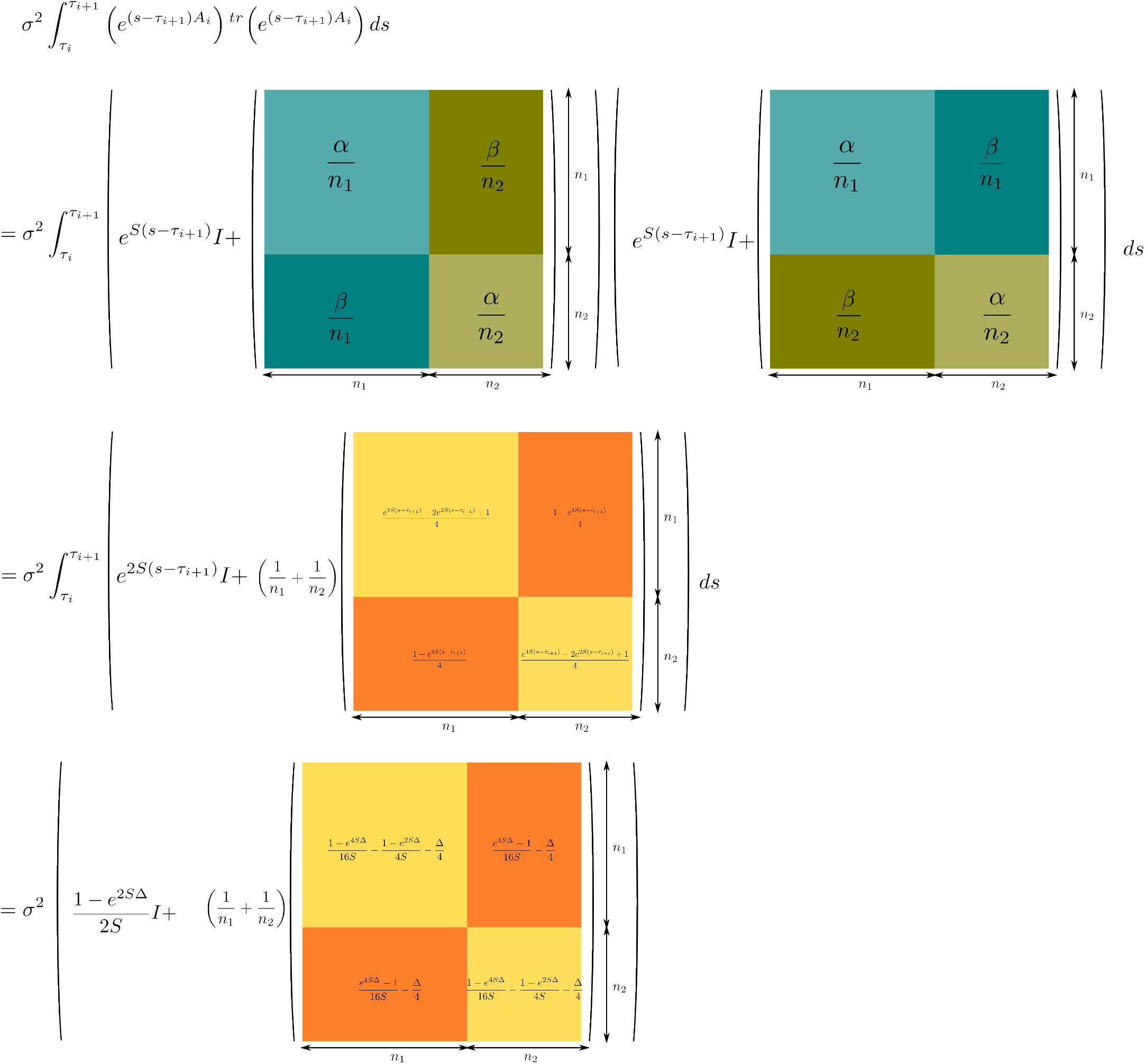
Generic elements of matrices that help us compute the covariance matrix of the distribution.

### D Simulation and Inference

We do not give any new result in this Appendix section. Instead, we present the ways we implemented numerically simulations and inferences for all models described in the paper. These have been previously described in a number of papers.

#### D.1 Numerical methods for simulating data

##### Simulating the whole trajectory of the process.—

We use the Euler-Maruyama scheme, which works like the Euler scheme for ODEs, but with the addition of a small Gaussian random variable at each time step (Gardiner et al. 1985). We discretize each epoch (*τ_i_, τ_i_*_+1_) with a mesh Δ*_t_*. We consider *m* standard Gaussian vectors of dimension *nd*: 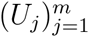. We approximate our SDE on this interval in the following way:

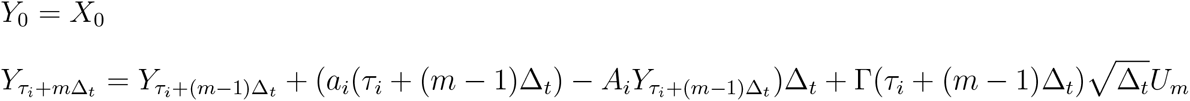

When a branching occurs, the values of the process on the splitting branch are duplicated at the end of the vector *Y*. We then iterate this operation from the root up to present time.

This simulation allows us to get the whole trajectory of the process on the tree, which can mainly be used to produce pictures as in Figure S5, and eventually get a useful intuition on the process. However, we rarely use the whole trajectories, because observed data are only composed of tip trait values.

##### Simulating values of the process at the tips only.—

This second simulation protocol allows us to simulate the process values at the tips only. Suppose that we know the vector *m* of expectations and the covariance matrix Σ at the tips of the tree.

We then simply simulate numerically a Gaussian vector with law:

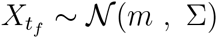

**Figure S5:**
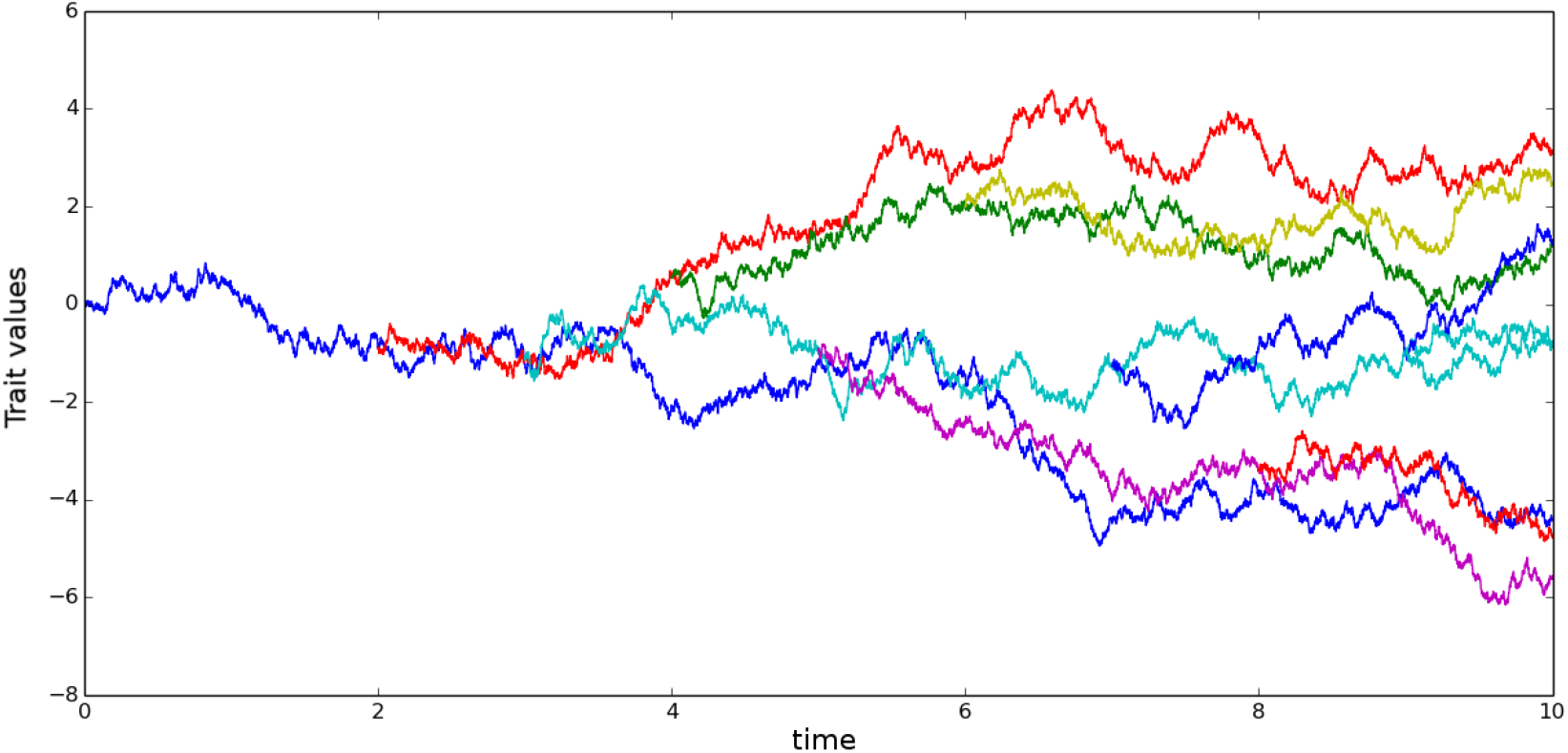
Evolution of a Brownian phenotypic trait along a tree, following the SDE: *dX_t_* = *σIdW_t_*.

This is by far the quickest way to get the tip values. However, as the inference protocol relies on the use of the same vector of expectations and covariance matrix, one may prefer to use the other simulation protocols to test the consistency between simulation and inference. In case there is an issue with the derivation of the tip distribution, there would be a discrepancy between simulations and inferences.

#### D.2 Parameter inference

##### Parameter inference principle.—

We consider here that we know the topology of the true phylogeny with *K* tips, its branch lengths, and the state of *d* phenotypic traits at the tip, denoted by 𝒳.

We assume any model of phenotypic evolution relying on linear SDEs, with vector of parameters *p*. We can compute the expectation *m_p_* and the covariance Σ*_p_* of the process *X* at tree tips. Its law is then: *X ∼ N* (*m_p_,* Σ*_p_*), and, assuming that the variance matrix is invertible, the density of the vector *X* is:

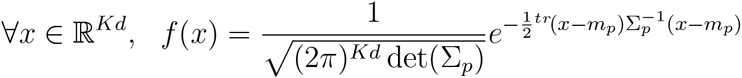

We can thus write the likelihood of the observed phenotypic traits as,

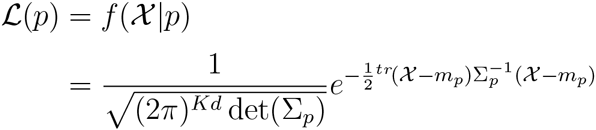

The maximum likelihood estimators (MLE) are the parameter values that maximize the likelihood function, that is,

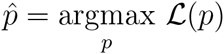

Equivalently, we can minimize the following function,

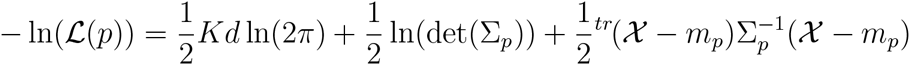

or, removing the constants,

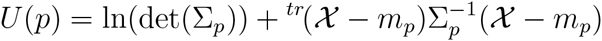

##### Analytical derivation of the MLE.—

Among all models described in the paper, only the BM model allows the analytic derivation of the MLE estimators. Take for illustration a BM model without drift starting with (*m*_0_, *v*_0_) = (0, 0). According to Table 1, the expectation *m* and covariance matrix Σ at the tips are *m* = 0 and Σ = *σ*^2^*T*, where matrix *T* has element *T* ^(*k,l*)^ = *t_k,l_*.

We get the MLE 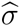 by looking analytically for the minimum of *U*,

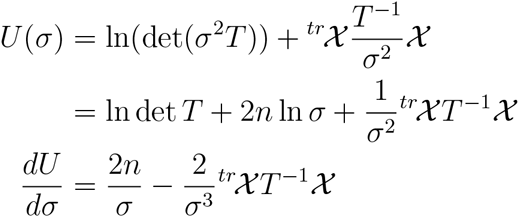

Thus leading to,

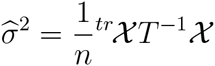

##### Speeding up the MLE estimation by reducing the dimension of the parameter space.—

Maximizing the likelihood can take a long time, especially when the dimension of the parameter space is large. It can thus be interesting to make assumptions that lower the number of parameters, when this is biologically tolerable. Examples include,

- starting an OU process with *m*_0_ = *θ*,
- considering no root variance, *v*_0_ = 0,
- starting a PM model with *m*_0_ = *θ* (in which case we easily show that the expectation remains *θ* in all lineages),
- putting *ψ* = 0 in the PM model.

In many models (e.g. BM, OU, ACDC, PM with *m*_0_ = *θ…*), distinct sets of parameters *p*_1_ and *p*_2_ are involved in the computation of *m* and Σ, and the expectation vector *m* can be expressed as *m* = *Cp*_1_. In this case, at a given *p*_2_, we can analytically get the parameters *p*_1_ maximizing ln(*L*(*p*_1_, *p*_2_)),

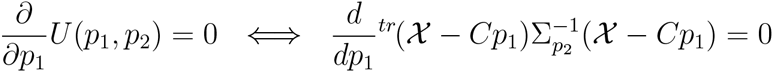

Doing so, we get the same formula as in (Hansen 1997; Butler and King 2004), i.e.:

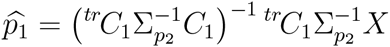

### E Tutorial: using the RPANDA codes to study trait coevolution

The aim of this section is to describe the R codes associated to our framework. We describe the class `PhenotypicModel`, we show how to manipulate the different methods included in the class, we illustrate their use around a simple (non-ultrametric) tree, and we finally explain how to use our codes to write new models fitting the framework.

We first need to load usefull R packages, along with our codes, and a small, non-ultrametric, tree.

~~~
In [219]: source(“Loading.R”)
          newick <- “((((A:1,B:0.5):2,(C:3,D:2.5):1):6,E:10.25):2,(F:6.5,G:8.25):3):1;”
          tree <- read.tree(text = newick)
          plot(tree)
~~~

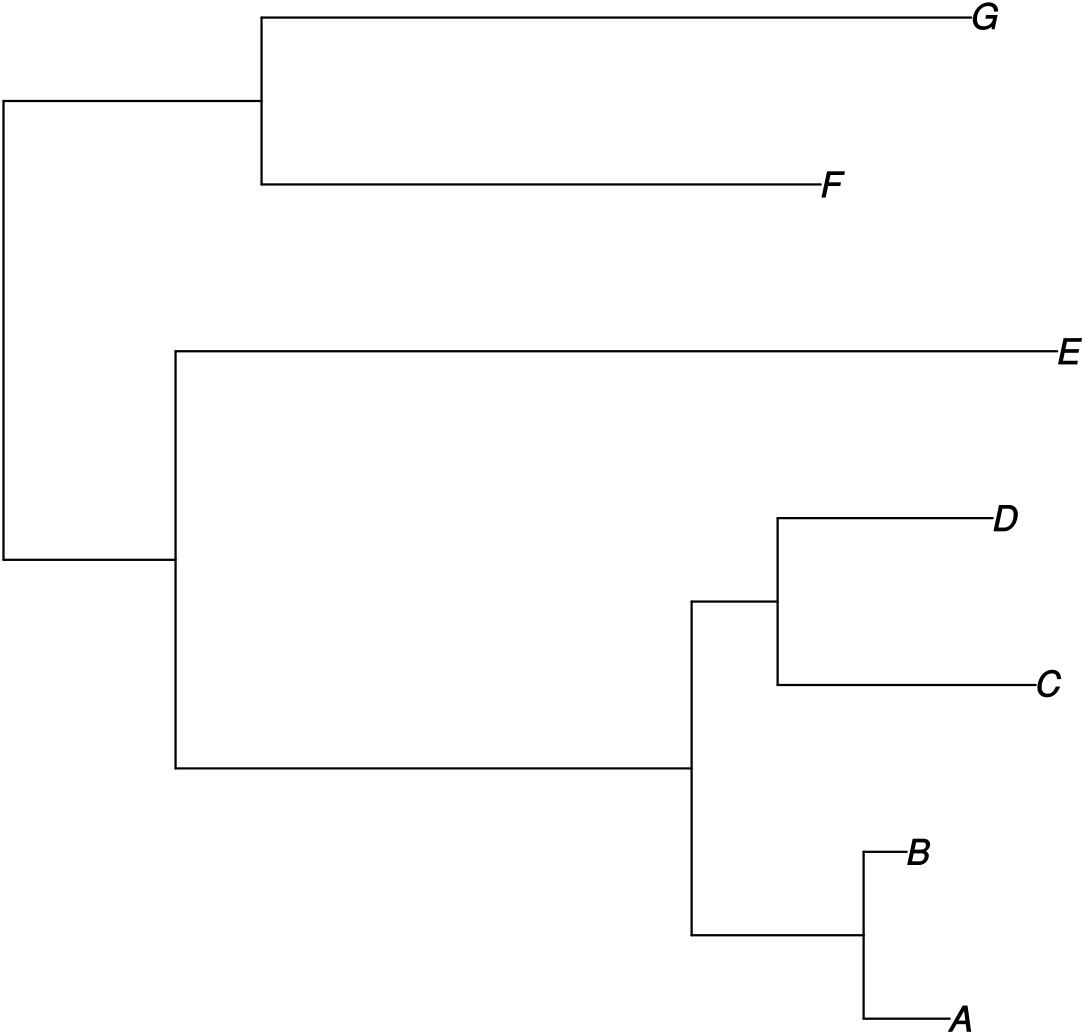

#### E.1 The ‘PhenotypicModel’ class

Our code is structured around one main R class that we called ‘PhenotypicModel’, which is intended to mimic the framework that we proposed in the main text. Each object of the ‘PhenotypicModel’ encompasses informations on the tree, on the parameters of the model, on the starting values, and, finally, on the collection of (*a_i_, A_i_*, Γ*_i_*) for all epochs.

##### Loading a pre-defined model.—

Because we wanted this code both to be user-friendly and to serve as an illustration of what can be written within this framework, we implemented all models in main Table 1 in a generic constructor `createModel`, in the file ‘ModelBank.R’, that takes for arguments the tree and the name of the required model.

Available models include:

**BM** Brownian Motion model with linear drift.

Starts with two lineages having the same value *X*_0_ ∼ 𝒩 (*m*_0_, *v*_0_).
One trait in each lineage, all lineages evolving independently after branching following the equation.

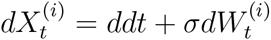

**BM_from0** Same as above, but starting with two lineages having the same value *X*_0_ ∼ 𝒩 (0, 0).

**BM_from0_driftless** Same as above, but with *d* = 0.

**OU** Ornstein-Uhlenbeck model.

Starts with two lineages having the same value *X*_0_ ∼ 𝒩 (*m*_0_, *v*_0_).
One trait in each lineage, all lineages evolving independently after branching, following the equation:

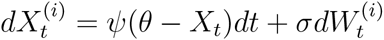

**OU_from0** Same as above, but starting with two lineages having the same value *X*_0_ ∼ 𝒩 (0, 0).

**ACDC** ACcelerating or DeCelerating model.

Starts with two lineages having the same value *X*_0_ ∼ 𝒩 (*m*_0_, *v*_0_).
One trait in each lineage, all lineages evolving independently after branching, following the equation:

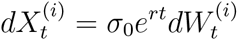

**DD** Diversity-Dependent model.

Starts with two lineages having the same value *X*_0_ ∼ 𝒩 (*m*_0_, *v*_0_).
One trait in each lineage, all lineages evolving independently after branching, following the equation:

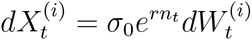

**PM** Phenotype Matching model.

Starts with two lineages having the same value *X*_0_ ∼ 𝒩 (*m*_0_, *v*_0_).
One trait in each lineage, all lineages evolving then non-independently following the expression:

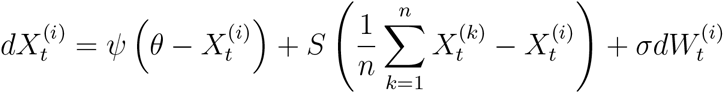

**PM_OUless** Simplified Phenotype Matching model.

Starts with two lineages having the same value *X*_0_ ∼ 𝒩 (*m*_0_, *v*_0_).
One trait in each lineage, all lineages evolving then non-independently following the expression:

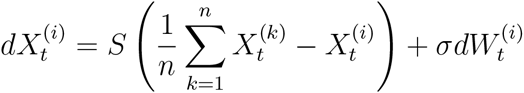

To get a first glimpse at ‘PhenotypicModel’ objects, we first create two such objects. The first one is a Brownian Motion (BM), the second one is an Ornstein-Uhlenbeck process (OU). Note that both models include *m*_0_ and *v*_0_ as parameters.

~~~
In [220]: modelBM <- createModel(tree, ‘BM’)
          modelOU <- createModel(tree, ‘OU’)
~~~

##### Access to the content of the model.—

The function `show` (resp. `print`) is intended to give basic (resp. full) information on a specific ‘PhenotypicModel’ object.

~~~
In [221]: show(modelBM)
****************************************************************
*** Object of Class PhenotypicModel ***
*** Name of the model: [1] “BM”
*** Parameters of the model: [1] “m0” “v0” “d” “sigma”
*** Description: Brownian Motion model with linear drift.
Starts with two lineages having the same value X_0 ∼ Normal(m0,v0).
One trait in each lineage, all lineages evolving independently after branching.
dX_t = d dt + sigma dW_t
*** Periods: the model is cut into 13 parts.
For more details on the model, call: print(PhenotypicModel)
****************************************************************
In [222]: print(modelOU)
****************************************************************
*** Object of Class PhenotypicModel ***
*** Name of the model: [1] “OU”
*** Parameters of the model: [1] “m0” “v0” “psi” “theta” “sigma”
*** Description: Ornstein-Uhlenbeck model.
Starts with two lineages having the same value X_0 ∼ Normal(m0,v0).
One trait in each lineage, all lineages evolving independently after branching.
dX_t = psi(theta- X_t) dt + sigma dW_t
*** Epochs: the model is cut into 13 parts.
[1] 0.00 2.00 3.00 8.00 9.00 9.50 10.00 10.50 11.00 11.25 11.50 12.00
[13] 12.25
*** Lineages branching (to be copied at the end of the corresponding period):
[1] 1 1 2 1 5 2 1 7 1 4 6 5 3
*** Positions of the new trait at the end of each period:
[1] 2 3 4 5 6 0 7 0 0 0 0 0 0
*** Initial condition: function (params)
return(list(mean = c(params[1]), var = matrix(c(params[2]))))
<environment: 0x9617460>
*** Vectors a_i, A_i, Gamma_i on each period i: function (i, params)
{
     vectorU <- getLivingLineages(i, eventEndOfPeriods)
     vectorA <- function(t) return(params[3] * params[4] * vectorU)
     matrixGamma <- function(t) return(params[5] * diag(vectorU))
     matrixA <- params[3] * diag(vectorU)
     return(list(a = vectorA, A = matrixA, Gamma = matrixGamma))
}
<environment: 0x9617460>
*** Constraints on the parameters:
function (params)
return(params[2] >= 0 && params[5] >= 0 && params[3] ! = 0)
<environment: 0 × 9617460>
*** Defaut parameter values: [1] 0 0 1 0 1
*** Tip labels:
[1] “A” “B” “C” “D” “E” “F” “G”
*** Tip labels for simulations:
[1] “A” “F” “E” “G” “C” “D” “B”
****************************************************************
~~~

##### List of class attributes.—

The latter command gave us some insight into how a PhenotypicModel is defined. It has the following list of attributes:

**name** a name,

**paramsNames** the names of all parameters,

**comment** a short description,

**period** the vector of times at which successive branching and death of lineages occur,

**numbersCopy** vector containing the lineage number which branches or dies at the end of each period,

**numbersPaste** vector containing the lineage number in which a daughter lineage is placed at the end of each period (zero if the end of the period corresponds to a death),

**initialCondition** a function of the parameters giving the initial mean and variance of the gaussian process at the root of the tree,

**aAGamma** the functions corresponding to *a_i_*(*t*), *A_i_*, and Γ*_i_*(*t*) that define the evolution of the process on each period, depending on parameters,

**constraints** a function of the parameters giving the definition range,

**params0** a vector of defaut parameter values.

Each of these attributes can be accessed and changed through the use of the following syntax.

~~~
In [223]: modelBM[’name’]
Out[223]: ‘BM’
In [224]: modelBM[’paramsNames’]
Out[224]: ‘m0’ ‘v0’ ‘d’ ‘sigma’
In [225]: modelOU[’paramsNames’] <- c(“mean0”, “var0”, “selectionStrength”, “equilibrium”, “noise”)
          show(modelOU)
****************************************************************
*** Object of Class PhenotypicModel ***
*** Name of the model: [1] “OU”
*** Parameters of the model: [1] “mean0” “var0” “selectionStrength”
[4] “equilibrium” “noise”
*** Description: Ornstein-Uhlenbeck model.
Starts with two lineages having the same value X_0 ∼ Normal(m0,v0).
One trait in each lineage, all lineages evolving independently after branching.
dX_t = psi(theta- X_t) dt + sigma dW_t
*** Periods: the model is cut into 13 parts.
For more details on the model, call: print(PhenotypicModel)
****************************************************************
~~~

However, changes must be made cautiously, in order to keep a coherent model. For example, changing ‘paramsNames’ for a shorter vector would not be authorized, but other deleterious actions could work and lead to issues with methods associated to `PhenotypicModel` objects.

~~~
In [226]: modelOU[’paramsNames’] <- c(“mean0”, “var0”)
Error in validityMethod(as(object, superClass)): [PhenotypicModel: validation]
There should be the same number of defaut parameters and parameter names.
~~~

#### E.2 Methods associated to the ‘PhenotypicModel’ class

All ‘PhenotypicModel’ objects are associated to methods intended to do the basic operations that we need to do with models of trait evolution, i.e.,

1. simulate tip trait data,
2. compute the likelihood of tip trait data,
3. fit the model to tip trait data.

##### Simulating tip trait data.—

The method `simulateTipData` works for any `PhenotypicModel` object. We simply give it the model and the set of parameters and it returns a realisation of the process (tip data).

~~~
In [227]: dataBM <- simulateTipData(modelBM, c(0,0,0,1))
             dataBM
*** Simulation of tip trait values ***
Simulates step-by-step the whole trajectory, but returns only the tip data.
Computation time: 0.3909395 secs
Out[227]:
~~~

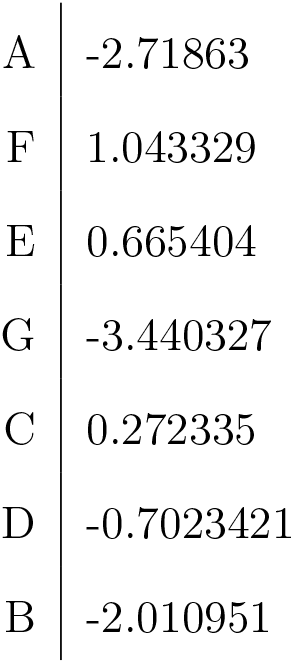

A third, optional, argument, changes the behaviour of the method.

- “method = 1”: first computes the tip distribution at present, before drawing a realization of this distribution,
- “method = 2”: simulates step-by-step the whole trajectory of the process, plots the trajectories through time, and returns the tip data.
- “method = 3”: (default) simulates step-by-step the whole trajectory of the process, before returning only the tip data.

~~~
In [228]: dataOU <- simulateTipData(modelOU, c(0,0,1,5,1), method = 1)
          dataOU
*** Simulation of tip trait values ***
Computes the tip distribution, and returns a simulated dataset drawn in this distribution.
Computation time: 0.0009741783 secs
Out[228]:
~~~

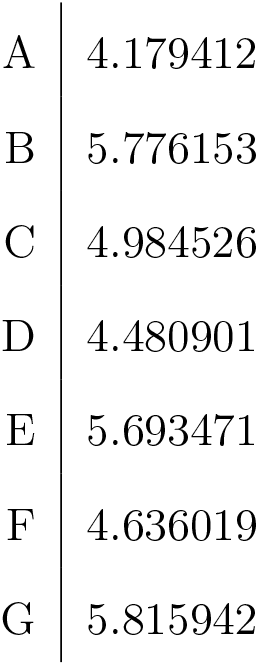

~~~
In [229]: simulateTipData(modelBM, c(0,0,0,1), method = 2)
*** Simulation of tip trait values ***
Simulates step-by-step the whole trajectory, plots it, and returns tip data.
Computation time: 0.479032 secs
Out[229]:
~~~

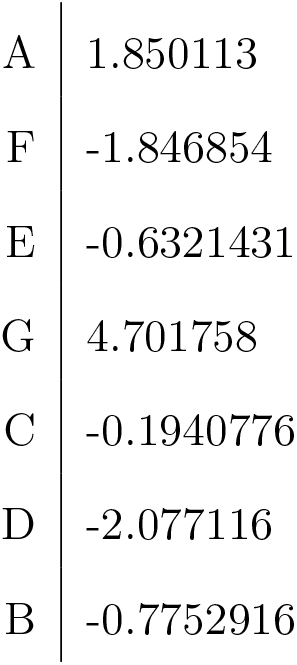

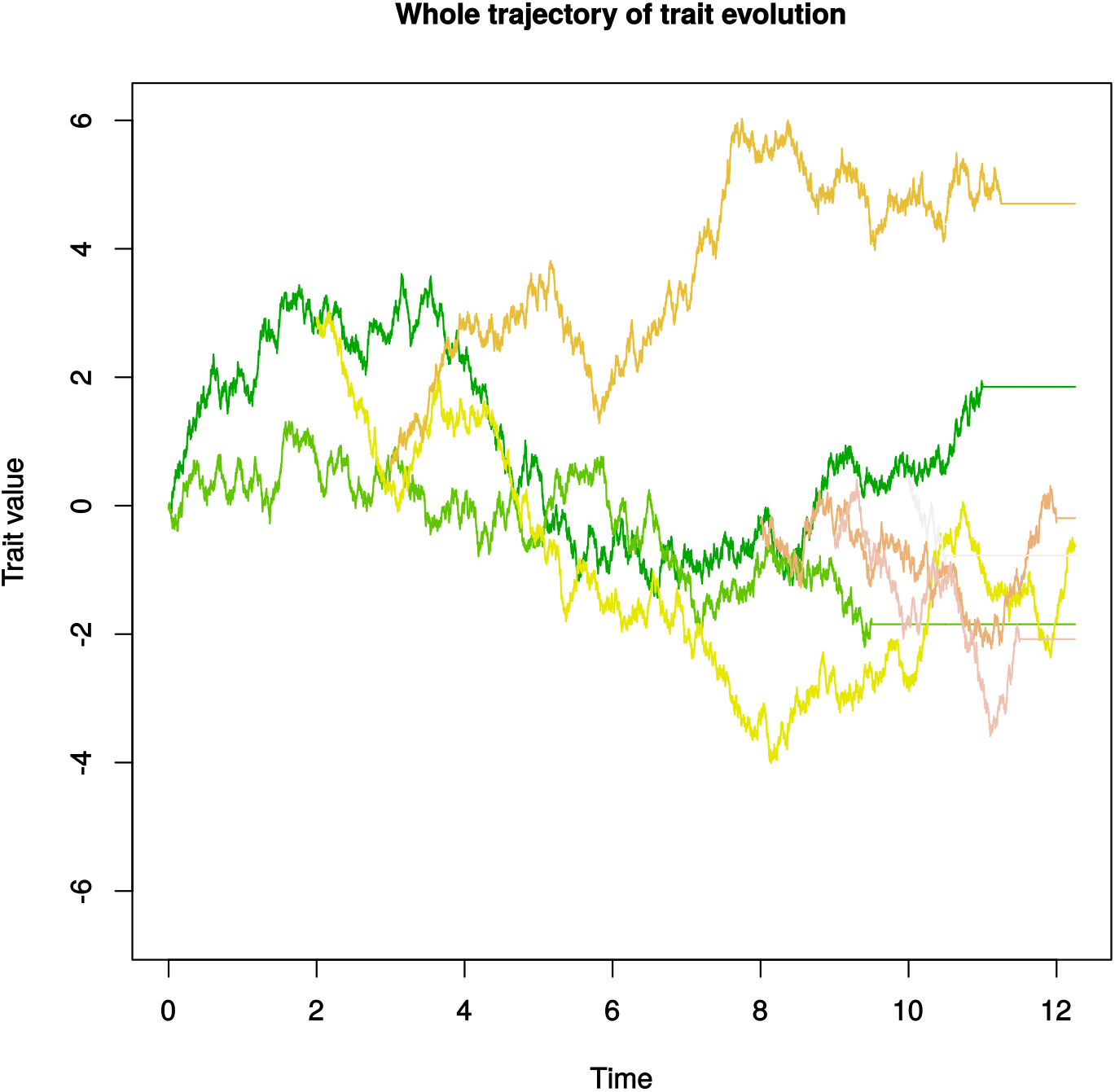

##### Getting the distribution of the model under a given set of parameters.—

The method `getTipDistribution` computes the mean vector *m* and variance matrix Σ such that, under the model, the tip trait data *X* follows 𝒩 (*m,* Σ).

The related method `getDataLikelihood` returns the -ln(likelihood) of a given data set under the model, with a given set of parameters.

~~~
In [230]: getTipDistribution(modelBM, c(0,0,1,1))
Out[230]:
~~~

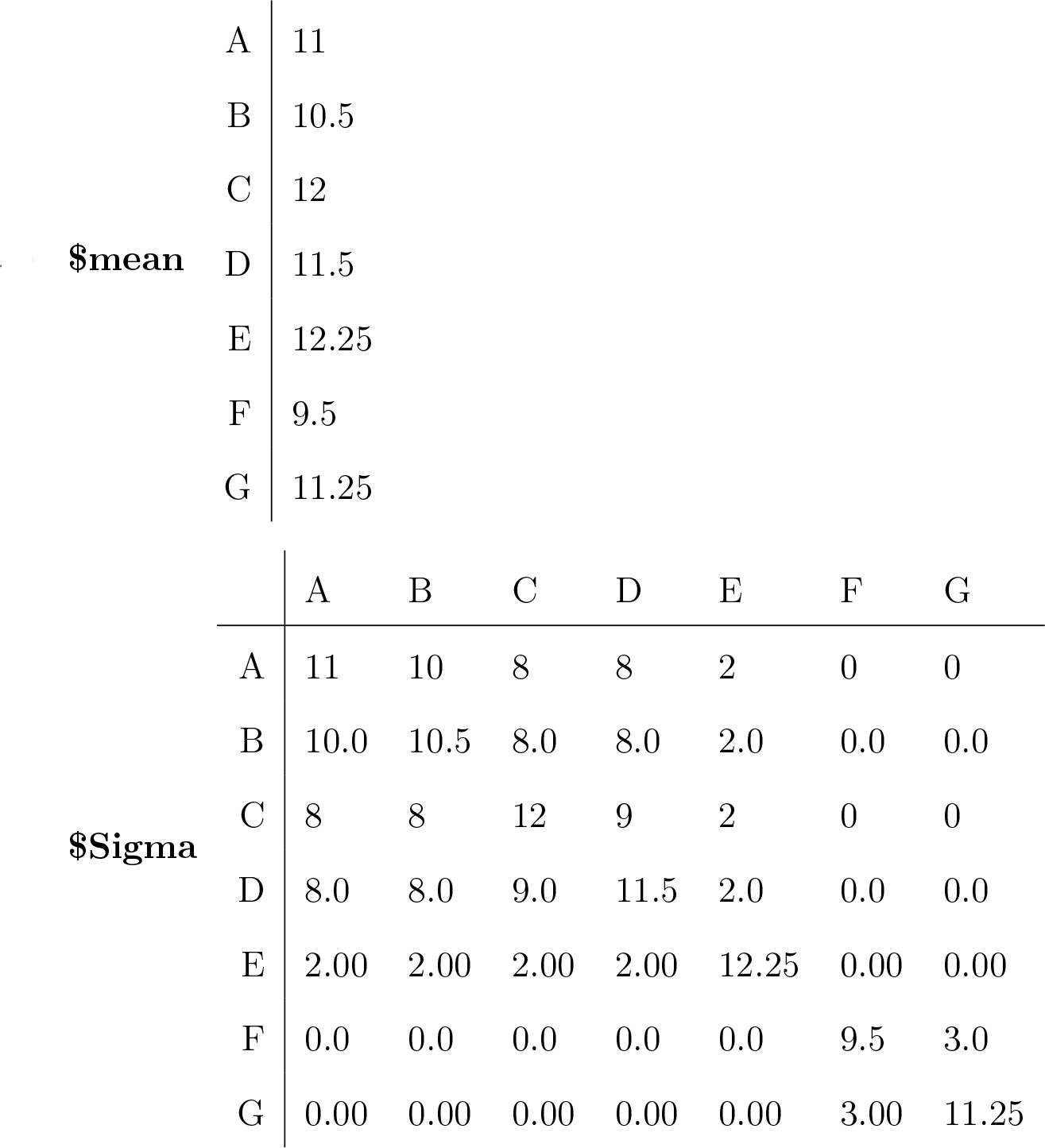

~~~
In [231]: getDataLikelihood(modelBM, dataBM, c(0,0,1,1))
Out[231]: 36.0510113479088
~~~

##### Maximum likelihood estimation of parameters.—

The method `fitTipData` uses the latter two methods to find the set of parameters that minimizes -ln(likelihood) for a given model, on a given data set. We can apply this method to simulated datasets, and compare the maximum likelihood estimators with the parameters used in the simulation.

Note that this function accepts a third, optional, parameter, that is the starting vector ‘params0’ given to optimize the likelihood. If no value is specified, the function takes the attribute ‘params0’ in the `PhenotypicModel` object.

~~~
In [232]: fitTipData(modelBM, dataBM)
*** Fit of tip trait data ***
Finds the maximum likelihood estimators of the parameters,
returns the likelihood and the inferred parameters.
Computation time: 0.02105212 secs
Out[232]:
**$value** 13.3539168672421
**$inferredParams m0** 0.112360024529455
          **v0** 4.3703974585017e-08
          **d** -0.0733871266399529
          **sigma** 0.64761762031608
In [233]: fitTipData(modelOU, dataOU)
*** Fit of tip trait data ***
Finds the maximum likelihood estimators of the parameters,
returns the likelihood and the inferred parameters.
Computation time: 0.2915776 secs
Out[233]:
**$value** 7.5162883379935
**$inferredParams mean0** 13.5665180225751
          **var0** 1.6815664554916e-05
          **selectionStrength** 0.648513938633288
          **equilibrium** 5.05532921748184
          **noise** 0.766630199120977
~~~

It doesn't seem quite good, but it also seems like the choice in the starting parameters *m*_0_, *v*_0_ has a bad influence. As presented in Appendix D.2, in many models (e.g. BM, OU, ACDC, PM with *m*_0_ = *θ*…), distinct sets of parameters *p*_1_ and *p*_2_ are involved in the computation of *m* and Σ, and the expectation vector *m* can be expressed as *m* = *C*_*p*1_. In particular, many models verify *m* = *^tr^*(*m*_0_, *m*_0_, *…m*_0_). When this is the case, the fit of tip data can be improved and speeded up by using the third parameter of the function `GLSstyle=TRUE`.

~~~
In [234]: fitTipData(modelBM, dataBM, GLSstyle = TRUE)
          fitTipData(modelOU, dataOU, GLSstyle = TRUE)
*** Fit of tip trait data ***
Finds the maximum likelihood estimators of the parameters,
returns the likelihood and the inferred parameters.
Computation time: 0.03260899 secs
Out[234]:
**$value** 13.5302740469078
**$inferredParams m0** -0.00550320295296933
         **v0** 2.28469756397133e-07
         **d** -0.313019528308928
         **sigma** 0.663621107698308
*** Fit of tip trait data ***
Finds the maximum likelihood estimators of the parameters,
returns the likelihood and the inferred parameters.
Computation time: 0.1760004 secs
Out[234]:
**$value** 7.82305350777471
**$inferredParams mean0** 5.10957361631891
         **Var0** 3.36222531349288e-05
         **selectionStrength** 1.87722870245168
         **equilibrium** -1.98889519193151
         **noise** 1.91905948952067
~~~

With so few data in hand, we could also prefer to consider directly models starting with (*m*_0_, *v*_0_) = (0, 0). We create two new models ‘BM_from0’ and ‘OU_from0’ with the subtle difference that (*m*_0_, *v*_0_) = (0, 0) and the models thus retain respectively only two and three parameters.

These two models are included in the ‘ModelBank’ file.

~~~
In [235]: modelBMfromZero <- createModel(tree, ‘BM_from0’)
          modelBMfromZero[’paramsNames’]
Out[235]: ‘d’ ‘sigma’
In [236]: modelOUfromZero <- createModel(tree, ‘OU_from0’)
          modelOUfromZero[’paramsNames’]
Out[236]: ‘psi’ ‘theta’ ‘sigma’
In [237]: fitTipData(modelBMfromZero, dataBM)
          fitTipData(modelOUfromZero, dataOU)
*** Fit of tip trait data ***
Finds the maximum likelihood estimators of the parameters,
returns the likelihood and the inferred parameters.
Computation time: 0.01061678 secs
Out[237]:
**$value** 13.3540474589618
**$inferredParams d** -0.0633929373190768
          **sigma** 0.647501517840828
*** Fit of tip trait data ***
Finds the maximum likelihood estimators of the parameters,
returns the likelihood and the inferred parameters.
Computation time: 0.3246026 secs
Out[237]:
**$value** 12.8406380062571
**$inferredParams psi** -0.00173618484753718
         **theta** -257.683877940727
         **sigma** 0.598311115102137
~~~

While the first inference seems quite consistent, the second one is obviously wrong.

We would like here to warn users about the use of the `fitTipData` method. We did not code an appropriate optimizer, and we use instead **optim**, the optimizer available in R, which sometimes seems to be attracted to a wrong region in the parameter space. Starting the optimization with different parameter sets might be the best practice to comfort the results.

For example, here, starting with another initial parameter set leads to a better likelihood optimization.

~~~
In [238]: getDataLikelihood(modelOUfromZero, dataOU, c(1,5,1))
Out[238]: 7.17243577660605
In [239]: fitTipData(modelOUfromZero, dataOU, c(1,5,1))
*** Fit of tip trait data ***
Finds the maximum likelihood estimators of the parameters,
returns the likelihood and the inferred parameters.
Computation time: 0.188132 secs
Out[239]:
**$value** 6.69894523491705
**$inferredParams psi** 9.38482325085794
         **theta** 5.08087953439187
         **sigma** 2.72996276707977
~~~

Finally, the functions `getTipDistribution, simulateTipData` and `fitTipData` all have a last optional argument, called v for “verbose mode”. With `v=TRUE`, the functions gives informations in the console, whereas with `v=FALSE` the function remains silent.

#### E.3 Toward an in-depth understanding of the code structure

This section can be skipped if you are not interested in using this framework to build your own model. Otherwise, it is worth understanding how the different models relate to each others.

##### Relationships between the different classes of models.—

The mother and most general class, for which all the above-mentionned functions are defined, is the `PhenotypicModel` class. When a model is only known as a `PhenotypicModel`, the method that computes the tip distribution, namely `getTipDistribution` is the most general one. It thus computes the distribution by resolving numerically the ODE system presented in main text Equations (4a, 4b), which can take a lot of time.

However, faster algorithms are available to compute the tip distribution under specific models (see e.g. analytical tip distribution formulas in Table S1). This is the rationale to define daughter-classes:

**PhenotypicBM** For the Brownian model.

**PhenotypicOU** For the Ornstein-Uhlenbeck model.

**PhenotypicACDC** For the Accelerating/Decelerating model.

**PhenotypicDD** For the Diversity-Dependent model.

**PhenotypicPM** For the Phenotype-Matching model.

**PhenotypicGMM** For the Generalist Matching Mutualism model.

**PhenotypicADiag** Models for which, *∀i, A_i_* is symmetric and Γ*_i_* = *σI*.

For each of these daughter-classes, an other, more appropriated, function `getTipDistribution` has been written. `PhenotypicModels` which are also `PhenotypicOU`, will preferentially use methods defined for PhenotypicOU when they exist.

##### Application: three different ways to define an OU.—

In the `createModel` function, the keyword ‘OU’ constructs a model in the class `PhenotypicOU`. In this class, the function `getTipDistribution` uses the analytical formula show in Appendix B.1 to speed up the computation of *m* and Σ.

Alternatively, the keyword ‘OUbis’ defines the exact same model, but as an instance of the class `PhenotypicADiag`. Thus, the function `getTipDistribution` uses the reduction show in Appendix C.1 to compute *m* and Σ.

Last, the keyword ‘OUter’ still defines the exact same model, but as an instance of the class `PhenotypicModel`. Thus, the function `getTipDistribution` uses the resolution of the ODE system to compute *m* and Σ.

The following lines of code show that the function returns the same value with the three different methods, but do not take the same amount of time.

~~~
In [240]: modelOU <- createModel(tree, ‘OU’)
          modelOUbis <- createModel(tree, ‘OUbis’)
          modelOUter <- createModel(tree, ‘OUter’)
          params <- c(0,0,0.2,1,2)
In [241]: getTipDistribution(modelOU, params, v = TRUE)
*** Computation of tip traits distribution through the analytical formula for an OU process ***
Computation time: 0.000497818 secs
Out[241]:
~~~

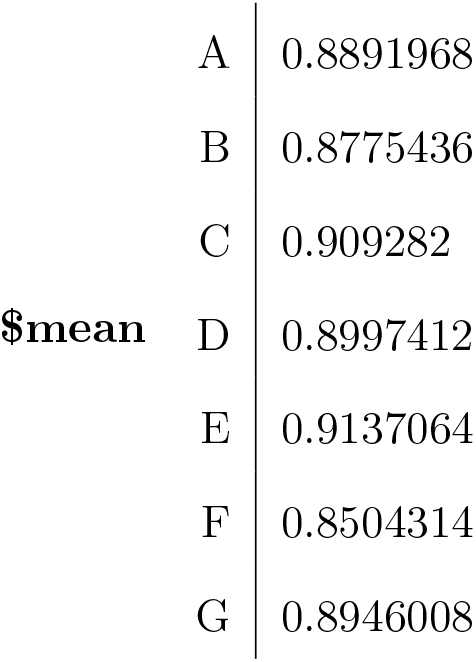

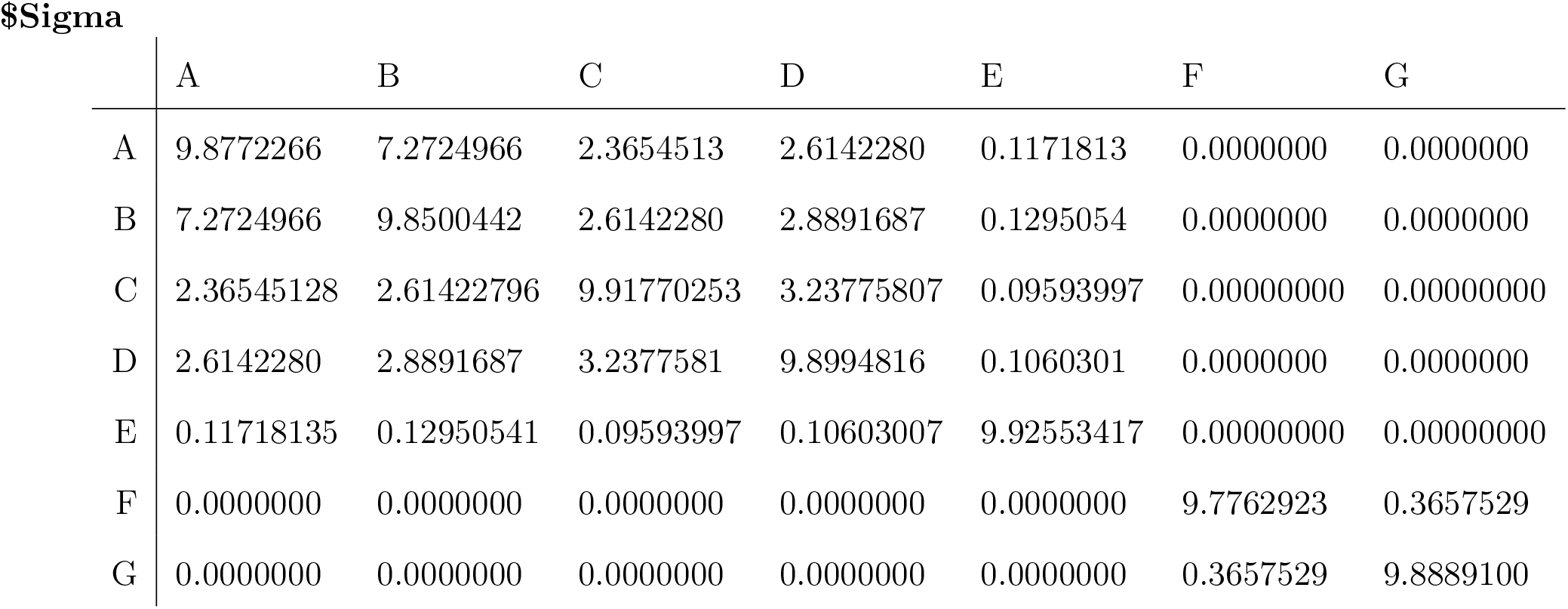

~~~
In [242]: getTipDistribution(modelOUbis, params, v = TRUE)
*** Computation of tip traits distribution through integrated formula ***
(Method working for models with a constant, A diagonalizable, and Gamma constant)
Computation time: 0.002770185 secs
Out[242]:
~~~

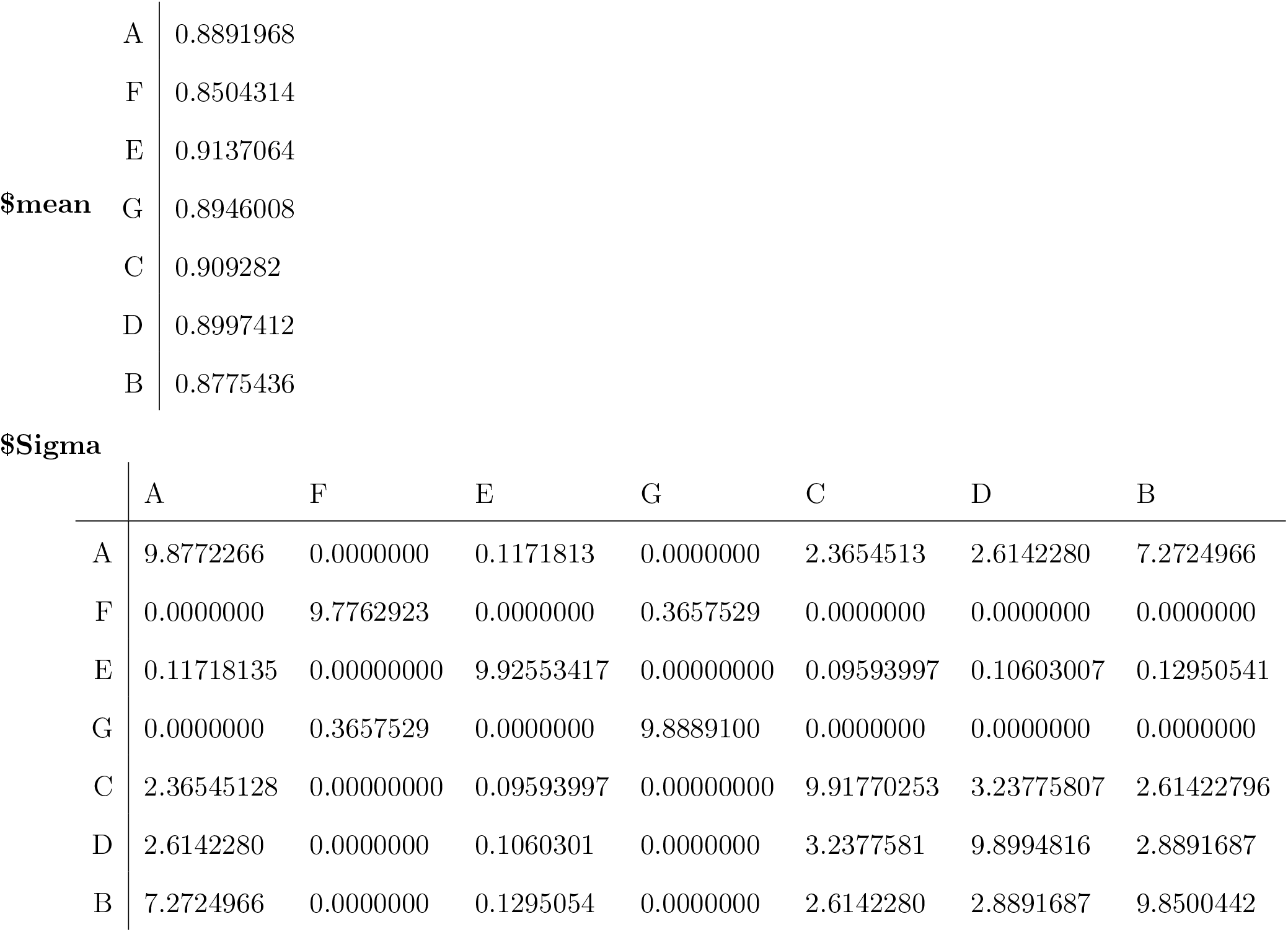

~~~
In [243]: getTipDistribution(modelOUter, params, v = TRUE)
*** Computation of tip traits distribution through ODE resolution ***
(Method working for any model)
Computation time: 0.01829243 secs
Out[243]:
~~~

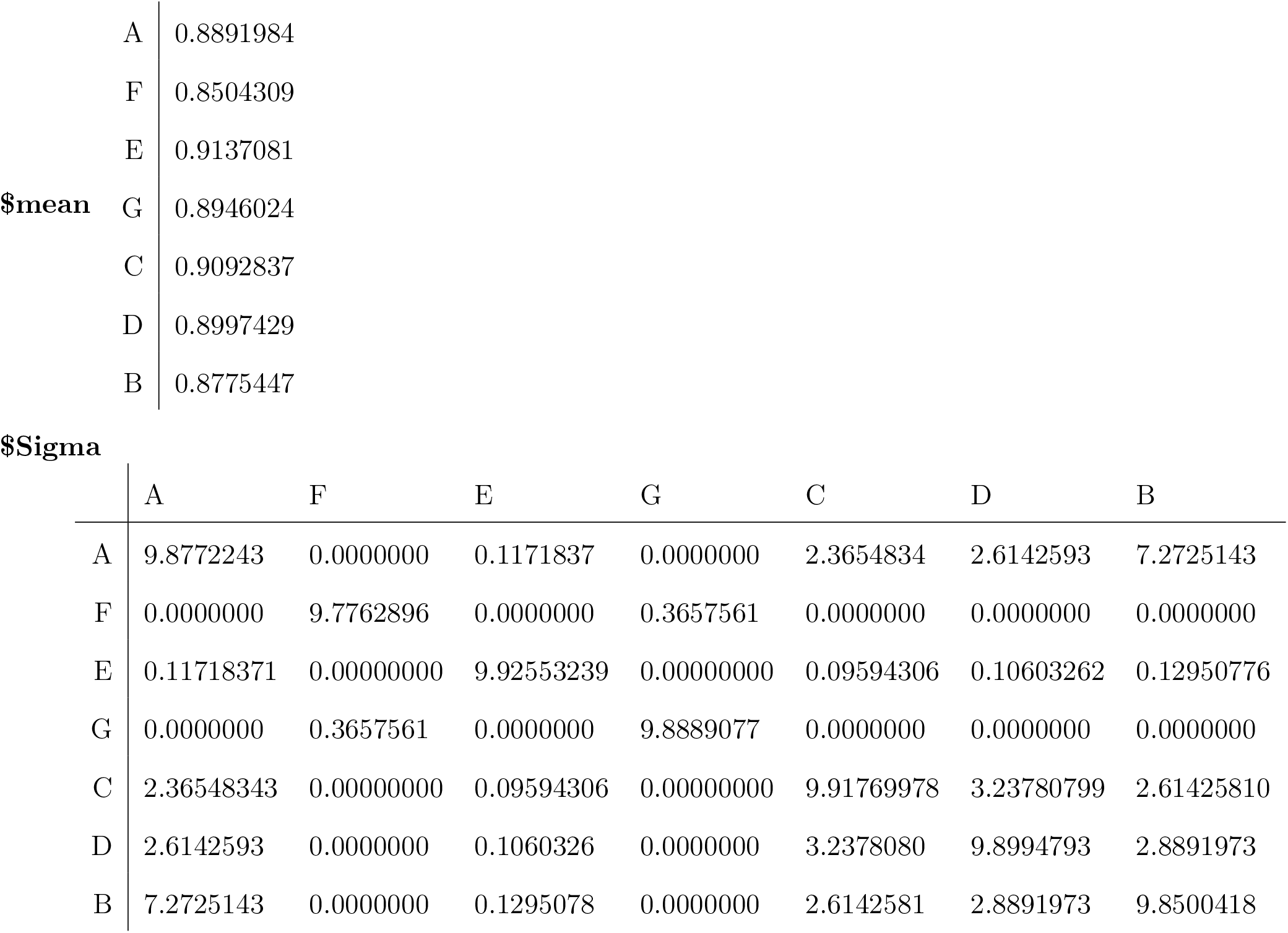

~~~
In [244]: dataOU <- simulateTipData(modelOU, c(0,0,0.2,1,2))
          fitTipData(modelOU, dataOU)
          fitTipData(modelOUbis, dataOU)
          fitTipData(modelOUter, dataOU)
*** Simulation of tip trait values ***
Simulates step-by-step the whole trajectory, but returns only the tip data.
Computation time: 0.2363398 secs
*** Fit of tip trait data ***
Finds the maximum likelihood estimators of the parameters,
returns the likelihood and the inferred parameters.
Computation time: 0.1814284 secs
Out[244]:
**$value** 15.0174906724384
**$inferredParams m0** -26.3722559360675
          **v0** 0.111663973605588
          **psi** 0.0973609295443122
          **theta** 14.9673044542728
          **sigma** 1.12338425846849
*** Fit of tip trait data ***
Finds the maximum likelihood estimators of the parameters,
returns the likelihood and the inferred parameters.
Computation time: 0.7557919 secs
Out[244]:
**$value** 15.0174906724384
**$inferredParams m0** -26.3722559360675
        **v0** 0.111663973605588
        **psi** 0.0973609295443122
        **theta** 14.9673044542728
        **sigma** 1.12338425846849
*** Fit of tip trait data ***
Finds the maximum likelihood estimators of the parameters, returns the likelihood and the inferred parameters.
Computation time: 6.088683 secs
Out[244]:
**$value** 15.0174914969285
**$inferredParams m0** -26.3722559360675
        **v0** 0.111663973605588
        **psi** 0.0973609295443122
        **theta** 14.9673044542728
        **sigma** 1.12338425846849
~~~

Focusing on the computation time, it is quite easily seen how interesting it can be to do some more analytical work and write more appropriated `getTipDistribution` functions. Still, the defaut function written for the mother class `PhenotypicModel` should always work.

##### Using the framework to define a new model.—

We illustrate here how the current code can be used to numerically study a specific model that has not been implemented elsewhere. We focus here on the implementation of the ‘GMM’ model described in the main text, explaining step by step the following procedure, that is generalizable to any model:

1. we identify what the periods are,
2. we write the model in a vectorial form on each period,
3. we implement it naively first,
4. we make analytical developments to speed up the computation time, and subsequently introduce a new class more appropriated to this model.

For simplicity, we implement GMM for two ultrametric trees here. In our example, the two trees will be:

~~~
In [245]: newick1 <- “(((A:1,B:1):3,(C:3,D:3):1):2,E:6);”
          tree1 <- read.tree(text = newick1)
          plot(tree1)
          newick2 <- “((X:1.5,Y:1.5):3,Z:4.5);”
          tree2 <- read.tree(text = newick2)
          plot(tree2)
~~~

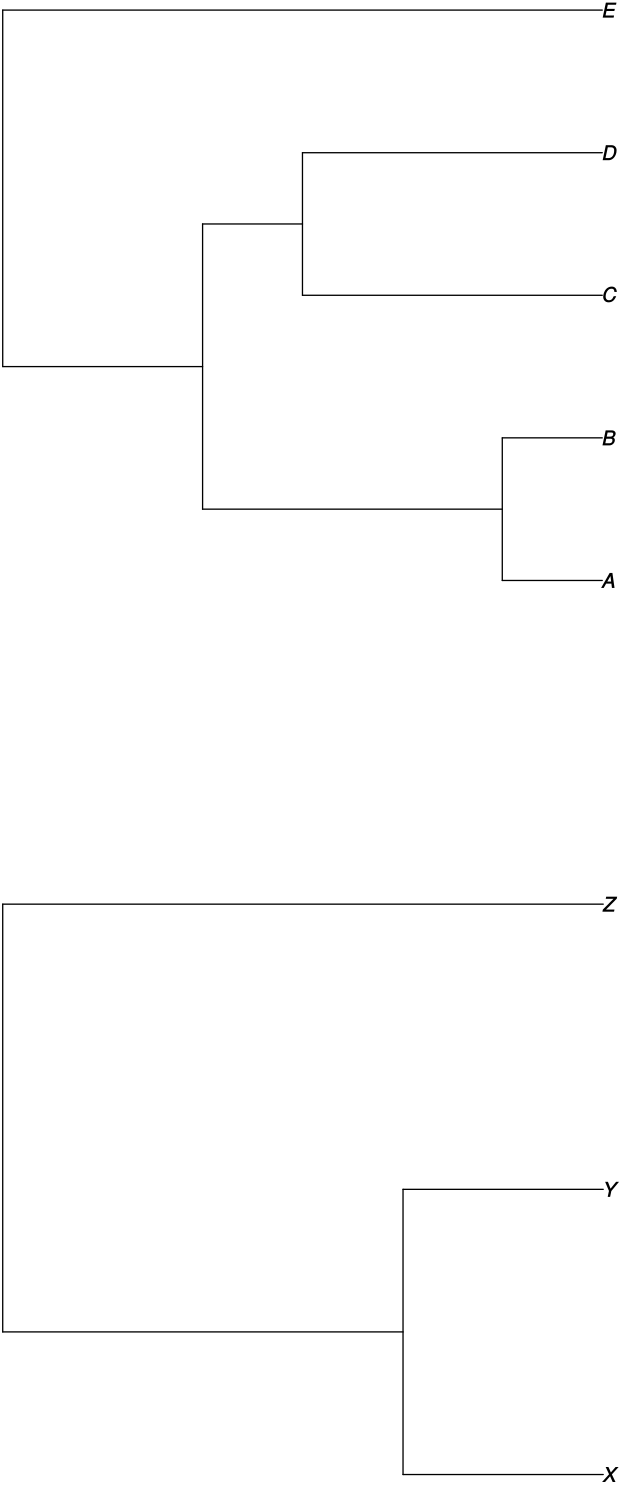

The first step consists in implementing a function `endOfPeriodsGMM(tree1, tree2)`, which takes as input two trees (the trees corresponding to our two interacting clades), and returns:

- the list of successive branching times (*τ_i_*) (vector `periods`),
- information on which branch gives birth at that time (vector `copy`),
- the number assigned to the newly created branch at that time (vector `paste`),
- the number of lineages in clade 1 and 2 at each time (vectors `nLineages1` and `nLineages2`),
- the label of tips at the end (vector `labeling`).

For example, our function, called on the two preceding trees, returns:

~~~
In [246]: endOfPeriodsGMM(tree1, tree2)
Out[246]:
**$periods** 0 1.5 2 3 4.5 5 6
**$copy** 1 3 1 3 5 1 0
**$paste** 2 4 3 4 7 5 0
**$nLineages1** 2 2 3 4 4 5 0
**$nLineages2** 1 2 2 2 3 3 0
**$labeling** ‘A’ ‘E’ ‘C’ ‘D’ ‘B’ ‘X’ ‘Z’ ‘Y’
~~~

The second step now consists in writing the model in the vectorial form required in the framework, on each epoch *i*. The form of the *a*, *A* and Γ matrices is shown in Appendix C.4, and depends on the number of lineages in the two clades on each epoch.

We introduce the constructor `createModelCoevolution(tree1, tree2)`, which is a function that takes as input two ultrametric trees corresponding to the two clades, and returns an object of class `PhenotypicModel`. It relies on the central function `aAGamma` that defines the collection of (*a_i_, A_i_*, Γ*_i_*) on each epoch.

This first version of the GMM implementation allows us to simulate tip data, to get the tip distribution under any parameter set, and to fit tip data.

~~~
In [248]: modelGMMbis <- createModelCoevolution(tree1, tree2, keyword=“GMMbis”)
          modelGMMbis
Out[248]:
****************************************************************
*** Object of Class PhenotypicModel ***
*** Name of the model: [1] “GMMbis”
*** Parameters of the model: [1] “m0” “v0” “d1” “d2” “S” “sigma”
*** Description: Generalist Matching Mutualism model.
Starts with 3 or 4 lineages having the same value X_0 ∼ Normal(m0,v0).
One trait in each lineage, all lineages evolving then non-independtly
according to the GMM expression.
*** Periods: the model is cut into 7 parts.
For more details on the model, call: print(PhenotypicModel)
****************************************************************
In [249]: dataGMM <- simulateTipData(modelGMMbis, c(0,0,5,-5, 1, 1), method = 2)
*** Simulation of tip trait values ***
Simulates step-by-step the whole trajectory, plots it, and returns tip data.
Computation time: 0.319762 secs
~~~

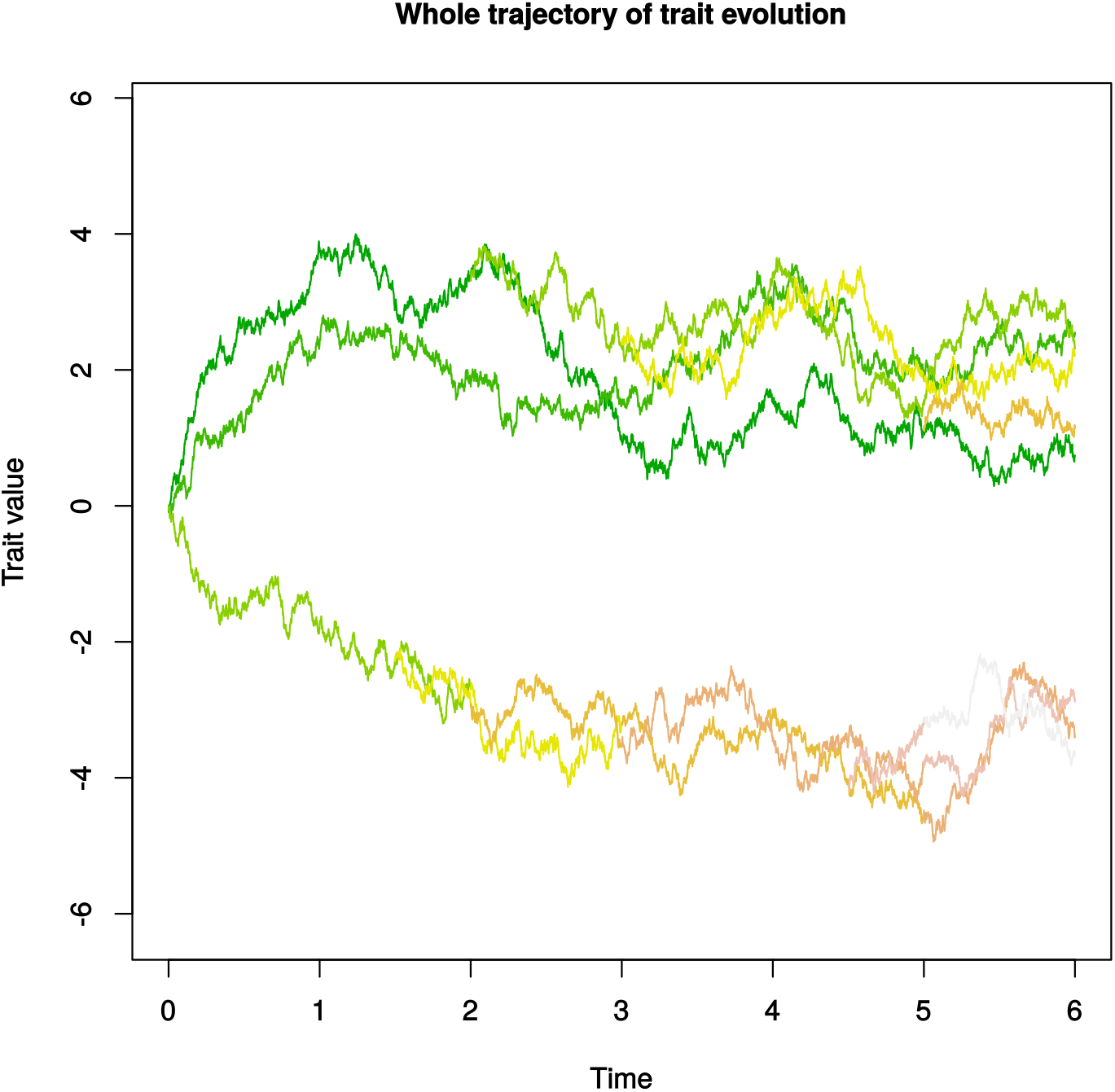

~~~
In [250]: getTipDistribution(modelGMMbis, c(0,0,5,-5,0.5,1))
Out[250]:
~~~

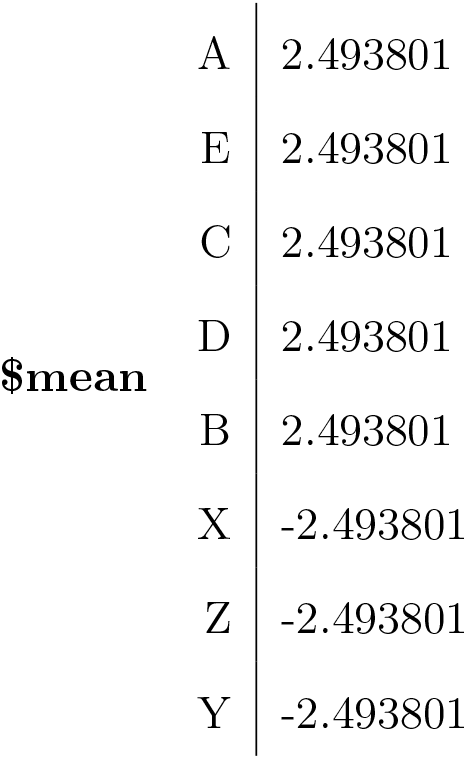

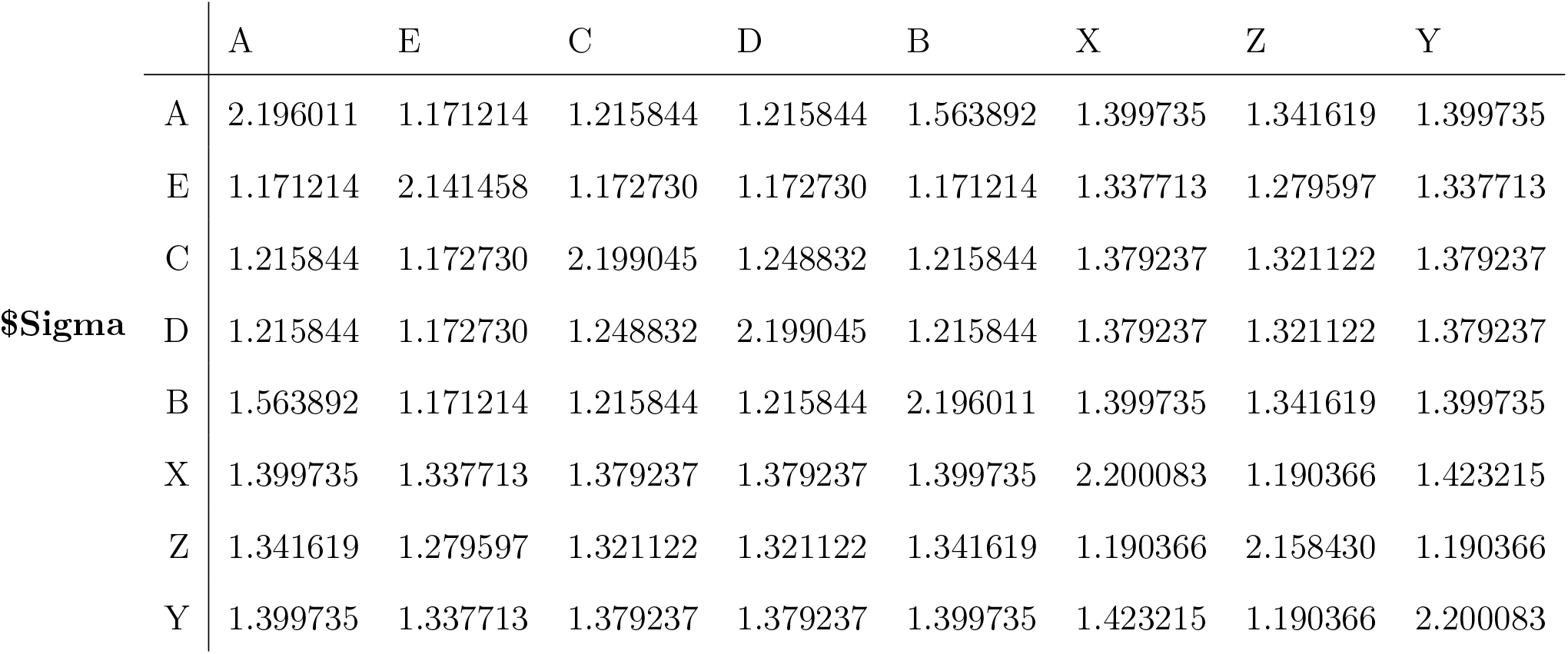

~~~
In [251]: fitTipData(modelGMMbis, dataGMM, c(0,0,5,-5,1,1))

*** Fit of tip trait data ***
Finds the maximum likelihood estimators of the parameters,
returns the likelihood and the inferred parameters.
Computation time: 3.728739 secs

Out[251]:

**$value** 6.61385667009296
**$inferredParams m0** 0.00512480151380221
   **v0** 2.69680996514239e-05
   **d1** 5.03536962882004
   **d2** -5.83517142115953
   **S** 0.231631941480316
   **sigma** 0.361942471141108
~~~

However, this first implementation relies on the `PhenotypicModel` class, which uses the method `getTipDistribution` that solves the ODE system on each epoch, and thus takes time.

The analytical reduction presented in Appendix C.4 can also be implemented. To this end, we create a new class named `PhenotypicGMM`, associated with an other function `getTipDistribution`. Using these developments allows us to compute more rapidly the tip distribution under the model.

~~~
In [252]: modelGMM <- createModelCoevolution(tree1, tree2, keyword="GMM”)
          modelGMM
Out[252]:
****************************************************************
*** Object of Class PhenotypicModel ***
*** Name of the model: [1] “GMM”
*** Parameters of the model: [1] “m0” “v0” “d1” “d2” “S” “sigma”
*** Description: Generalist Matching Mutualism model.
Starts with 3 or 4 lineages having the same value X_0 ∼ Normal(m0,v0). One trait in each lineage, all lineages evolving then non-independtly according to the GMM expression.
*** Periods: the model is cut into 7 parts.
For more details on the model, call: print(PhenotypicModel)
****************************************************************
In [253]: getTipDistribution(modelGMM, c(0,0,5,-5,0.5,1), v = TRUE)
          getTipDistribution(modelGMMbis, c(0,0,5,-5,0.5,1), v = TRUE)
*** Analytical computation of tip traits distribution ***
(Method working for the GMM model only)
Computation time: 0.0008528233 secs
Out[253]:
~~~

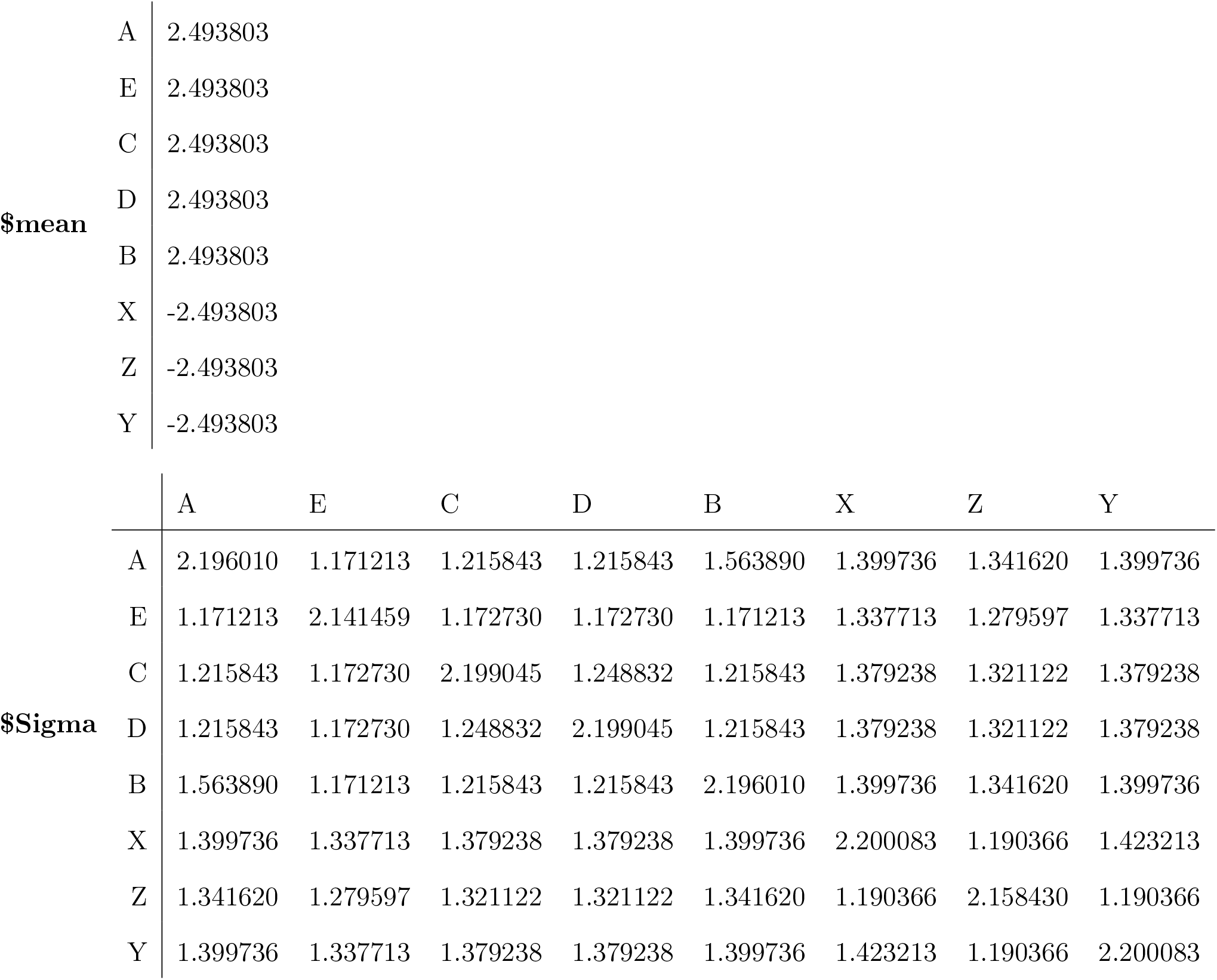

~~~
*** Computation of tip traits distribution through ODE resolution ***
(Method working for any model)
Computation time: 0.01734638 secs
Out[253]:
~~~

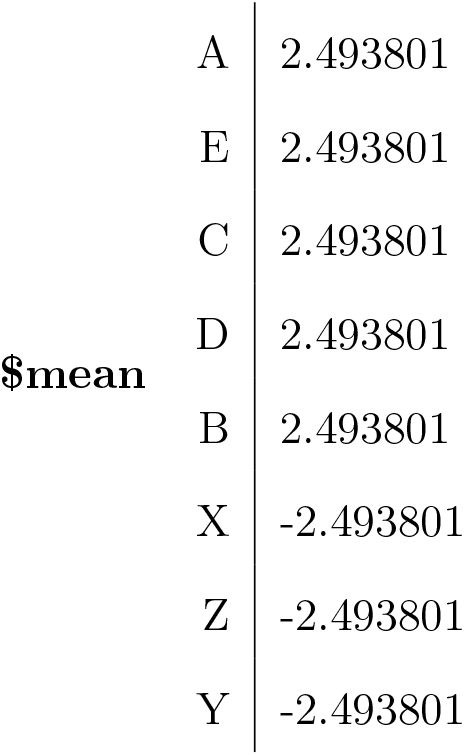

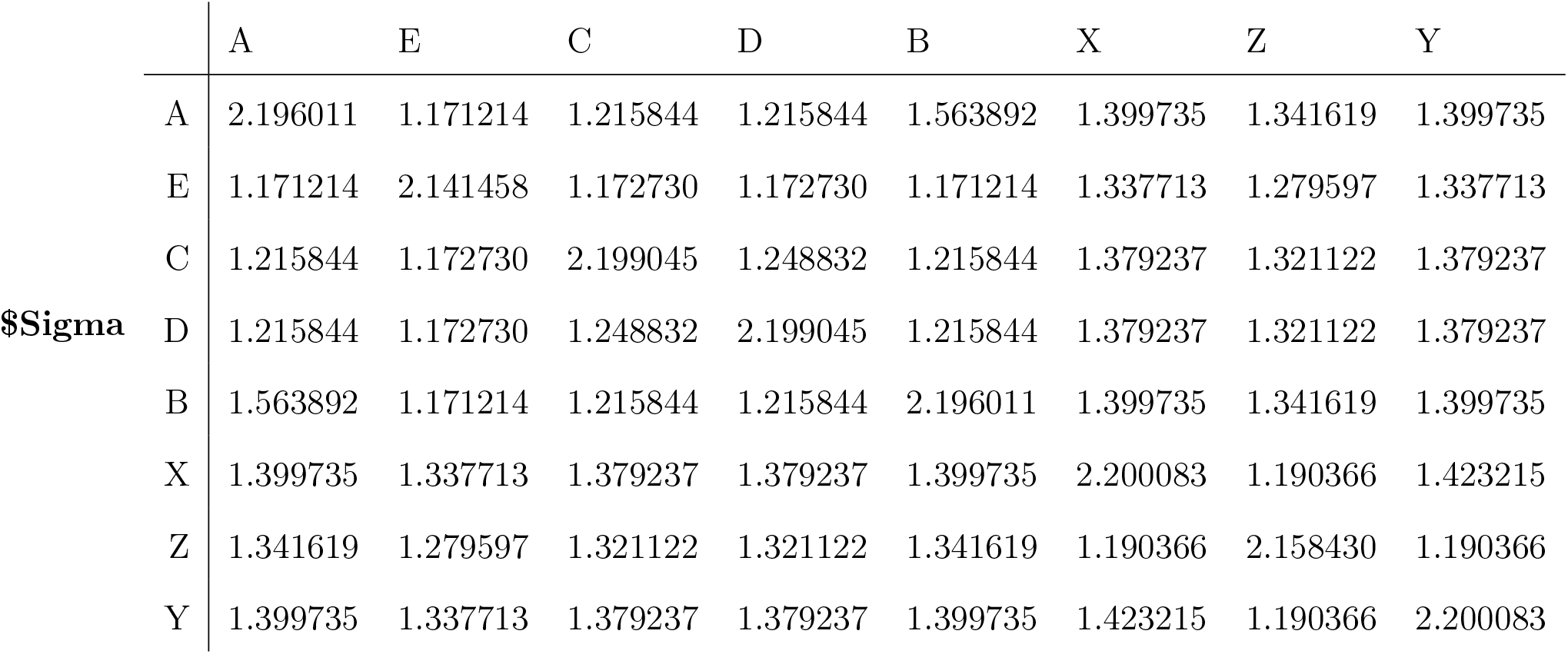

